# ELECTROPHYSIOLOGICAL CORRELATES OF CONSCIOUS EXPERIENCES DURING SLEEP: LUCID DREAMS, SLEEP PARALYSIS, OUT-OF-BODY EXPERIENCES, AND FALSE AWAKENINGS

**DOI:** 10.1101/2025.02.10.637510

**Authors:** Nerea L. Herrero, Yohann Corfdir, Aylin A. Vázquez-Chenlo, Lucila Capurro, Cecilia Forcato

**Author notes:** Corresponding author Laboratorio de Sueño y Memoria, Departamento de Ciencias de la Vida. Instituto Tecnológico de Buenos Aires (ITBA). Iguazú 341, (1437) Capital Federal, Buenos Aires, Argentina.

## Abstract

Consciousness does not always fade during sleep. Instead, it can re-emerge, giving rise to lucid dreams (LDs), sleep paralysis (SP), out-of-body experiences (OBEs), and false awakenings (FAs). While some of these states have been studied phenomenologically, their neurophysiological underpinnings remain unclear.

Here, we investigate their electrophysiological correlates and distinguish them from standard sleep stages. We conducted overnight polysomnography in frequent experiencers, capturing 10 episodes (3 LDs, 2 SP, 2 OBEs, 3 FAs). Eye movement markers identified periods of lucidity. Relative spectral power was analyzed using principal component analysis (PCA) and permutation-based multivariate analysis of variance (PERMANOVA).

Our results indicate that these conscious sleep states are distinct from wakefulness, yet share features with both stage 1 (S1) and rapid eye movement (REM) sleep. Notably, we provide the first documented eye movement markers during FAs and OBEs.

## Introduction

During wakefulness, all our experiences, thoughts, emotions, and feelings combine and unfold as a whole, resulting in the unique and irreplaceable sensation of having a subjective experience and thus being individuals endowed with consciousness. Consciousness remains stable during wakefulness, and while it may be absent during certain moments of sleep, it can also emerge under special conditions such as lucid dreams (LDs), sleep paralysis (SP), out-of-body experiences (OBEs) and false awakenings (FAs).

LDs are dreams in which individuals become aware that they are dreaming. During a LD, people can act voluntarily within the dream and are often able to recall and execute pre-planned actions that were agreed upon before sleep^1–3^. LDs are primarily a REM sleep phenomenon, with few exceptions^4^. Lucid dreamers can signal the onset of their lucid state through pre-arranged eye movements, enabling communication with researchers in laboratory settings. Electroencephalographic (EEG) studies have identified distinct brain activity associated with lucid dreaming, such as an increase in low-beta frequencies (13-19 Hz) in parietal regions^2^ and an increase in frontolateral low-gamma activity (40 Hz)^5,6^. However, recent studies have questioned the previously reported low-gamma increase during this state. For instance, Baird et al. (2022) argued that this phenomenon might be an artifact caused by saccadic spike potentials, which are linked to the heightened REM density observed during LDs. Conversely, a more recent study reported augmented low-gamma activity during LDs, leaving the role of low-gamma inconclusive^7^.

SP is a parasomnia characterized by the temporary inability to perform voluntary movements during the transitions between sleep and wakefulness^8,9^. During an SP episode, individuals often perceive themselves as being awake and aware of their surroundings, despite still being in a sleep state. SP can cause significant distress due to the presence of negative emotions and vivid hallucinations that commonly accompany episodes^10,11^. EEG studies on episodes of SP are scarce and have been conducted in both healthy individuals^12–14^ and patients with narcolepsy^12,15,16^. Most of these studies performed qualitative analyses of the EEG signal, highlighting the presence of “alpha trains” during SP^13–15^. More recent findings, however, have revealed a significant increase in relative alpha power during SP^12^ accompanied by a significant decrease in relative theta power compared to REM sleep (Mainieri et al., 2021). Additionally, Terzaghi et al. (2012) described the coexistence of alpha-like rhythms and high-frequency peaks (∼18 Hz) during SP^16^.

OBEs are an altered state of consciousness in which individuals perceive themselves as being outside their physical body, observing the world from an external perspective^17,18^. OBEs have been reported across various cultures throughout history and remain one of the great mysteries in the study of human consciousness. They are often described as extremely vivid, with perceptual qualities similar to veridical perception^11,17^, and are associated with lasting psychological and existential changes, including increased spirituality, reduced fear of death, heightened empathy, and an expanded perception of reality^19–21^. Neuroscientific research suggests that OBEs may result from a temporary disruption in the integration of multisensory information, with the temporo-parietal junction (TPJ) playing a critical role in this process^18,22,23^. OBEs associated with sleep are often linked to other phenomena, such as lucid dreaming, sleep paralysis, and false awakenings^11,24,25^. To date, no studies have successfully captured OBEs occurring during sleep with EEG or other neurophysiological measures.

FAs are dreams in which the subjects have an erroneous belief that they are waking up in a familiar place, starting a daytime routine, only to later realize that they are still dreaming^26^. There are only two EEG studies of FA published^12,13^. Of these, Takeuchi et al. (1992) did not formally distinguish FAs from other episodes and included them within the broader category of SP in their analyses. In contrast, Mainieri et al. (2021) conducted a detailed examination of FAs, finding that their EEG profile closely resembles SP, characterized by a significant reduction in theta relative power and a notable increase in alpha relative power compared to REM sleep.

Here, we aimed to characterize the electrophysiological correlates of different conscious experiences occurring during sleep. Additionally, we sought to differentiate these states from standard sleep stages, such as REM sleep and S1, as well as wakefulness.

## Results

To characterize LDs, SP, OBEs, and FAs, we conducted polysomnography during full-night sleep sessions in the laboratory with individuals who frequently experience these states (3 LDs, 2 SP, 2 OBEs and 3FAs). Participants were trained to leave an eye mark whenever they became aware while sleeping. We first applied independent component analysis (ICA) to remove eye movement artifacts. Subsequently, we performed principal component analysis (PCA), and permutation-based multivariate analyses of variance (PERMANOVA) for each subject, to identify key patterns in spectral power distributions across different brain regions and sleep stages (Fig. 1). This approach provided insights into how neural activity correlates with each state of consciousness.

**Fig. 1.**
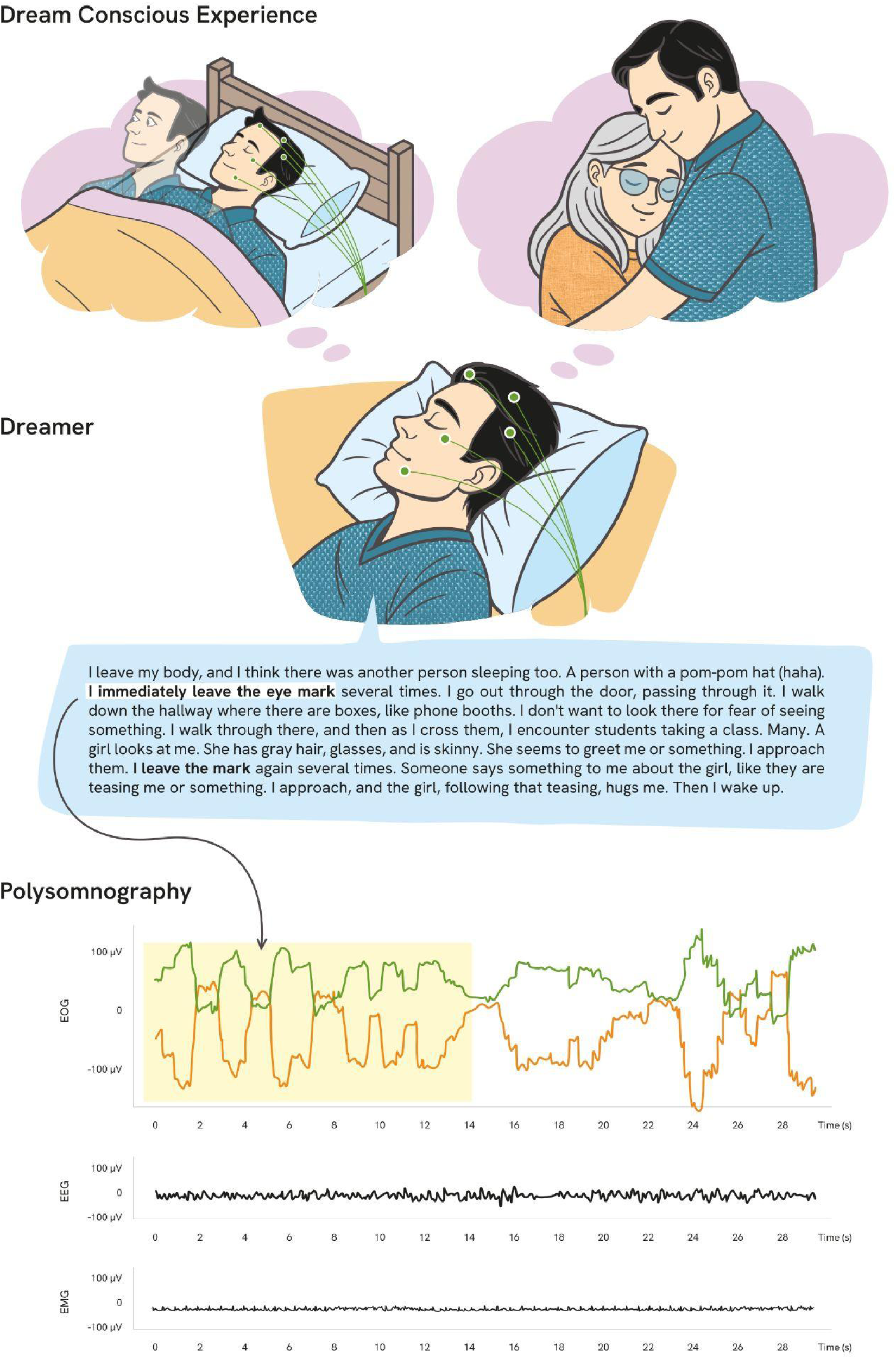
Representation of one of the dream conscious experiences of the study. Subject’s report detailing the experience and polysomnographic recording showing the eye marks and real-time physiological activity during the experience.

### Lucid Dreaming

First, the PCA revealed that LDs were clearly distinguishable from wakefulness in all three subjects (Fig. 2). However, they shared characteristics with both S1 and REM sleep, except in subject 2, where LD was also distinct from S1. Specifically, all LDs exhibited a stronger influence of delta and theta waves similar to REM (LD_1_: Dimension 1, cos² = 0.440, contribution = 10.802%, Dimension 2, cos^2^ = 0.150, contribution = 13.221%; LD_2_: Dimension 1, contribution = 12.854%, cos² = 0.443, Dimension 2, cos^2^ = 0.118, contribution = 11.181%, LD_3:_ Dimension 1, contribution = 12.944%, cos² = 0.242, Dimension 2, contribution = 38.610%, cos² = 0.212). In contrast, wakefulness exhibited a stronger influence of alpha activity. Additionally, during LDs, higher-frequency bands such as beta and low-gamma showed an influence, particularly in subjects 1 and 3, resembling patterns observed in S1. See supplementary Tables 1-15 and supplementary Figs. 1-10 for the complete analysis.

**Fig. 2.**
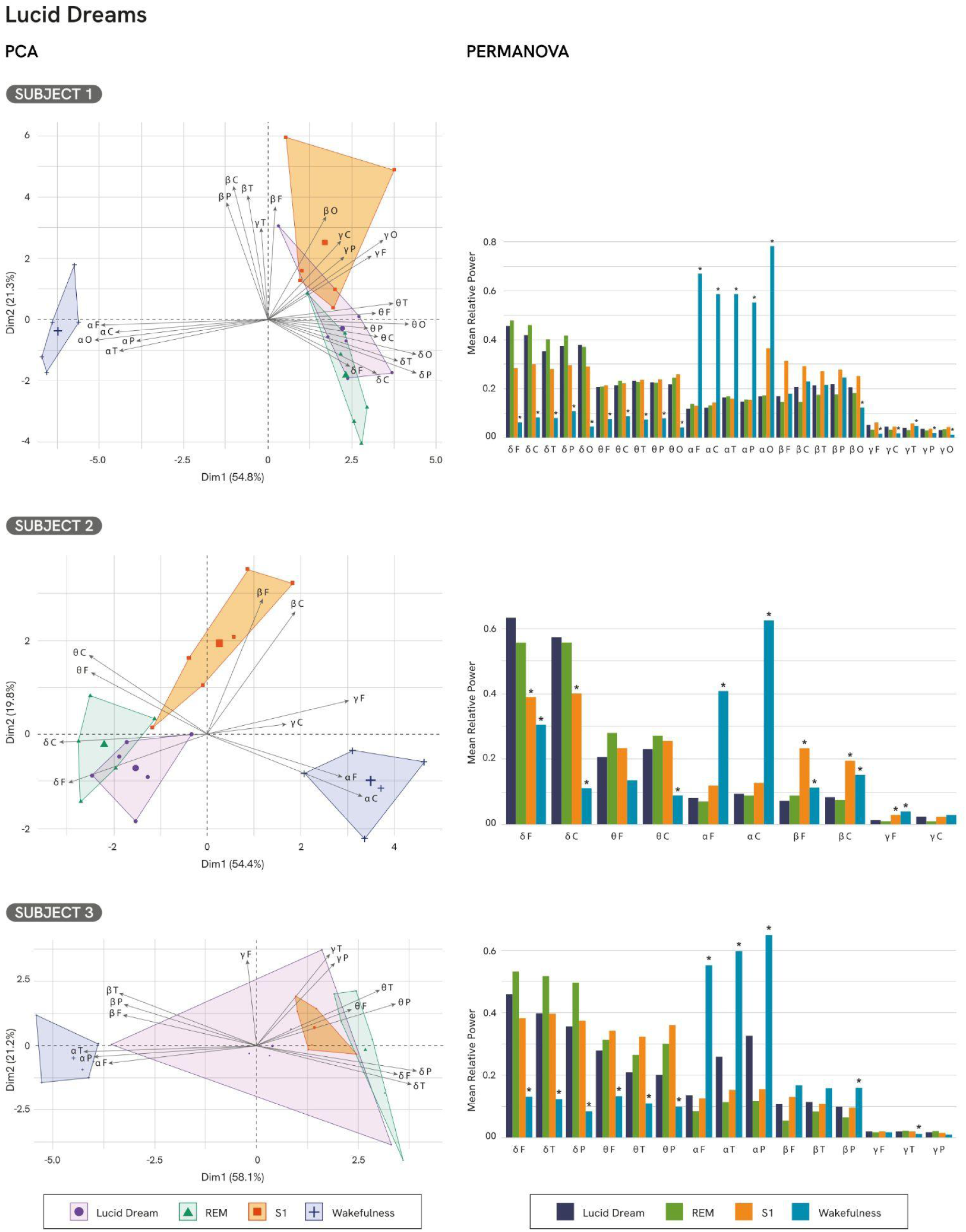
Principal Component Analysis (PCA) and PERMANOVA results for lucid dreams Subjects 1, 2, and 3. The PCA biplot illustrates the distribution of lucid dreams (LD), wakefulness, REM sleep, and S1 across the principal components. The percentage of variance explained by each dimension (Dim) is indicated on the axes. For PERMANOVA, all significant differences (p < 0.05) are marked with an asterisk (*), indicating comparisons against the respective condition. Abbreviations: F = frontal, C = central, T = temporal, P = parietal, O = occipital.

Additionally, the PERMANOVA revealed significant differences in oscillatory activity across the states of consciousness (Subj_1_: F(3,20) = 36.499, p < 0.001, R² = 0.845; Subj_2_: F(3, 20) = 24.705, p < 0.001, R² = 0.788; Subj_3_: F(3, 20) = 15,831, p < 0.001, R**²** = 0,704). Specifically, all LDs differed significantly from Wakefulness (PERMANOVA, Subj_1_: F(1,10) = 72.452, p = 0.011, R² = 0.879, Subj_2_: F(1,10) = 46.144, p = 0.013, R² = 0.822, Subj_3_: F(1,10) = 15.275, p = 0.013, R² = 0.604).

Furthermore, the analysis across regions revealed that all LDs exhibited significant increases in slow-frequency bands (delta and theta) and decreases in fast-frequency bands (alpha, beta and low-gamma) compared to wakefulness. Notably, subject 1 showed a decrease in low-gamma activity in the temporal region alongside increases in frontal, central, parietal, and occipital low-gamma compared to wakefulness.

No significant differences were observed between S1 and LD periods for subjects 1 and 3 (PERMANOVA, Subj_1_: F(1,10)= 3.244, p = 0.217, R² = 0.245; Subj_3_: F(1,10) = 2.303 p = 0.500, R² = 0.187). However, LDs significantly differed from S1 in subject 2 (PERMANOVA, Subj_2_: F(1,10) = 12.345, R² = 0.552, p = 0.017), with an increase in fronto-central delta activity during LDs.

Finally, no significant differences were found between LD and REM sleep (Fig. 2, PERMANOVA, Subj_1_: F(1,10) = 0.563, p = 1.000, R² = 0.054; Subj_2_: F(1,10) = 1.597, R² = 0.14, p = 1.000; Subj_3_: F(1,10) = 2.534, p = 0.522, R² = 0.202). See supplementary Tables 16-25 for complete analyses.

### Sleep Paralysis

PCA revealed distinct patterns of frequency band influence in the two episodes of SP. In subject 4, a greater influence of higher-frequency bands, such as beta and low-gamma was observed, accompanied by a less pronounced contribution from delta and theta bands, distinguishing this episode from REM (Subj_4_: Dimension 1, contribution = 8.355, cos^2^ = 0.141; Dimension 2, contribution = 68.268, cos^2^ = 0.420). Conversely, subject 5 displayed a prominent influence of alpha and beta bands, reflecting dominance of higher-frequency activity (Subj_5_: Dimension 1, contribution = 4.781, cos² = 0.184; Dimension 2, contribution = 16.209, cos² = 0.239) (Fig. 3). This pattern highlights clear distinctions from REM sleep . See supplementary Tables 26-35 and supplementary Figs. 11-20 for the complete analysis.

Furthermore, there was a significant difference in the oscillatory activity among the states of consciousness (PERMANOVA, Subj_4_: F(3, 20) = 18.832, p < 0.001, R**²** = 0.738, Subj_5_: F(3, 20) = 17.174, p < 0.001, R**²** = 0.720).

Compared to REM, both episodes showed significant differences (PERMANOVA, Subj_4_: F(1,10) = 5.367, p=0.014, R² = 0.361; Subj_5_: F(1,10) = 17.87, p = 0.016, R² = 0.64). In particular, both episodes showed significant decreases in theta activity. Subject 4 exhibited reduced theta activity across all regions, while subject 5 showed localized reductions in temporal, parietal, and occipital regions. Delta activity yielded mixed results, as subject 4 exhibited a significant increase in the central region, while subject 5 showed reductions across all regions, rendering the role of delta activity inconclusive. Both episodes showed increased low-gamma activity, with subject 4, displaying increases in frontal, temporal, and occipital regions, and subject 5 in frontal, central, temporal, and parietal regions. Subject 5 also demonstrated significant increases in alpha and beta activity across all regions.

Compared to S1, both SP exhibited significant differences (PERMANOVA, Subj_4_: F(1,10) 5.462, p = 0.042, R² = 0.353; Subj_5_: F(1,10)= 5.895, p = 0.031, R² = 0.370). Theta power was significantly reduced in both episodes. Specifically, subject 4 exhibited reductions in the frontal, central, temporal, and occipital regions and subject 5 showed decreases across all regions. Once again, the findings regarding delta activity remains inconclusive. Specifically, subject 4 exhibited a significant decrease in parietal delta accompanied by an increase in central delta. Conversely, subject 5 exhibited a significant decrease in central delta, further highlighting the inconclusive role of delta activity in SP. Concerning fast-frequency bands, both SP episodes showed a significant increase in low-gamma activity. Subject 4 exhibited increased low-gamma relative power across frontal, temporal, parietal, and occipital regions, while subject 5 demonstrated a localized increase in centro-parietal regions. Additionally, both subjects 4 and 5 showed increases in beta power, with subject 4 displaying elevated beta activity in the centro-parietal region, and subject 5 showing increased beta power across all areas. Subject 4 also showed increased alpha activity in fronto-parietal areas, a pattern not observed in subject 5.

Compared to wakefulness, both episodes showed significant differences (PERMANOVA, Subj_4_: F(1,10) = 21.756, p = 0.016, R² = 0.685; Subj_5_: F(1,10)= 11.123, p = 0.013, R² = 0.526). Slow-frequency bands (delta and theta) increased significantly, in both subjects displaying higher delta relative power in the central, temporal, parietal, and occipital regions. Theta activity increased in parieto-occipital regions for subject 4, and across all regions for subject 5. Furthermore, both SP episodes exhibited significant reductions in the fast-frequency bands (alpha and beta activity). Specifically, subject 5 showed decreased alpha in the parietal region, while subject 4 displayed reduced alpha across all regions. Regarding beta activity, subject 4 showed a decrease in the central region, whereas subject 5 showed a decrease in the temporal region. low-gamma activity was reduced in subject 5 across all regions, while subject 4 showed increased low-gamma activity in the parieto-occipital regions. See supplementary Tables 36-46 for complete analyses.

**Fig. 3.**
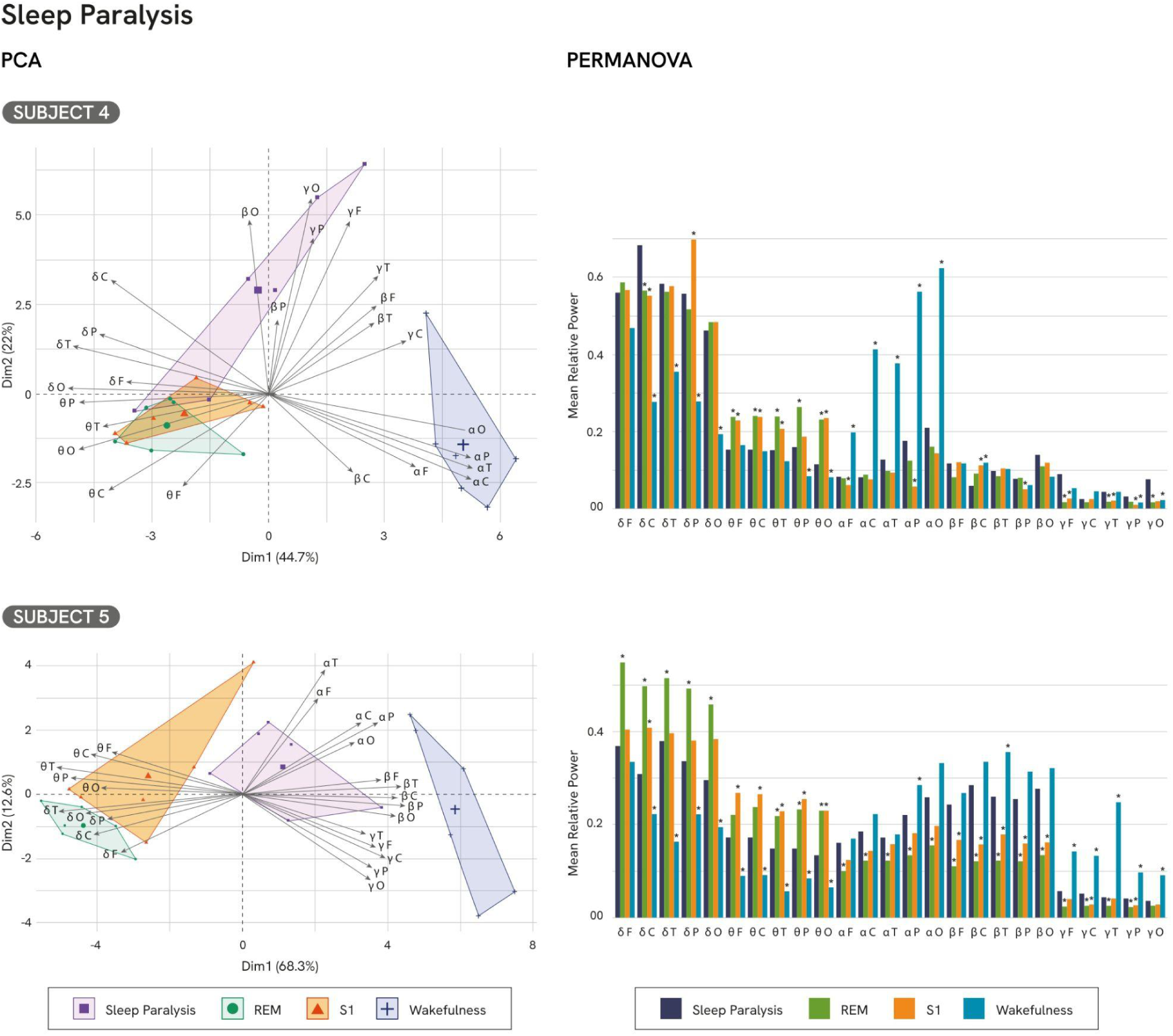
Principal Component Analysis (PCA) and PERMANOVA results for sleep paralysis (Subjects 4 and 5). The PCA biplot illustrates the distribution of sleep paralysis (SP), wakefulness, REM sleep, and S1 sleep across the principal components. The PERMANOVA results indicate significant differences in mean relative power between subjects, with post-hoc analyses revealing the specific frequency bands driving these differences. All significant differences (p < 0.05) are marked with an asterisk (*), including highly significant values.

### Out-of-body Experiences

The PCA revealed that both experiences (OBE_1_ and OBE_2_) of subject 3, were primarily characterized by delta and theta frequency bands (OBE_1_: Dimension 1, contribution = 20.363%, cos^2^ = 0.615, Dimension 2, contribution = 16.331%, cos^2^ = 0.126; OBE_2_: Dimension 1, contribution = 15.820%, cos^2^ = 0.398, Dimension 2, contribution = 15.265%, cos^2^ = 0.353) (Fig. 3). Both episodes differed from wakefulness and S1 while sharing similarities with REM, predominantly due to delta and theta bands. However, OBE_1_ showed additional differences from REM, including stronger influence from slow-frequency bands and a reduction of fast-frequency bands. See supplementary Tables 47-56 and supplementary Figs. 21-29 for the complete analysis.

PERMANOVA confirmed a significant effect of the state of consciousness on oscillatory activity for both, OBE_1_ (F(3, 20) = 53.134, p < 0.001, R² = 0.889) and OBE_2_ (F(3, 20) = 15.831, p < 0.001, R² = 0.704). Specifically, we found significant differences between OBE_1_ and REM (PERMANOVA, F(1,10) = 6.075, p = 0.049, R² = 0.378), but no for OBE_2_ (PERMONOVA, F(1,10) = 1.524, p = 1.000, R² = 0.132).

OBE_1_ exhibited significant increases in delta relative power in frontal, temporal, and parietal regions and reductions in the relative power of high-frequency bands, including alpha in frontal, central, temporal, and parietal regions, beta in frontal, central, and temporal regions and temporal low-gamma .

Both OBEs showed significant differences compared to S1 (PERMANOVA, OBE_1_: F(1,10) = 11.652, p = 0.029, R² = 0.538; OBE_2_: F(1,10) = 11.616, p = 0.017, R² = 0.537). Both episodes showed a significant increase in delta . OBE_1_ showed a significant reduction in fronto-temporal theta power along with decreases in beta and low-gamma across multiple regions).

There was a significant difference between wakefulness and both OBEs (PERMANOVA, OBE_1_ F(1,10) = 178.139, p = 0.010, R² = 0.947; OBE_2_ F(1,10)= 74.655, p = 0.013, R² = 0.882). Particularly, both OBEs showed significant increases in low-frequency bands, including delta and theta power, across all regions . In addition, significant decreases were observed in high-frequency bands, including alpha, beta, and low-gamma relative power, with region-specific patterns over OBEs. For alpha activity, both OBEs showed reductions across all regions. For beta relative power, OBE_1_ demonstrated significant decreases in frontal, central, temporal, and parietal regions. OBE_2_, on the other hand, showed reductions restricted to fronto-central regions. Finally, for low-gamma relative power, OBE_1_ exhibited significant reductions in frontal, central, and temporal regions whereas OBE_2_ showed reductions in parieto-occipital regions). See supplementary Tables 57-66 for complete analyses.

**Fig. 3.**
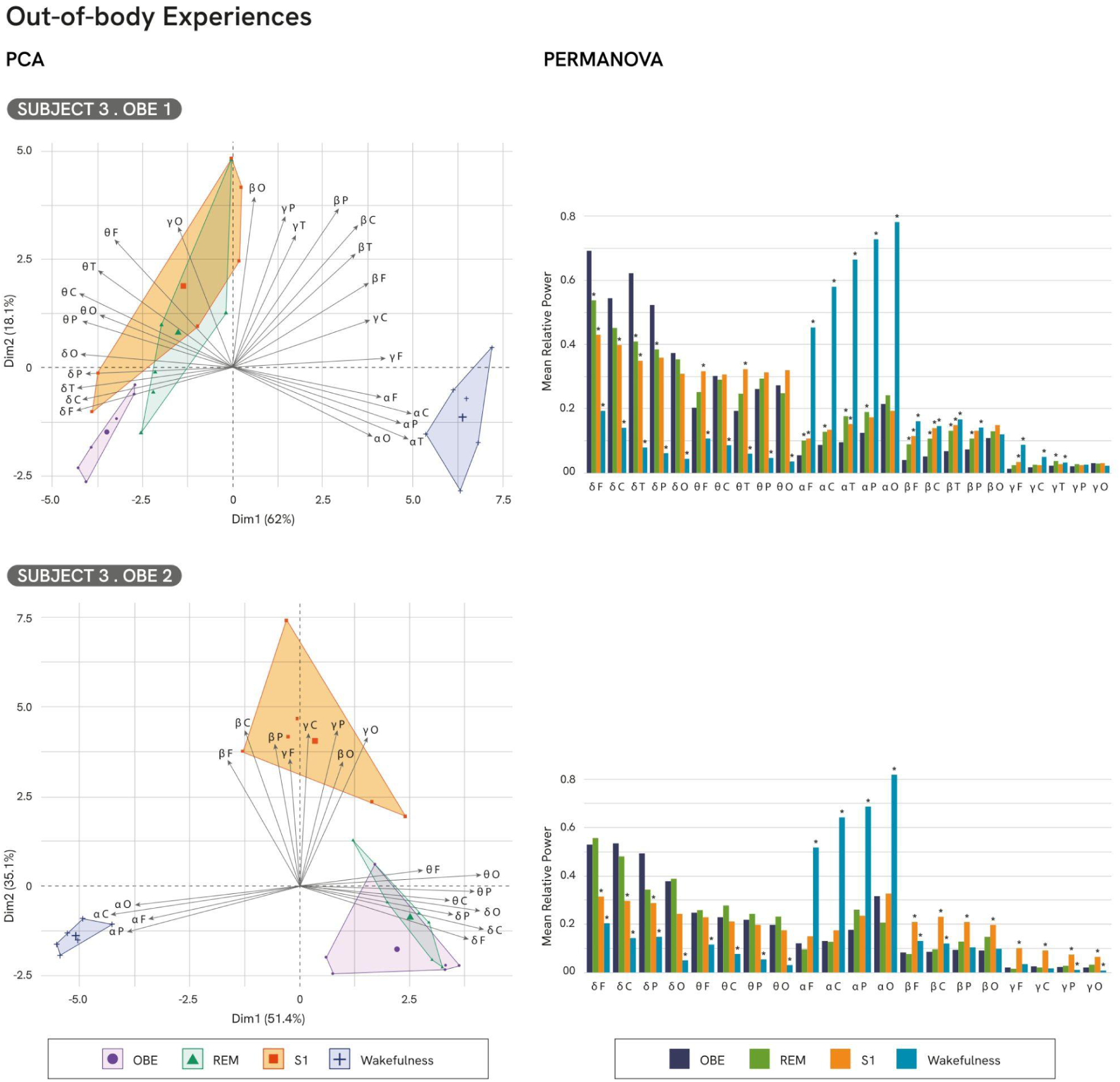
Principal Component Analysis (PCA) and PERMANOVA results for OBE from subject 3. The PCA biplot illustrates the distribution of out-of-body experience (OBE), wakefulness, REM sleep, and S1 sleep across the principal components. The PERMANOVA results indicate significant differences in mean relative power between conditions, with post-hoc analyses revealing the specific frequency bands driving these differences. All significant differences (p < 0.05) are marked with an asterisk (*), including highly significant values.

### False Awakenings

The PCA for the three FA cases revealed the influence of theta activity along distinct frequency bands (Fig. 4). FA of subject 3 was characterized by a notable influence of fast-frequency bands, including beta and low-gamma (mostly parietal and occipital), along with theta activity (Subj_3_: Dimension 1, contribution = 11.971%, cos^2^ = 0.235; Dimension 2, contribution = 35.685%, cos^2^ = 0.322). Subject 6 showed influence of alpha activity along with smaller contributions from beta, low-gamma, theta, and delta (Subj_6_: Dimension 1, contribution = 10.700%, cos^2^ = 0.244; Dimension 2, contribution = 63.827%, cos^2^ = 0.579). Subject 7 exhibited influence from alpha, delta and theta frequency bands (Subj_7_: Dimension 1, contribution = 12.59%, cos^2^ = 0.224; Dimension 2, contribution = 16.37%, cos^2^ = 0.578). Compared to REM, only subject 3 exhibited a distinctly different frequency profile, characterized by a pronounced influence of fast-frequency bands absent in REM. Subject 6 and subject 7 displayed profiles more similar to REM, with subtle variations such as reduced delta activity in subject 6, and a slight influence of alpha in subject 7. See supplementary Tables 67-81 and Figs. 30-40 for complete analyses.

**Fig. 4.**
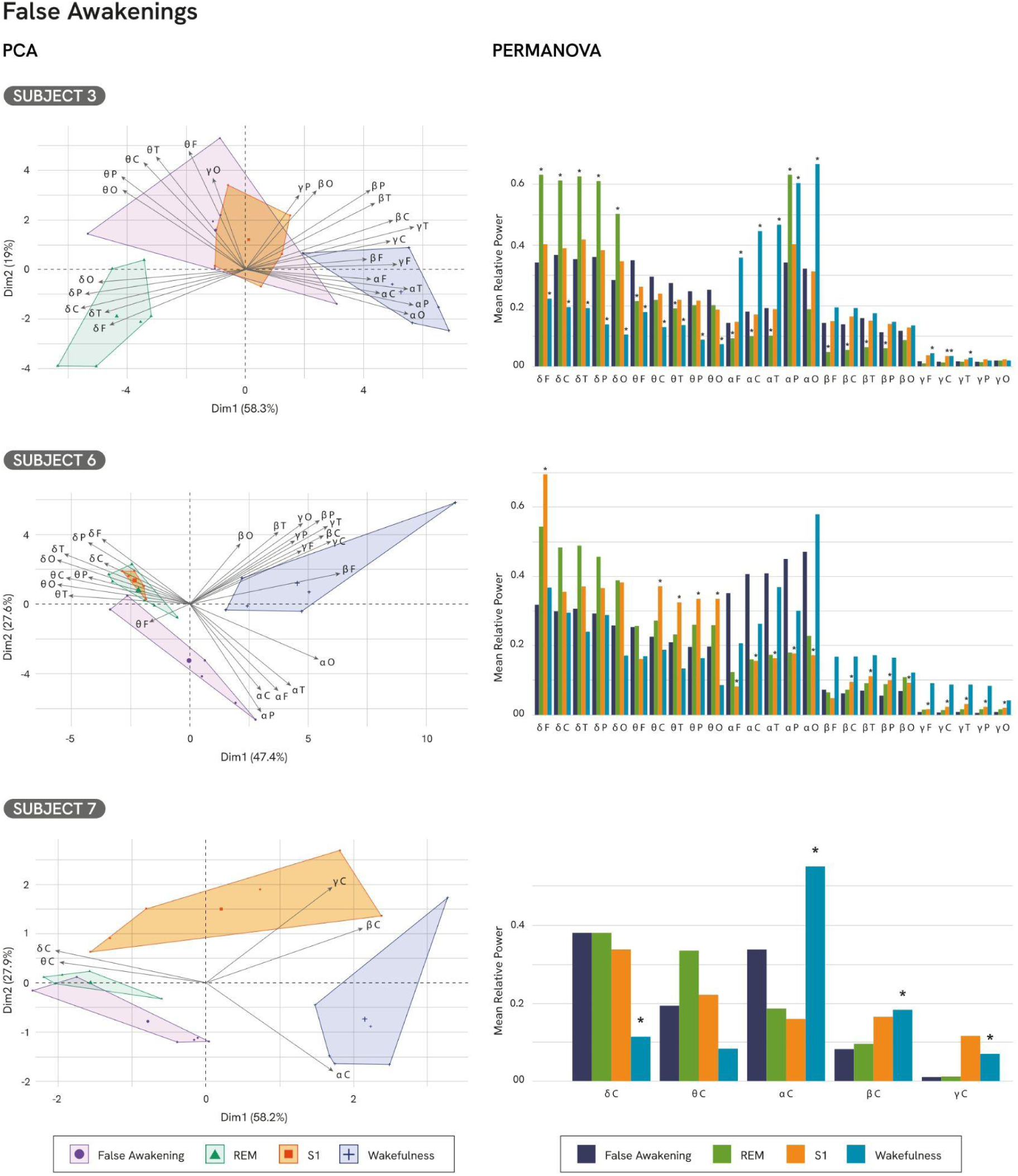
Principal Component Analysis (PCA) and PERMANOVA results for False Awakenings from subject 6 and 7. The PCA biplot illustrates the distribution of False Awakening (FA), wakefulness, REM sleep, and S1 sleep across the principal components. The PERMANOVA results indicate significant differences in mean relative power between conditions, with post-hoc analyses revealing the specific frequency bands driving these differences. All significant differences (p < 0.05) are marked with an asterisk (*), including highly significant values.

In comparison to S1, subject 6 and subject 7 exhibited differences, while subject 3 remained highly similar. Specifically, subject 6 showed a greater influence of alpha compared to S1, along with reduced influence from delta and theta, which were more pronounced in S1 but less evident in subject 6. Additionally, subject 7 displayed a reduced contribution from the fast-frequency bands beta and low-gamma relative to S1. All three FA cases showed clear differences when compared to wakefulness.

PERMANOVA revealed a significant effect of the state of consciousness on oscillatory activity (Subj_3_: F(3, 20) = 17.978, p < 0.001, R**²** = 0.729; Subj_6_: F(3, 20) = 6.19, p < 0.001, R**²** = 0.481; Subj_7_: F(3, 20) = 10.374, p < 0.001, R**²** = 0.609).

Post-hoc analyses demonstrated a significant difference between subject 3 and REM (PERMANOVA, Subj_3_: F(1,10) = 9.252, p = 0.031, R² = 0.481) while non significant differences were observed for subject 6 or subject 7 (PERMANOVA, Subj_6_: F(1,10) = 5.718, p = 0.227, R² = 0.364; Subj_7_: F(1,10) = 3.282, p = 0.344, R² = 0.247). Subject 3 showed a significant reduction in delta relative power across all regions and an increase in fronto-temporal theta activity. Additionally, subject 3 exhibited increases in alpha across frontal, central, and temporal areas, and in beta activity in the frontal, central, temporal, and parietal regions.

Significant differences were also observed between subject 6 and S1 (PERMANOVA, F(1,10) = 9.575, p = 0.036, R² = 0.489). Specifically, subject 6 exhibited a significant reduction in frontal delta and theta across central, temporal, parietal, and occipital regions . Beta relative power decreased across central, temporal, parietal, and occipital regions, accompanied by reduction in low-gamma activity across all regions. Subject 6 also exhibited an increase in alpha power across all regions.

Two FA episodes showed significant differences compared to wakefulness (PERMANOVA, Subj_3_: (F(1,10) = 15.087, p = 0.023, R² = 0.601; Subj_7_: F(1,10) = 12.442, p = 0.020, R² = 0.554), whereas one did not (PERMANOVA, Subj_6_: F(1,10) = 3.069, p = 0.217, R² = 0.235). Both episodes exhibited decreases in fast-frequency activity. Subject 3 showed a significant reduction in low-gamma activity across frontal, central, and temporal regions subject 7 showed decreases in the central region (the only area recorded for this subject). Beta activity was reduced in subject 7 across all regions. Alpha activity decreases in subject 3 and subject 7 across all regions. In slow-frequency bands, subject 3 and subject 7 showed significant increases in delta activity across all regions. Theta activity increased significantly in subject 3 across all regions. See supplementary Tables 82-93 for complete analyses.

## Discussion

In this study, we investigated the neurophysiological bases of conscious states associated with LDs, SP, OBEs, and FAs. First, through PCA, we found that these ten conscious experiences clearly differentiate from wakefulness, indicating that they exhibit distinct electrophysiological profiles that do not fit within the traditional proposal of a hybrid state between wakefulness and sleep^6,8,12^. Instead, they display characteristics of both S1 and REM sleep.

While conscious states during sleep share traits with both REM and S1, they also exhibit specific differences. For example, LDs show a combination of delta and theta contributions, along with moderate involvement from faster frequency bands such as beta and low-gamma. This pattern suggests that LDs share features with REM sleep while also exhibiting more active cortical modulation. However, PERMANOVA showed no significant differences between LDs and REM without consciousness. Baird et al. (2022) proposed that LDs occur within normal REM sleep; however, they found significant reductions in delta (2–4 Hz) and beta (12.5–35 Hz) power during LDs compared to REM^27^. The difference between studies could be due to variations in the selection of segments for analysis. While we included only segments of phasic REM (for REM without consciousness) and incorporated the eye-mark in the LD period analyzed, they included both tonic and phasic REM and excluded the eye-mark from the lucid segment studied. Thus, the inclusion of tonic REM could be increasing delta power in the period of REM without consciousness^28^.

Regarding SP we observed stronger influences from fast-frequency bands (alpha, beta and low-gamma) during the episodes. Furthermore, compared to REM, both episodes exhibited reduced theta relative power and increased low-gamma activity. Both subjects reported full conscious awareness during SP, although only one successfully performed the eye movement marker. Comparisons with S1 revealed consistent reductions in theta power, mixed effects in delta activity, and localized increases in beta and low-gamma activity. Compared to wakefulness, SP episodes were characterized by increased slow-frequency activity (delta and theta) and reduced fast-frequency activity (alpha and beta), underscoring their distinctiveness from other states of consciousness. Our results partially replicate findings by Mainieri et al. (2021), who reported reduced theta and increased alpha during SP compared to quiet wakefulness^12^. The authors introduced the term "lucid paralysis" to describe SP episodes characterized by heightened self-awareness and cognitive clarity, akin to LDs.

Regarding OBEs, they represent a distinct state, distinguishable from both S1 and wakefulness, while exhibiting partial similarities with REM. The two OBE episodes analyzed in PCA showed considerable similarity to REM, primarily characterized by delta and theta frequency bands. However, OBE_1_ demonstrated a reduced influence of fast-frequency bands compared to REM, as well as a greater influence from slow-frequency bands. These findings were replicated in the PERMANOVA results.

Only OBE_1_ exhibited a significant increase in delta power, along with a reduction in high-frequency bands power, when compared to REM. Specifically, OBE is characterized by a notable increase in slow-frequency bands (e.g., delta and theta) and a reduction in fast-frequency bands (e.g., alpha, beta, and low-gamma) relative to S1 and wakefulness. Notably, across all three LDs, delta activity did not increase but remained at REM-like levels. Delta waves have been characterized according to their amplitude and context of occurrence^29^. Within this characterization, low-to-medium amplitude delta waves (5-50 μV) have been identified. Delta waves (<15 μV) have been related to cognitive processing during wakefulness, for instance, being linked to the inhibition of irrelevant neuronal activity during cognitive tasks^30^. They can also be found during REM sleep (5-50 μV) in fronto-central and medial-occipital regions^28^. Unlike high-amplitude slow waves associated with deep sleep or anesthesia states, which reflect generalized cortical deactivation^31^, these delta waves do not indicate such a state. Instead, they represent localized activity in primary sensory areas and have been hypothesized to facilitate disconnection from the external environment^29^.

Moreover, these low-amplitude delta waves are also associated with psychedelic states and have been observed in experiences with DMT and ketamine, where they are linked to increased phenomenological richness^32^. Thus, the delta activity related to OBE experiences, as observed during the three LDs, could potentially be connected to medium-amplitude slow waves, facilitating disconnection from the external environment and immersion in the dreamlike experience.

Regarding FA states, they were influenced by theta, alongside contributions from fast-frequency bands such as alpha, beta, and low-gamma. However, the contributions of these bands varied across the FA episodes with no clear pattern across subjects. Notably, and differently from LDs and OBEs, two subjects (subjects 3 and 6) showed a significant decrease in delta relative power compared to REM; however, the increase observed in subject 6 does not survive corrections for multiple comparisons. It is important to note that subject 7, unlike subjects 3 and 6, left a marker of lucidity during the false awakening. Thus, we can consider that the differences with this subject may be due to the fact that the first two woke up when they recognized they were experiencing an FA, whereas the analyzed fragment of subject 7 represents a "lucid" FA, similar to the proposal by Manieri et al. (2021) for SP^12^.

It is worth noting that we found partially similar results to Manieri et al. (2021), with a significant increase in alpha in one of the subjects, while in the other two, the increase does not survive multiple comparison corrections^12^.

On the one hand, it is important to highlight that, from a methodological perspective, we demonstrate for the first time that OBEs and FAs can be marked using eye movements, similar to LDs, representing a significant advancement in the objective validation of these experiences. On the other hand, from a theoretical point of view, our findings show that conscious states during sleep are clearly distinct from wakefulness, even though they occur within sleep stages. Most previous studies on conscious experiences during sleep focus on the involvement of fast-frequency activity in these states^1,6,12,27,33^. We further propose that delta activity could also play a role, especially in altered states of consciousness, possibly by facilitating disconnection from the external environment and immersion in the dreamlike experience.

## Materials & Methods

### Participants

Participants were recruited via social media advertisements targeting individuals with experiences of LDs, SPs, OBEs or FAs during sleep. Eligibility criteria included experiencing at least one episode of LD, SP, or OBE per week, with no history of psychiatric disorders, substance abuse, epilepsy, or migraines. The study protocol was reviewed and approved by the Biomedical Research Ethics Committee of the Alberto C. Taquini Institute for Translational Medicine Research (IATIMET) and the Human Ethic Committee, Faculty of Medicine, University of Buenos Aires (Comité de Ética Humana, Facultad de Ciencias Médicas, Universidad de Buenos Aires). Informed consent was obtained from all participants prior to their inclusion in the study. Of the initial pool, eleven participants (n = 11) met the eligibility criteria, and seven (n = 7) successfully experienced conscious experiences during sleep.

### Procedure

#### Initial Screening and Training

Participants attended an online interview where they were instructed on maintaining a sleep diary for 15 days and trained to signal episodes of LD, SP, or OBEs using eye movements. The eye movement signal consisted of a specific pattern: moving their eyes as if attempting to look at their ears, repeating the motion three times (left-right, left-right, left-right). Participants who successfully completed this eye-mark signal at least twice in their home environment were invited to the sleep laboratory for further study.

#### In-Lab Procedure

In the sleep laboratory, participants were instructed to perform an eye-mark signal immediately upon becoming aware of a LD, SP, OBE, or FA during polysomnographic recordings. Upon waking from the episode, they provided a verbal report of their experience. Subsequently, two independent observers evaluated the reports to determine whether they corresponded to any of these experiences.

### Polysomnography

Polysomnographic recordings were obtained using electroencephalography systems (BrainVision), including electroencephalography (EEG), electromyography (EMG), and electrooculography (EOG). EEG electrodes were placed at F3, F4, F7, F8, Fz, C3, C4, Cz, T3, T4, T5, T6, P3, P4, Pz, O1, and O2, according to the International 10–20 system, referenced to electrodes attached to both mastoids. The EOG activity was recorded using two electrodes: one placed 1 cm above the right eyebrow and another 1 cm below the left eye. Sleep stages were scored according to standard criteria^34^. Data were sampled at 250 Hz, with electrode impedance maintained below 10 kΩ. A detailed summary of data preprocessing is provided in Table 1.

**Table 1.** Details of Each Subject’s Data Processing. . Summary of data preprocessing for each subject, including the type of conscious experience analyzed, removed EEG channels due to artifacts, availability of eye signaling, and criteria used to define the EEG segment for analysis. EEG segments were either defined arbitrarily (30 s), based on reported awakening, or extended until awakening when possible.

| Subject | Type of conscious experience | Removed Channels | Eye signaling | EEG segment definition |
| --- | --- | --- | --- | --- |
| 1 | Lucid Dream | Fz, F8, C3, C4, T5 | Yes | Arbitrary 30 s segment |
| 2 | Lucid Dream | F4, F7, C3, C4, T3, T4, T5, T6, P3, Pz, P4, O1, O2 | Yes | Arbitrary 30 s segment |
| 3 | Lucid Dream | C3, C4, Cz, T6, Pz, P4, O1, O2 | Yes | Segment ended upon awakening |
|  | OBE | F3, Fz, C3, Cz, P4, O1 | No | 30 s before reported awakening |
|  | OBE | T3, T4, T5,<br>T6, P3, Pz,<br>O1 | Yes | Only available<br>one REM<br>fragment with<br>the<br>experience; an<br>additional<br>REM fragment<br>from a<br>different night<br>was used for<br>comparison |
|  | False<br>Awakening | F4, T5 | No | 30 s before<br>reported<br>awakening |
| 4 | Sleep<br>Paralysis | T4, T6, P3, P5 | No | The segment<br>corresponds to<br>the REM<br>period<br>immediately<br>preceding the<br>subject's<br>report upon<br>awakening. |
| 5 | Sleep<br>Paralysis | F8, T6, O2 | Yes | Until<br>awakening |
| 6 | False<br>Awakening | None | No | 30 s before<br>reported<br>awakening |
| 7 | False<br>Awakening | None (only<br>C3, C4, Cz<br>available) | Yes | Until<br>awakening |

### Details of Each Subject’s Data Processing

### EEG Data Preprocessing

The obtained EEG signal was filtered between 0.16 Hz and 45 Hz, using a 50 Hz notch filter. Then, The EEG data were preprocessed using Independent Component Analysis (ICA) to correct ocular artifacts.

ICA was performed using the MNE Python library with the mne.preprocessing.ICA function^35^. The procedure involves decomposing the EEG signals into a specified number of components. The selection of components to be removed was determined subjectively, based on visual inspection of the independent components.

### EEG Spectral Analysis

To analyze the spectral properties of EEG data across different states of consciousness, segments of phasic REM sleep, S1, wakefulness, and the conscious experience (i.e. lucid dreaming, out-of-body experiences, sleep paralysis, or false awakenings) were divided into consecutive 2.5-second windows with 50% overlap. For each 2.5-second window, the power spectral density (PSD) was estimated using the Welch method, which employs Hanning windows to reduce spectral leakage and improve the accuracy of the PSD estimate. PSD estimates were averaged across all windows for each fragment of EEG data. This averaging process provided a comprehensive measure of spectral power within each state of consciousness. The spectral power was then calculated for specific frequency bands of interest: delta (1-4 Hz), theta (4-8 Hz), alpha (8-13 Hz), beta (13-30 Hz), and low-gamma (30-45 Hz). By averaging the PSD estimates within each state and frequency band, the analysis aimed to identify distinctive spectral features associated with each state of consciousness. All analyses were conducted using the MNE-Python library^35^.

### Calculation of the Area Under the Curve (AUC)

The area under the curve (AUC) for each frequency band was computed using numerical integration with the trapezoidal rule (numpy.trapz from NumPy Python library^36^). Absolute AUC values (always positive) and relative AUC values (each absolute AUC divided by the total sum of AUCs) were obtained. To ensure consistency across different window sizes, relative AUC was used in the analyses. Additionally, the sample size was increased by segmenting the original time windows into smaller sub-windows, recalculating the PSD, and determining the AUC for each frequency band.

### Principal Component Analysis (PCA)

PCA is a dimensionality reduction technique that transforms original variables into uncorrelated components, known as principal components. These components capture the maximum variance, with the first component explaining the most variance and subsequent components explaining progressively less in orthogonal directions. PCA uses the eigenvectors and eigenvalues of the data’s correlation or covariance matrix to define the components, where the eigenvectors determine the projection directions and eigenvalues indicate the variance explained. Metrics like squared cosine (cos²) and contribution (contrib) help interpret the relationship between variables and components.

In this study, PCA was conducted separately for each episode within each condition: lucid dreams (3 episodes), sleep paralysis (2 episodes), and out-of-body experiences (OBEs) (2 episodes) and false awakening (3 episodes). The analysis was performed using relative AUC values of spectral power across five scalp regions—frontal, central, temporal, occipital, and parietal—within five frequency bands: delta, theta, alpha, beta, and low-gamma. PCA was performed in R using the FactoMineR package^37^ for principal component computation and the factoextra package^38^ for visualization and interpretation of results. This approach allowed for an exploration of how spectral power distributions across brain regions contribute to distinct conscious states.

### Permutation-based Multivariate Analysis of Variance (PERMANOVA)

The analysis was conducted in R using the vegan package^39^, with the adonis2 function applied for PERMANOVA (9999 permutations) and betadisper for assessing homogeneity of dispersions. The dependent variable comprised multivariate measures of the AUC of relative spectral power across different frequency bands and brain regions. Initially, PERMANOVA (9999 permutations) was used to compare the AUC across four primary states: wakefulness, S1, REM sleep, and conscious states (LD, SP, OBE, and FA). Following this, pairwise post-hoc comparisons were performed using the pairwise.adonis^40^ function with Bonferroni correction (9999 permutations) to identify specific group differences. A further post-hoc analysis was carried out within each brain region and frequency band to identify variables with significant differences between consciousness states.

### Author Contributions

NLH and CF made substantial contributions to the conception and design of the work; NLH and CF performed the research; NLH acquired the data; NLH performed the statistical analysis; NLH, YC, AAVC and LC performed the EEG analysis; NLH and CF wrote the paper; NLH, YC, AAVC, LC and CF contributed to revising it critically; CF contributed in project administration.

