## Supplementary Material for "ELECTROPHYSIOLOGICAL CORRELATES OF CONSCIOUS EXPERIENCES DURING SLEEP: LUCID DREAMS, SLEEP PARALYSIS, OUT-OF-BODY EXPERIENCES, AND FALSE AWAKENINGS"

### Lucid dreaming

Below are the results of the Principal Component Analysis (PCA) and PERMANOVA for subjects 1, 2, and 3 in the Lucid Dream condition. First, the PCA results are presented, followed by the PERMANOVA results.

**SUBJECT 1, PCA Results**

| Components | Eigenvalue | Variance Explained (%) | Cumulative Variance (%) |
|---|---|---|---|
| 1 | 13.701 | 54.805 | 54.805 |
| 2 | 5.335 | 21.341 | 76.146 |
| 3 | 2.712 | 10.849 | 86.994 |
| 4 | 1.412 | 5.646 | 92.641 |
| 5 | 0.546 | 2.183 | 94.824 |
| 6 | 0.312 | 1.249 | 96.072 |
| 7 | 0.275 | 1.100 | 97.172 |
| 8 | 0.214 | 0.855 | 98.027 |
| 9 | 0.132 | 0.527 | 98.555 |
| 10 | 0.109 | 0.434 | 98.989 |
| 11 | 0.079 | 0.317 | 99.306 |
| 12 | 0.055 | 0.219 | 99.525 |

| | | | |
|---|---|---|---|
| 13 | 0.044 | 0.174 | 99.699 |
| 14 | 0.023 | 0.092 | 99.791 |
| 15 | 0.020 | 0.080 | 99.871 |
| 16 | 0.014 | 0.056 | 99.926 |
| 17 | 0.010 | 0.042 | 99.968 |
| 18 | 0.006 | 0.023 | 99.991 |
| 19 | 0.002 | 0.007 | 99.998 |
| 20 | 0.000 | 0.002 | 100.000 |
| 21 | 0.000 | 0.000 | 100.000 |
| 22 | 0.000 | 0.000 | 100.000 |
| 23 | 0.000 | 0.000 | 100.000 |

**Supplementary Table S1. Lucid dreaming. PCA Results.** Eigenvalues, Percentage of Explained Variance, Cumulative Variance and Main Contributing variables for Principal Components. The first two components account for most of the variance, indicating that a small number of dimensions effectively capture the main structure of the data.

**SUBJECT 1. Contribution of frequency bands to the Principal Components (% Contribution)**

| Variable | Dim.1 | Dim.2 | Dim.3 | Dim.4 | Dim.5 |
|---|---|---|---|---|---|
| Delta Central | 5.150 | 2.585 | 0.476 | 5.728 | 8.500 |
| Theta Central | 4.665 | 0.094 | 0.060 | 14.214 | 15.800 |
| Alpha Central | 6.824 | 0.101 | 1.264 | 0.574 | 0.349 |
| Beta Central | 0.312 | 13.942 | 5.836 | 0.232 | 0.030 |
| Low-Gamma Central | 2.291 | 4.732 | 13.252 | 0.462 | 3.563 |
| Delta Frontal | 4.628 | 2.050 | 0.557 | 12.891 | 0.401 |
| Theta Frontal | 5.456 | 0.042 | 0.050 | 5.282 | 4.296 |
| Alpha Frontal | 6.851 | 0.140 | 0.965 | 1.146 | 0.005 |
| Beta Frontal | 0.018 | 11.812 | 4.610 | 10.654 | 5.118 |
| Low-Gamma Frontal | 2.402 | 4.194 | 13.653 | 0.037 | 3.799 |
| Delta Temporal | 5.728 | 2.133 | 0.039 | 3.540 | 1.811 |

| | | | | | |
|---|---|---|---|---|---|
| Theta Temporal | 5.575 | 0.057 | 1.024 | 8.977 | 0.031 |
| Alpha Temporal | 6.972 | 0.180 | 0.816 | 0.520 | 0.008 |
| Beta Temporal | 0.093 | 13.756 | 5.438 | 3.801 | 5.959 |
| Low-Gamma Temporal | 0.050 | 7.935 | 13.378 | 0.005 | 17.382 |
| Delta Parietal | 5.445 | 2.948 | 0.026 | 3.046 | 2.480 |
| Theta Parietal | 5.314 | 0.039 | 0.171 | 14.786 | 1.862 |
| Alpha Parietal | 6.782 | 0.220 | 1.231 | 0.217 | 0.808 |
| Beta Parietal | 0.505 | 12.638 | 6.551 | 2.436 | 3.236 |
| Low-Gamma Parietal | 1.988 | 4.102 | 16.154 | 1.176 | 2.481 |
| Delta Occipital | 5.977 | 1.281 | 0.390 | 0.383 | 1.225 |
| Theta Occipital | 5.789 | 0.024 | 0.051 | 5.716 | 5.982 |
| Alpha Occipital | 7.010 | 0.180 | 0.747 | 0.002 | 0.226 |
| Beta Occipital | 1.056 | 10.078 | 4.925 | 3.315 | 11.159 |
| Low-Gamma Occipital | 3.117 | 4.738 | 8.337 | 0.861 | 3.488 |

**Supplementary Table S2. Lucid dreaming. PCA results.** Contribution of frequency bands to the Principal Components revealed that Alpha activity contributes the most to Dim.1 (7.01% in Occipital, 6.97% in Temporal), while Beta and Low-Gamma bands are predominant in Dim.2 and Dim.3. Theta and Delta bands show higher contributions in Dim.4 and Dim.5, particularly in the Central and Parietal regions.

**SUBJECT 1. Squared Cosine (Cos²) Values Indicating Variable Representation Across Principal Components**

| Variable | Dim.1 | Dim.2 | Dim.3 | Dim.4 | Dim.5 |
|---|---|---|---|---|---|
| Delta Central | 0.706 | 0.138 | 0.013 | 0.081 | 0.046 |
| Theta Central | 0.639 | 0.005 | 0.002 | 0.201 | 0.086 |
| Alpha Central | 0.935 | 0.005 | 0.034 | 0.008 | 0.002 |
| Beta Central | 0.043 | 0.744 | 0.158 | 0.003 | 0.000 |
| Low-Gamma Central | 0.314 | 0.252 | 0.359 | 0.007 | 0.019 |
| Delta Frontal | 0.634 | 0.109 | 0.015 | 0.182 | 0.002 |
| Theta Frontal | 0.748 | 0.002 | 0.001 | 0.075 | 0.023 |

| | | | | | |
|---|---|---|---|---|---|
| Alpha Frontal | 0.939 | 0.007 | 0.026 | 0.016 | 0.000 |
| Beta Frontal | 0.002 | 0.630 | 0.125 | 0.150 | 0.028 |
| Low-Gamma Frontal | 0.329 | 0.224 | 0.370 | 0.001 | 0.021 |
| Delta Temporal | 0.785 | 0.114 | 0.001 | 0.050 | 0.010 |
| Theta Temporal | 0.764 | 0.003 | 0.028 | 0.127 | 0.000 |
| Alpha Temporal | 0.955 | 0.010 | 0.022 | 0.007 | 0.000 |
| Beta Temporal | 0.013 | 0.734 | 0.147 | 0.054 | 0.033 |
| Low-Gamma Temporal | 0.007 | 0.423 | 0.363 | 0.000 | 0.095 |
| Delta Parietal | 0.746 | 0.157 | 0.001 | 0.043 | 0.014 |
| Theta Parietal | 0.728 | 0.002 | 0.005 | 0.209 | 0.010 |
| Alpha Parietal | 0.929 | 0.012 | 0.033 | 0.003 | 0.004 |
| Beta Parietal | 0.069 | 0.674 | 0.178 | 0.034 | 0.018 |
| Low-Gamma Parietal | 0.272 | 0.219 | 0.438 | 0.017 | 0.014 |
| Delta Occipital | 0.819 | 0.068 | 0.011 | 0.005 | 0.007 |
| Theta Occipital | 0.793 | 0.001 | 0.001 | 0.081 | 0.033 |
| Alpha Occipital | 0.961 | 0.010 | 0.020 | 0.000 | 0.001 |
| Beta Occipital | 0.145 | 0.538 | 0.134 | 0.047 | 0.061 |
| Low-Gamma Occipital | 0.427 | 0.253 | 0.226 | 0.012 | 0.019 |

**Supplementary Table S3. Lucid Dreaming. PCA results.** Squared Cosine (Cos²) values indicate that Alpha activity has the highest representation in Dim.1, particularly in the Occipital (0.961), Temporal (0.955), and Frontal (0.939) regions. Beta activity is primarily represented in Dim.2, while Low-Gamma contributions are more distributed across multiple components. Delta and Theta bands show significant representation in Dim.1 and Dim.4, suggesting their relevance in these dimensions.

**SUBJECT 1. PCA results. Contribution of Conditions to the Principal Components (% Contribution)**

| Condition | Dim.1 | Dim.2 | Dim.3 | Dim.4 | Dim.5 |
|---|---|---|---|---|---|
| LD | 10.802 | 13.221 | 9.731 | 42.958 | 48.340 |
| REM | 10.461 | 29.745 | 1.829 | 34.204 | 35.069 |
| S1 | 7.382 | 50.385 | 79.420 | 19.790 | 15.223 |
| Wakefulness | 71.354 | 6.649 | 9.020 | 3.048 | 1.368 |

**Supplementary Table S4. Lucid Dreaming. PCA results. Contribution of Conditions to the Principal Components (% Contribution).** Wakefulness heavily influences Dim.1, while lucid dreaming exhibits similar contribution in both, dimension 1 and dimension 2. REM and S1 showed a strong contribution in dimension 2.

**SUBJECT 1. PCA results. Squared Cosine ($Cos^2$) Values Indicating Conditions Representation Across Principal Components.**

| Condition | Dim.1 | Dim.2 | Dim.3 | Dim.4 | Dim.5 |
|---|---|---|---|---|---|
| LD | 0.440 | 0.150 | 0.066 | 0.146 | 0.066 |
| REM | 0.342 | 0.294 | 0.018 | 0.163 | 0.062 |
| S1 | 0.204 | 0.290 | 0.262 | 0.088 | 0.045 |
| Wakefulness | 0.925 | 0.032 | 0.023 | 0.004 | 0.001 |

**Supplementary Table 5. Lucid Dreaming. PCA results. Squared Cosine ($Cos^2$) Values Indicating Conditions Representation Across Principal Components.** Dimension 1 showed a better reconstruction of Wakefulness and LD, while the values of REM and S1 were similar in both Dimension 1 and Dimension 2.

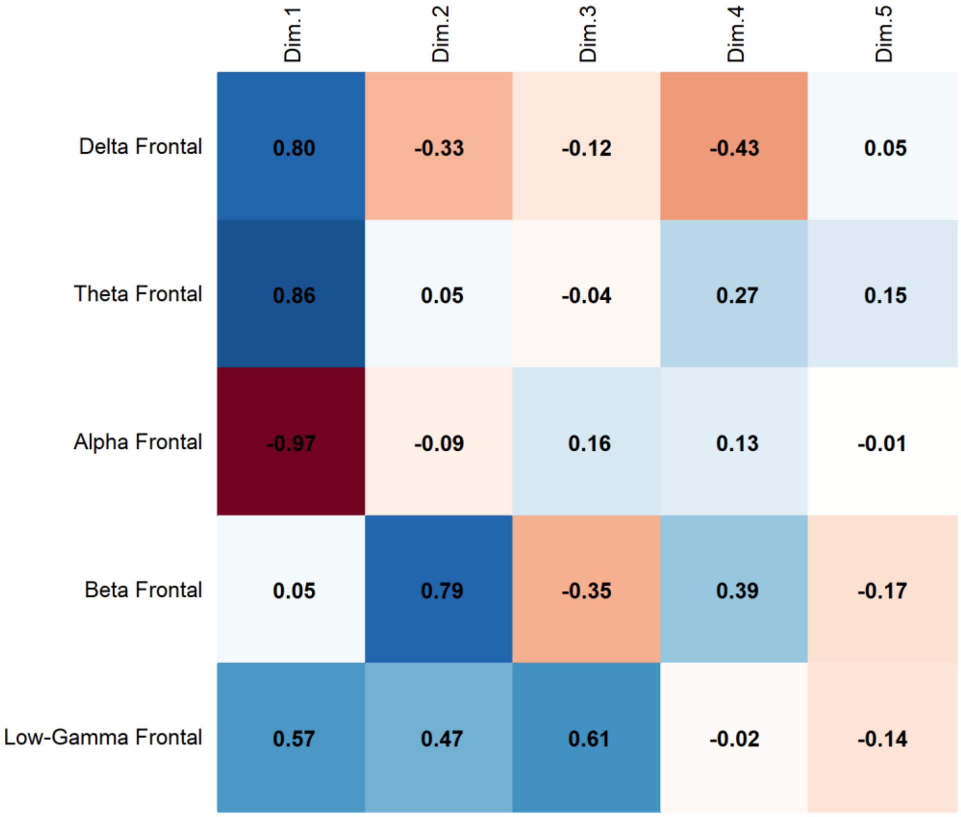

**Supplementary Fig. 1. Lucid Dreaming. PCA results**. Correlation coefficients between frontal band frequencies across dimensions revealed a strong negative correlation between Alpha activity and Dim.1 (-0.97), while Theta (0.86) and Delta (0.80) show strong positive correlations with Dim.1. Beta (0.79) is mainly associated with Dim.2, and Low-Gamma (0.61) with Dim.3

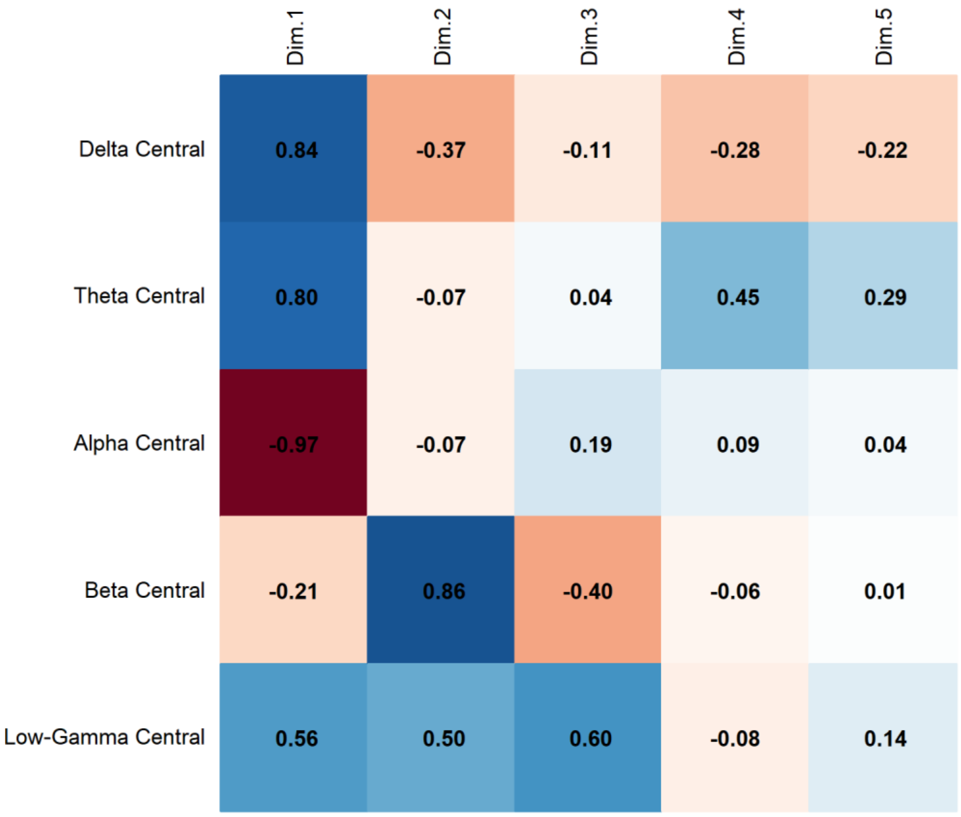


**Supplementary Fig. 2. Lucid Dreaming. PCA Results.** Correlation coefficients between central band frequencies across dimensions revealed that Alpha activity exhibits a strong negative correlation with Dim.1 (-0.97), whereas Delta (0.84) and Theta (0.80) correlate positively with Dim.1. Beta (0.86) is primarily associated with Dim.2, and Low-Gamma (0.60) with Dim.3.

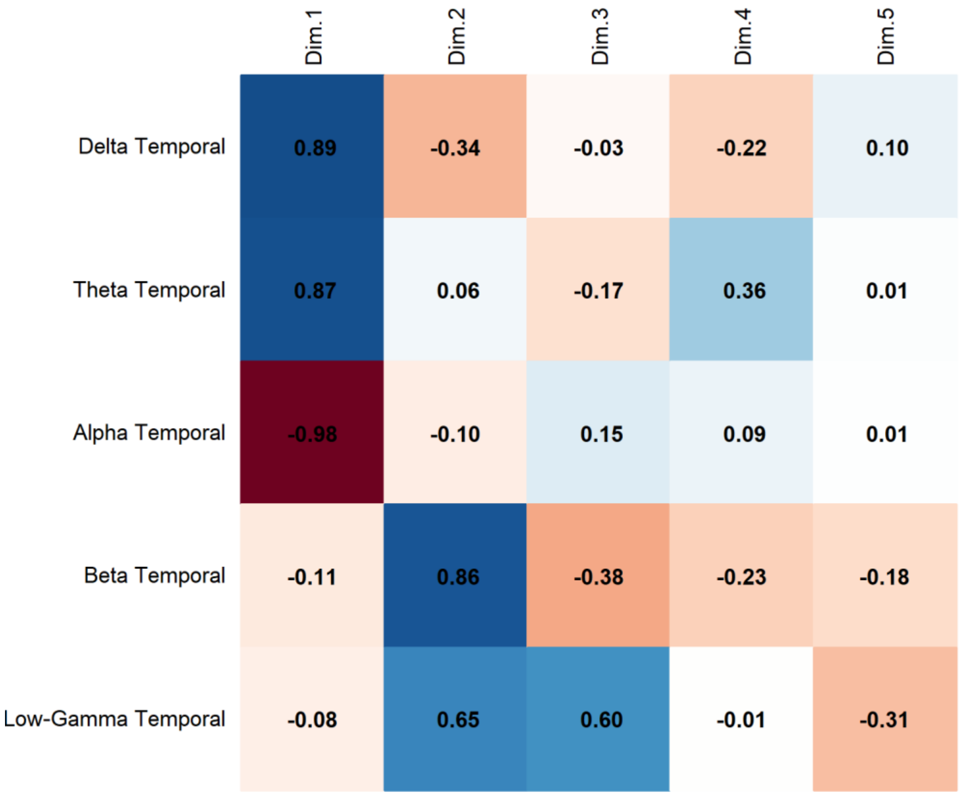


**Supplementary Fig. 3. Lucid Dreaming. PCA Results.** Correlation of Temporal frequencies across dimensions revealed a Strong negative correlation of Alpha activity with Dim.1 (-0.98), while Delta (0.89) and Theta (0.87) show strong positive correlations with Dim.1. Beta (0.86) is most associated with Dim.2, and Low-Gamma (0.65) with Dim.2 and Dim.3 (0.60).

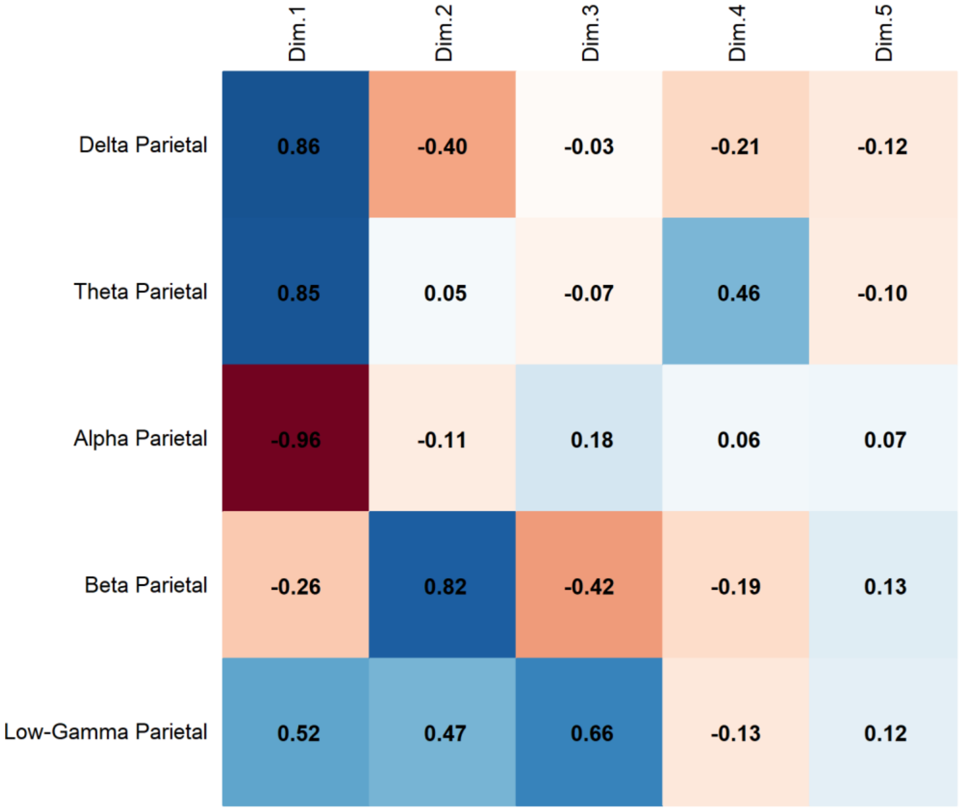


**Supplementary Fig. 4. Lucid Dreaming. PCA Results.** Correlation of Parietal Band Activity with Principal Components revealed that Alpha activity strongly correlates negatively with Dim.1 (-0.96), while Delta (0.86) and Theta (0.85) have strong positive correlations with Dim.1. Beta (0.82) is mainly associated with Dim.2, and Low-Gamma (0.66) with Dim.3

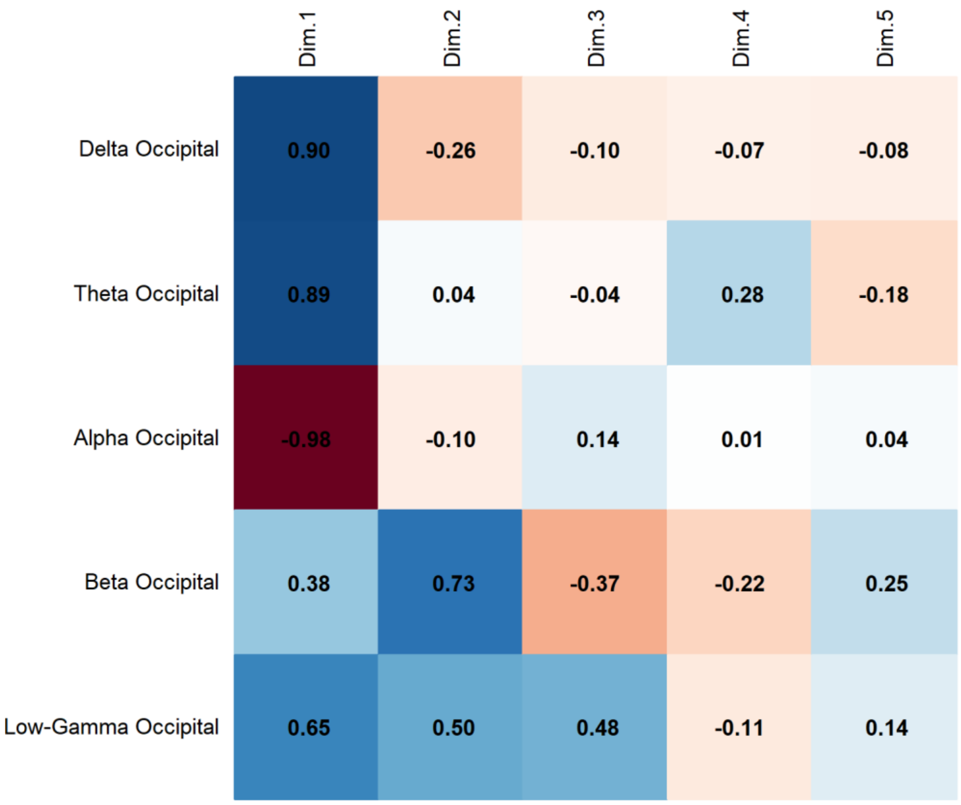


**Supplementary Fig. 5. Lucid Dreaming. PCA Results**. Correlation of Occipital Band Activity with Principal Components revealed that Alpha activity presents a strong negative correlation with Dim.1 (-0.98), while Delta (0.90) and Theta (0.89) correlate positively with Dim.1. Beta (0.73) is associated with Dim.2, and Low-Gamma (0.65) with Dim.1.

**SUBJECT 2. PCA Results.**

| Component | Eigenvalue | Variance Explained (%) | Cumulative Variance (%) |
|---|---|---|---|
| 1 | 5.443 | 54.431 | 54.431 |
| 2 | 1.976 | 19.764 | 74.195 |
| 3 | 0.921 | 9.211 | 83.406 |
| 4 | 0.686 | 6.856 | 90.262 |
| 5 | 0.401 | 4.012 | 94.274 |
| 6 | 0.300 | 3.000 | 97.275 |
| 7 | 0.238 | 2.379 | 99.653 |
| 8 | 0.035 | 0.347 | 100.000 |
| 9 | 0.000 | 0.000 | 100.000 |
| 10 | 0.000 | 0.000 | 100.000 |

**Supplementary Table 6. Lucid Dreaming. PCA results.** Eigenvalues, Percentage of Explained Variance, Cumulative Variance and Main Contributing variables for Principal Components. The first two components account for most of the variance.

**Subject 2. PCA Results. Contribution of Variables to the Principal Components (% Contribution)**

| Variables | Dim.1 | Dim.2 | Dim.3 | Dim.4 | Dim.5 |
|---|---|---|---|---|---|
| Delta Central | 15.111 | 0.086 | 0.994 | 2.624 | 34.173 |
| Theta Central | 8.895 | 10.412 | 2.886 | 21.593 | 12.206 |
| Alpha Central | 14.743 | 6.437 | 2.041 | 0.000 | 7.540 |
| Beta Central | 4.768 | 25.759 | 11.455 | 6.491 | 14.707 |
| Low-Gamma Central | 7.248 | 0.959 | 25.064 | 41.738 | 3.781 |
| Delta Frontal | 11.751 | 3.910 | 26.049 | 0.023 | 3.356 |
| Theta Frontal | 8.935 | 6.919 | 21.332 | 10.443 | 0.001 |
| Alpha Frontal | 13.735 | 5.414 | 9.573 | 0.081 | 0.069 |
| Beta Frontal | 2.289 | 38.217 | 0.602 | 10.872 | 5.539 |
| Low-Gamma Frontal | 12.525 | 1.887 | 0.005 | 6.135 | 18.628 |

**Supplementary Table 7. Lucid Dreaming. PCA Results**. Distribution of the main contributing variables across the first five principal components. Delta Central shows the highest contribution to Dim.1 (15.11%), while Beta Frontal is the dominant contributor to Dim.2 (38.22%).

**Subject 2. PCA results. Squared Cosine (Cos²) Values Indicating Variable Representation Across Principal Components.**

| Variables | Dim.1 | Dim.2 | Dim.3 | Dim.4 | Dim.5 |
|---|---|---|---|---|---|
| Delta Central | 0.822 | 0.002 | 0.009 | 0.018 | 0.137 |
| Theta Central | 0.484 | 0.206 | 0.027 | 0.148 | 0.049 |
| Alpha Central | 0.802 | 0.127 | 0.019 | 0.000 | 0.030 |
| Beta Central | 0.260 | 0.509 | 0.106 | 0.045 | 0.059 |
| Low-Gamma Central | 0.395 | 0.019 | 0.231 | 0.286 | 0.015 |
| Delta Frontal | 0.640 | 0.077 | 0.240 | 0.000 | 0.013 |
| Theta Frontal | 0.486 | 0.137 | 0.196 | 0.072 | 0.000 |
| Alpha Frontal | 0.748 | 0.107 | 0.088 | 0.001 | 0.000 |
| Beta Frontal | 0.125 | 0.755 | 0.006 | 0.075 | 0.022 |
| Low-Gamma Frontal | 0.682 | 0.037 | 0.000 | 0.042 | 0.075 |

**Supplementary Table 8. Lucid Dreaming. PCA Results**. Cos² values indicate how well each variable is represented within the principal components. Delta Central (0.822) and Alpha Central (0.802) are strongly represented in Dim.1, while Beta Frontal (0.755) and Beta Central (0.509) dominate Dim.2.

**Subject 2. PCA Results. Contribution of Conditions to the Principal Components (% Contribution)**

| Condition | Dim.1 | Dim.2 | Dim.3 | Dim.4 | Dim.5 |
|---|---|---|---|---|---|
| LD | 12.854 | 11.181 | 32.878 | 26.543 | 23.128 |
| REM | 23.901 | 7.324 | 26.074 | 27.332 | 20.613 |
| S1 | 4.549 | 64.614 | 2.116 | 20.461 | 15.214 |
| Wakefulness | 58.696 | 16.880 | 38.932 | 25.664 | 41.045 |

**Supplementary Table 9. Lucid Dreaming. PCA results.** Contribution of conditions to the Principal Components revealed that Wakefulness contributes the most to Dim.1 (58.70%), followed by REM (23.90%), while Stage 1 sleep (S1) is predominantly associated with Dim.2 (64.61%). Lucid Dreaming (LD) presents a notable contribution to Dim.3 (32.88%) and Dim.4 (26.54%), suggesting its distinct variance distribution across components.

**Subject 2. PCA results. Squared Cosine (Cos²) Values Indicating Condition Representation Across Principal Components**

| Condition | Dim.1 | Dim.2 | Dim.3 | Dim.4 | Dim.5 |
|---|---|---|---|---|---|
| LD | 0.443 | 0.118 | 0.174 | 0.130 | 0.054 |
| REM | 0.660 | 0.068 | 0.093 | 0.110 | 0.030 |
| S1 | 0.181 | 0.586 | 0.025 | 0.049 | 0.073 |
| Wakefulness | 0.688 | 0.071 | 0.094 | 0.045 | 0.048 |

**Supplementary Table 10. Lucid Dreaming. PCA results.** Squared Cosine (Cos²) values indicate that Wakefulness (0.688) and REM (0.660) have the highest representation in Dim.1, while Stage 1 sleep (S1) is primarily associated with Dim.2. Lucid Dreaming (LD) shows a more distributed representation across dimensions, with its highest loading in Dim.1 (0.443).

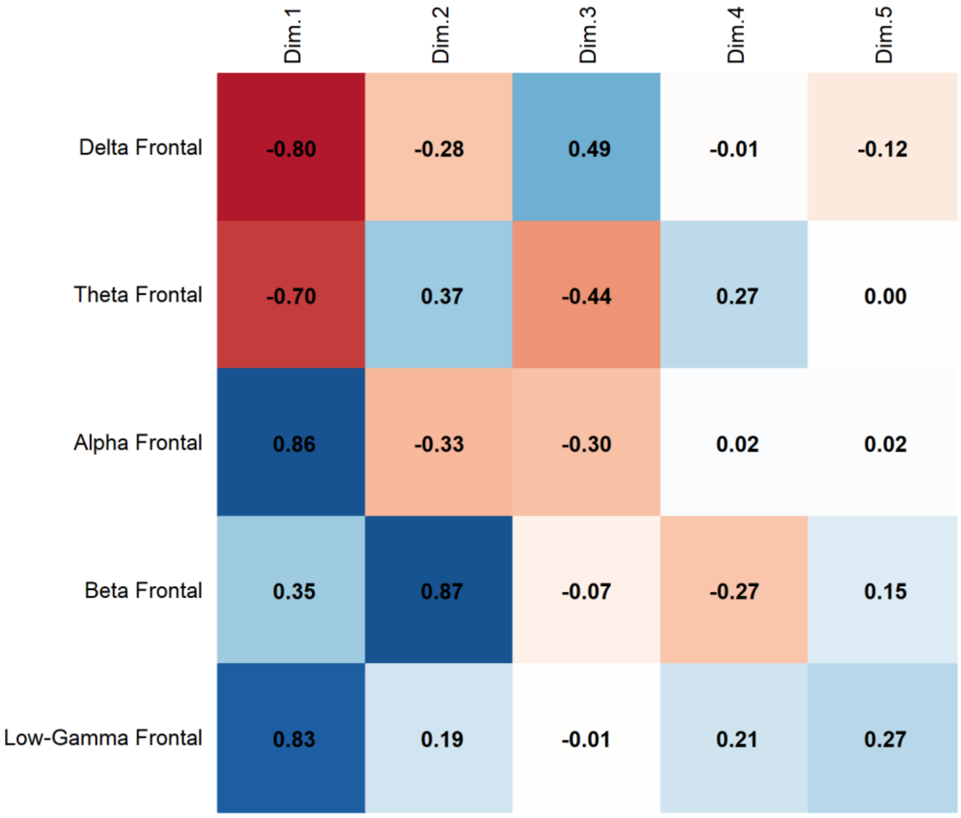


**Supplementary Fig. 6. Lucid Dreaming. PCA Results.** Correlation of Frontal Band Activity with Principal Components revealed that Alpha activity presents a strong positive correlation with Dim.1 (0.86), while Delta (-0.80) and Theta (-0.70) correlate negatively with Dim.1. Beta (0.87) is associated with Dim.2, and Low-Gamma (0.83) with Dim.1.

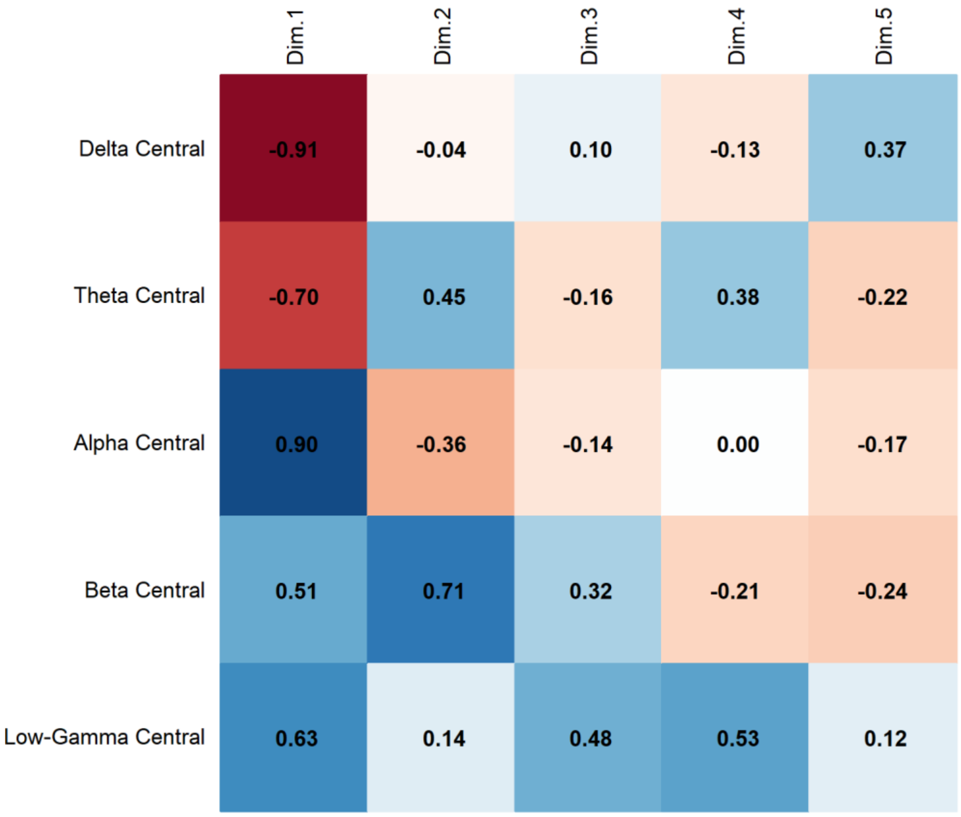


**Supplementary Fig. 7. Lucid Dreaming. PCA Results.** Correlation of Central Band Activity with Principal Components revealed that Alpha activity presents a strong positive correlation with Dim.1 (0.90), while Delta (-0.91) and Theta (-0.70) correlate negatively with Dim.1. Beta (0.71) is associated with Dim.2, and Low-Gamma (0.63) with Dim.1.

**SUBJECT 3. PCA Results.**

| Component | Eigenvalue | Variance Explained (%) | Cumulative Variance (%) |
|---|---|---|---|
| 1 | 8.716 | 58.108 | 58.108 |
| 2 | 3.175 | 21.166 | 79.274 |
| 3 | 1.475 | 9.833 | 89.107 |
| 4 | 0.573 | 3.821 | 92.928 |
| 5 | 0.353 | 2.355 | 95.284 |
| 6 | 0.249 | 1.657 | 96.941 |
| 7 | 0.204 | 1.357 | 98.298 |
| 8 | 0.120 | 0.799 | 99.097 |
| 9 | 0.062 | 0.411 | 99.509 |
| 10 | 0.049 | 0.326 | 99.834 |
| 11 | 0.014 | 0.092 | 99.927 |
| 12 | 0.011 | 0.073 | 100.000 |
| 13 | 0.000 | 0.000 | 100.000 |
| 14 | 0.000 | 0.000 | 100.000 |
| 15 | 0.000 | 0.000 | 100.000 |

**Supplementary Table 11. Lucid Dreaming**. **PCA Results. Eigenvalues, Percentage of Explained Variance, Cumulative Variance and Main Contributing variables for Principal Components**. The first two components account for most of the variance, indicating that a small number of dimensions effectively capture the main structure of the data.

**SUBJECT 3. PCA Results. Contribution of Variables to the Principal Components (% Contribution)**

| Variables | Dim.1 | Dim.2 | Dim.3 | Dim.4 | Dim.5 |
|---|---|---|---|---|---|
| Delta Frontal | 8.337 | 2.531 | 6.312 | 0.532 | 12.172 |
| Theta Frontal | 4.071 | 4.058 | 25.669 | 8.388 | 6.295 |
| Alpha Frontal | 10.225 | 0.304 | 0.263 | 1.501 | 0.941 |
| Beta Frontal | 6.359 | 4.041 | 0.008 | 47.535 | 4.448 |
| Low-Gamma Frontal | 0.036 | 21.107 | 14.570 | 3.680 | 16.465 |
| Delta Temporal | 9.013 | 3.295 | 5.106 | 2.703 | 0.005 |

| | | | | | |
|---|---|---|---|---|---|
| Theta Temporal | 5.342 | 7.359 | 17.651 | 1.133 | 0.208 |
| Alpha Temporal | 11.029 | 0.101 | 0.041 | 4.140 | 1.100 |
| Beta Temporal | 7.311 | 7.323 | 0.054 | 0.383 | 15.185 |
| Low-Gamma Temporal | 1.997 | 21.099 | 6.832 | 4.656 | 0.137 |
| Delta Parietal | 9.463 | 1.643 | 4.105 | 1.097 | 0.087 |
| Theta Parietal | 6.744 | 4.163 | 11.624 | 5.980 | 0.850 |
| Alpha Parietal | 10.613 | 0.232 | 0.006 | 4.220 | 2.087 |
| Beta Parietal | 7.143 | 4.621 | 0.212 | 0.957 | 37.680 |
| Low-Gamma Parietal | 2.318 | 18.124 | 7.547 | 13.093 | 2.341 |

**Supplementary Table 12. Lucid Dreaming. PCA Results. Contribution of Variables to the Principal Components (% Contribution).** Contribution of variables to the Principal Components revealed that Alpha activity has the highest contribution to Dim.1 (10.61% in Parietal, 11.03% in Temporal), while Beta activity dominates Dim.4 (47.53% in Frontal) and Dim.5 (37.68% in Parietal). Low-Gamma is strongly associated with Dim.2 (21.11% in Frontal) and Dim.3 (16.57% in Parietal), indicating its relevance across multiple dimensions.

**Subject 3. PCA Results. Squared Cosine ($Cos^2$) Values Indicating Variable Representation Across Principal Components.**

| Variables | Dim.1 | Dim.2 | Dim.3 | Dim.4 | Dim.5 |
|---|---|---|---|---|---|
| Delta Frontal | 0.727 | 0.080 | 0.093 | 0.003 | 0.043 |
| Theta Frontal | 0.355 | 0.129 | 0.379 | 0.048 | 0.022 |
| Alpha Frontal | 0.891 | 0.010 | 0.004 | 0.009 | 0.003 |
| Beta Frontal | 0.554 | 0.128 | 0.000 | 0.272 | 0.016 |
| Low-Gamma Frontal | 0.003 | 0.670 | 0.215 | 0.021 | 0.058 |
| Delta Temporal | 0.786 | 0.105 | 0.075 | 0.015 | 0.000 |
| Theta Temporal | 0.466 | 0.234 | 0.260 | 0.006 | 0.001 |
| Alpha Temporal | 0.961 | 0.003 | 0.001 | 0.024 | 0.004 |
| Beta Temporal | 0.637 | 0.232 | 0.001 | 0.002 | 0.054 |
| Low-Gamma Temporal | 0.174 | 0.670 | 0.101 | 0.027 | 0.000 |
| Delta Parietal | 0.825 | 0.052 | 0.061 | 0.006 | 0.000 |
| Theta Parietal | 0.588 | 0.132 | 0.171 | 0.034 | 0.003 |

| | | | | | |
|---|---|---|---|---|---|
| Alpha Parietal | 0.925 | 0.007 | 0.000 | 0.024 | 0.007 |
| Beta Parietal | 0.623 | 0.147 | 0.003 | 0.005 | 0.133 |
| Low-Gamma Parietal | 0.202 | 0.575 | 0.111 | 0.075 | 0.008 |

**Supplementary Table 13. Lucid Dreaming. PCA Results.** Squared Cosine (Cos²) values indicate that Alpha activity has the highest representation in Dim.1, particularly in the Temporal (0.961), Parietal (0.925), and Frontal (0.891) regions. Beta activity is mainly represented in Dim.1 and Dim.4, while Low-Gamma has its strongest contribution in Dim.2. Delta and Theta bands show distributed representation across components, with Delta being most relevant in Dim.1.

**SUBJECT 3. PCA Results. Contribution of Conditions to the Principal Components (% Contribution).**

| Condition | Dim.1 | Dim.2 | Dim.3 | Dim.4 | Dim.5 |
|---|---|---|---|---|---|
| LD | 12.944 | 38.610 | 43.395 | 23.777 | 47.943 |
| REM | 21.533 | 43.155 | 32.842 | 18.238 | 5.431 |
| S1 | 6.545 | 9.968 | 22.100 | 33.801 | 13.417 |
| Wakefulness | 58.978 | 8.267 | 1.663 | 24.184 | 33.208 |

**Supplementary Table 14. Lucid Dreaming. PCA Results**. Contribution of conditions to the Principal Components revealed that Lucid Dreaming (LD) has its strongest representation in Dim.5 (47.94%) and Dim.3 (43.39%). REM sleep is highly associated with Dim.2 (43.16%), while Stage 1 sleep (S1) contributes most to Dim.4 (33.80%). Wakefulness is predominantly represented in Dim.1 (58.98%), indicating its distinct role in the principal component structure.

**SUBJECT 3. PCA Results. Squared Cosine (Cos²) Values Indicating Condition Representation Across Dimensions**

| Condition | Dim.1 | Dim.2 | Dim.3 | Dim.4 | Dim.5 |
|---|---|---|---|---|---|
| LD | 0.242 | 0.212 | 0.185 | 0.048 | 0.108 |
| REM | 0.519 | 0.284 | 0.123 | 0.035 | 0.005 |
| S1 | 0.363 | 0.175 | 0.202 | 0.115 | 0.025 |
| Wakefulness | 0.880 | 0.041 | 0.005 | 0.022 | 0.021 |

**Supplementary Table 15. Lucid Dreaming. PCA Results.** Squared Cosine (Cos²) values indicate that Wakefulness has the highest representation in Dim.1 (0.880), followed by REM sleep (0.519). Lucid Dreaming (LD) and Stage 1 sleep (S1) show a more distributed representation across dimensions, with LD contributing most to Dim.2 (0.212) and Dim.3 (0.185), while S1 is primarily associated with Dim.1 (0.363) and Dim.3 (0.202).

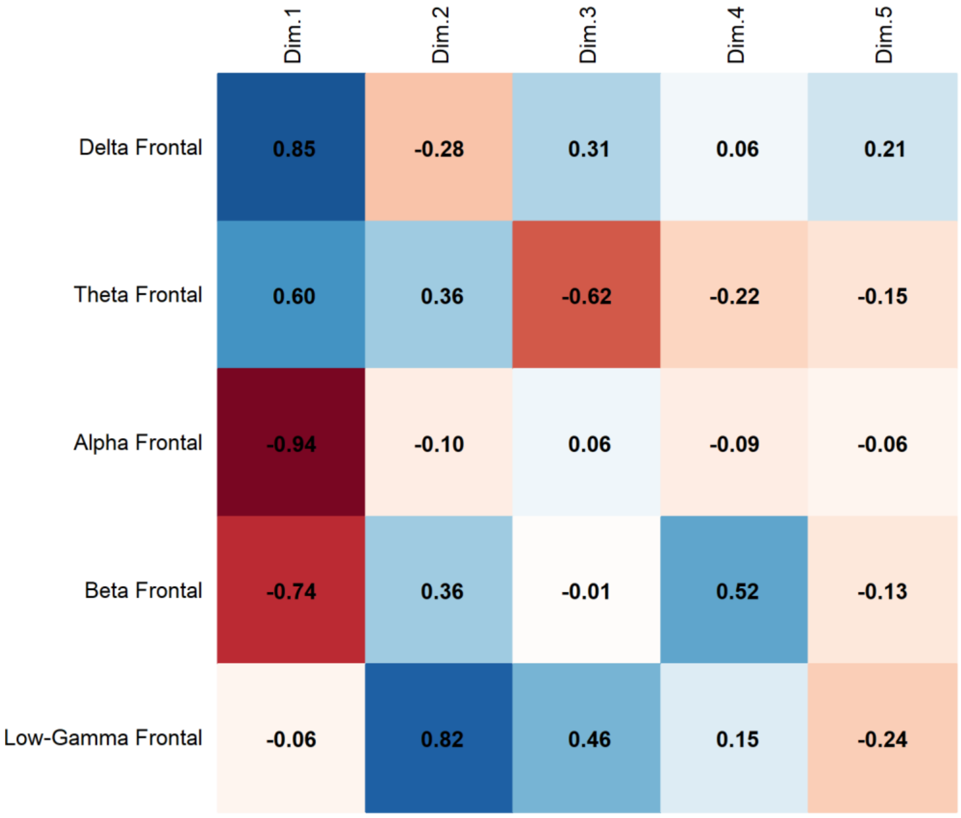


**Supplementary Fig. 8. Lucid Dreaming. PCA Results.** Correlation of Frontal Band Activity with Principal Components revealed that Alpha activity presents a strong negative correlation with Dim.1 (-0.94), while Delta (0.85) and Theta (0.60) correlate positively with Dim.1. Beta (0.52) is associated with Dim.4, and Low-Gamma (0.82) with Dim.2.

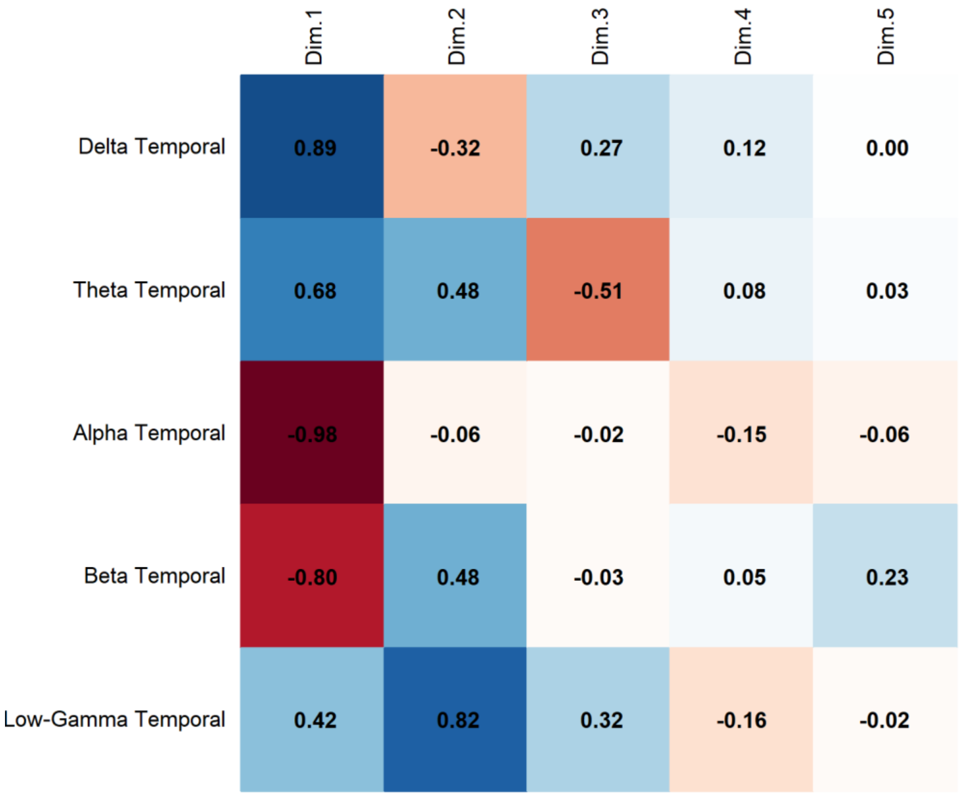


**Supplementary Fig. 9. Lucid Dreaming. PCA Results.** Correlation of Temporal Band Activity with Principal Components revealed that Alpha activity presents a strong negative correlation with Dim.1 (-0.98), while Delta (0.89) and Theta (0.68) correlate positively with Dim.1. Beta (-0.80) is associated with Dim.1, and Low-Gamma (0.82) with Dim.2.

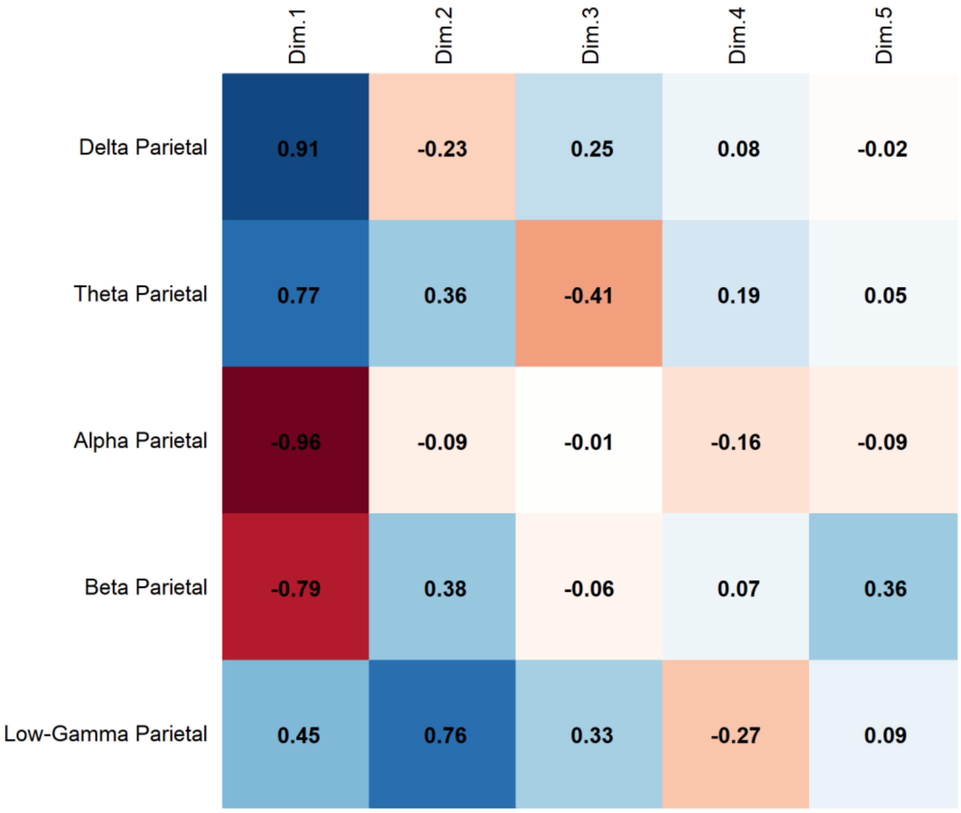


**Supplementary Fig. 10. Lucid Dreaming. PCA Results.** Correlation of Parietal Band Activity with Principal Components revealed that Alpha activity presents a strong negative correlation with Dim.1 (-0.96), while Delta (0.91) and Theta (0.77) correlate positively with Dim.1. Beta (-0.79) is associated with Dim.1, and Low-Gamma (0.76) with Dim.2.

### PERMANOVA

**SUBJECT 1. PERMANOVA Results.**

| | df | Sum of Squares | $R^2$ | F | p-value |
|---|---|---|---|---|---|
| Model | 3 | 8.646 | 0.846 | 36.499 | 2.2e-16 *** |
| Residual | 20 | 1.579 | 0.154 | | |
| Dispersion (Homogeneity) | 3 | 0.065 | | 2.269 | 0.112 |
| Dispersion Residuals | 20 | 0.191 | | | |

**Supplementary Table 16. Lucid Dreaming. PERMANOVA Results**. The model explains 84.6% of the variance ($R^2$ = 0.846) and is statistically significant ($p < 0.001$). The homogeneity of dispersion test suggests no significant differences in variance across groups ($p = 0.112$), indicating that the assumption of homogeneity is met.

**SUBJECT 1. Post-hoc PERMANOVA Results.**

| Comparison | Df | Sums Of Squares | F-value | $R^2$ | p-value | Adjusted p-value |
|---|---|---|---|---|---|---|
| Lucid Dream vs REM | 1 | 0.008144755 | 0.563 | 0.054 | 0.7043 | 1.000 |
| S1 vs Lucid Dream | 1 | 0.041217688 | 3.244 | 0.245 | 0.0362 | 0.217 |
| Lucid Dream vs Wakefulness | 1 | 0.786600437 | 77.596 | 0.886 | 0.0022 | 0.013 |
| REM vs Wakefulness | 1 | 0.842799327 | 94.868 | 0.905 | 0.0021 | 0.013 |
| S1 vs REM | 1 | 0.067925687 | 5.931 | 0.372 | 0.0039 | 0.023 |
| S1 vs Wakefulness | 1 | 0.709940221 | 99.881 | 0.909 | 0.0026 | 0.016 |

**Supplementary Table 17. Lucid Dreaming. Post-hoc PERMANOVA Results.** The model assesses differences between sleep states with Bonferroni-adjusted p-values.

**SUBJECT 1. Comparisons between Lucid Dream and Wakefulness**

| Frequency Band | Brain Region | F-value | $R^2$ | p-value |
|---|---|---|---|---|
| Delta | Frontal | 74.791 | 0.882 | 0.002 |
| | Central | 45.757 | 0.821 | 0.002 |
| | Temporal | 122.019 | 0.924 | 0.002 |
| | Parietal | 33.164 | 0.768 | 0.001 |
| | Occipital | 41.451 | 0.806 | 0.002 |
| Theta | Frontal | 49.154 | 0.831 | 0.002 |
| | Central | 14.808 | 0.597 | 0.003 |
| | Temporal | 68.777 | 0.873 | 0.002 |
| | Parietal | 38.106 | 0.792 | 0.002 |
| | Occipital | 64.802 | 0.866 | 0.002 |
| Alpha | Frontal | 393.831 | 0.975 | 0.003 |

| | | | | |
|---|---|---|---|---|
| | Central | 121.754 | 0.924 | 0.002 |
| | Temporal | 322.076 | 0.970 | 0.002 |
| | Parietal | 104.899 | 0.913 | 0.002 |
| | Occipital | 426.070 | 0.977 | 0.002 |
| Beta | Frontal | 0.198 | 0.019 | 0.671 |
| | Central | 0.289 | 0.028 | 0.619 |
| | Temporal | 0.006 | 0.001 | 0.932 |
| | Parietal | 0.330 | 0.032 | 0.551 |
| | Occipital | 5.115 | 0.338 | 0.019 |
| Low-Gamma | Frontal | 29.245 | 0.745 | 0.003 |
| | Central | 19.194 | 0.657 | 0.005 |
| | Temporal | 6.747 | 0.403 | 0.040 |
| | Parietal | 30.822 | 0.755 | 0.002 |
| | Occipital | 29.781 | 0.749 | 0.003 |

**Supplementary Table 18. Lucid Dreaming. Comparisons between Lucid Dream and Wakefulness.** The model evaluates uncorrected p-values for differences in spectral power across frequency bands and brain regions. Values are uncorrected for multiple comparisons.

**SUBECT 2. PERMANOVA Results.**

| | df | Sum of Squares | $R^2$ | F-value | p-value |
|---|---|---|---|---|---|
| Model | 3 | 3.2549 | 0.78749 | 24.705 | 1e-04 *** |
| Residual | 20 | 0.8783 | 0.21251 | | |
| Dispersion (Homogeneity) | 3 | 0.036846 | | 1.8868 | 0.1644 |
| Dispersion Residuals | 20 | 0.130191 | | | |

**Supplementary Table 19. Lucid Dreaming. PERMANOVA Results.** The model explains 78.7% of the variance ($R^2$ = 0.78749) and is statistically significant ($p < 0.001$, Bonferroni-corrected). The homogeneity of dispersion test does not indicate significant differences in variance across groups ($p = 0.1644$), suggesting that the assumption of homogeneity is met.

**SUBJECT 2. Post-hoc PERMANOVA Results.**

| Comparison | Df | Sum Of Squares | F-value | $R^2$ | p-value | Adjusted p-value |
|---|---|---|---|---|---|---|
| Lucid Dream vs REM | 1 | 0.01524696 | 1.597 | 0.138 | 0.2075 | 1.000 |
| S1 vs Lucid Dream | 1 | 0.13131032 | 12.345 | 0.552 | 0.0028 | 0.017 |
| Lucid Dream vs Wakefulness | 1 | 0.71057642 | 46.144 | 0.822 | 0.0022 | 0.013 |
| REM vs Wakefulness | 1 | 0.75113399 | 46.450 | 0.823 | 0.0022 | 0.013 |
| S1 vs REM | 1 | 0.11097685 | 9.727 | 0.493 | 0.0043 | 0.026 |
| S1 vs Wakefulness | 1 | 0.49785385 | 28.843 | 0.743 | 0.0020 | 0.012 |

**Supplementary Table 20. Lucid Dreaming. Post-hoc PERMANOVA Results.** The model assesses differences between sleep states with Bonferroni-adjusted p-values.

**SUBJECT 2. Comparisons between S1 and Lucid Dream.**

| Frequency Band | Brain Region | F-value | $R^2$ | p-value |
|---|---|---|---|---|
| Delta | Frontal | 24.755 | 0.712 | 0.001 |
| | Central | 8.841 | 0.469 | 0.021 |
| Theta | Frontal | 1.689 | 0.145 | 0.214 |
| | Central | 0.645 | 0.061 | 0.447 |
| Alpha | Frontal | 4.752 | 0.322 | 0.048 |
| | Central | 2.203 | 0.181 | 0.159 |
| Beta | Frontal | 15.908 | 0.614 | 0.004 |
| | Central | 14.779 | 0.596 | 0.004 |
| Low-Gamma | Frontal | 6.017 | 0.376 | 0.002 |
| | Central | 0.097 | 0.010 | 0.761 |

**Supplementary Table 21. Lucid Dreaming. Comparisons between S1 and Lucid Dream.** The model evaluates differences in spectral power across frequency bands and brain regions. Values are uncorrected for multiple comparisons.

**SUBJECT 2. PERMANOVA Results. Comparisons between Lucid Dream and Wakefulness.**

| Frequency Band | Brain Region | F-value | $R^2$ | p-value |
|---|---|---|---|---|
| Delta | Frontal | 17.062 | 0.630 | 0.004 |
| | Central | 90.661 | 0.901 | 0.002 |
| Theta | Frontal | 11.819 | 0.542 | 0.005 |
| | Central | 24.215 | 0.708 | 0.002 |
| Alpha | Frontal | 24.197 | 0.708 | 0.003 |
| | Central | 140.884 | 0.934 | 0.002 |
| Beta | Frontal | 7.722 | 0.436 | 0.019 |
| | Central | 5.061 | 0.336 | 0.051 |
| Low-Gamma | Frontal | 13.609 | 0.576 | 0.002 |
| | Central | 0.994 | 0.090 | 0.332 |

**Supplementary Table 22. Lucid Dreaming. PERMANOVA Results. Comparisons between Lucid Dream and Wakefulness.** The model evaluates differences in spectral power across frequency bands and brain regions. Values are uncorrected for multiple comparisons.

**SUBJECT 3. PERMANOVA Results.**

| | df | Sum of Squares | $R^2$ | F-value | p-value |
|---|---|---|---|---|---|
| Model | 3 | 5.0139 | 0.71407 | 16.649 | 1e-04 *** |
| Residual | 20 | 2.0077 | 0.28593 | | |
| Dispersion (Homogeneity) | 3 | 0.19918 | | 2.2149 | 0.1179 |
| Dispersion Residuals | 20 | 0.59951 | | | |

**Supplementary Table 23. Lucid Dreaming. PERMANOVA Results.** The model explains 71.4% of the variance ($R^2$ = 0.71407) and is statistically significant ($p < 0.001$, Bonferroni-corrected). The homogeneity of dispersion test does not indicate significant differences in variance across groups ($p = 0.1179$), suggesting that the assumption of homogeneity is met.

**SUBJECT 3. Post-hoc PERMANOVA Results**

| Comparison | Df | Sum Of Squares | F-value | $R^2$ | p-value | Adjusted p-value |
|---|---|---|---|---|---|---|
| Lucid Dream vs REM | 1 | 0.09600629 | 2.534 | 0.202 | 0.0870 | 0.522 |
| S1 vs Lucid Dream | 1 | 0.06875371 | 2.303 | 0.187 | 0.0833 | 0.500 |
| Lucid Dream vs Wakefulness | 1 | 0.47219331 | 15.275 | 0.604 | 0.0022 | 0.013 |
| REM vs Wakefulness | 1 | 0.99911575 | 68.220 | 0.872 | 0.0021 | 0.013 |
| S1 vs REM | 1 | 0.05193045 | 3.821 | 0.276 | 0.0188 | 0.113 |
| S1 vs Wakefulness | 1 | 0.75813810 | 114.559 | 0.920 | 0.0026 | 0.016 |

**Supplementary Table 24. Lucid Dreaming. Post-hoc PERMANOVA Results.** The model assesses differences between sleep states with Bonferroni-adjusted p-values.

**SUBJECT 3. Comparisons between Lucid Dream and Wakefulness**

| Frequency Band | Brain Region | F-value | $R^2$ | p-value |
|---|---|---|---|---|
| Delta | Frontal | 15.128 | 0.602 | 0.002 |
| | Temporal | 10.586 | 0.514 | 0.001 |
| | Parietal | 8.569 | 0.461 | 0.004 |
| Theta | Frontal | 6.346 | 0.388 | 0.020 |
| | Temporal | 11.308 | 0.531 | 0.016 |
| | Parietal | 10.760 | 0.518 | 0.013 |
| Alpha | Frontal | 52.793 | 0.841 | 0.002 |
| | Temporal | 27.823 | 0.736 | 0.002 |
| | Parietal | 12.338 | 0.552 | 0.008 |
| Beta | Frontal | 2.645 | 0.209 | 0.144 |
| | Temporal | 3.779 | 0.274 | 0.072 |
| | Parietal | 5.467 | 0.353 | 0.026 |
| Low-Gamma | Frontal | 0.231 | 0.023 | 0.766 |
| | Temporal | 3.708 | 0.271 | 0.029 |
| | Parietal | 2.454 | 0.197 | 0.141 |

**Supplementary Table 25. Lucid Dreaming. Comparisons between Lucid Dream and Wakefulness.** The model evaluates differences in spectral power across frequency bands and brain regions. Values are uncorrected for multiple comparisons.

### SLEEP PARALYSIS

Below are the results of the Principal Component Analysis (PCA) and PERMANOVA for subjects 4 and 5 in the Sleep Paralysis condition. First, the PCA results are presented, followed by the PERMANOVA results.

**SUBJECT 4. PCA Results.**

| Component | Eigenvalue | Variance Explained (%) | Cumulative Variance (%) |
|---|---|---|---|
| 1 | 11.169 | 44.676 | 44.676 |
| 2 | 5.495 | 21.978 | 66.654 |
| 3 | 3.172 | 12.688 | 79.342 |
| 4 | 2.167 | 8.667 | 88.009 |
| 5 | 0.766 | 3.064 | 91.073 |
| 6 | 0.649 | 2.596 | 93.669 |
| 7 | 0.433 | 1.732 | 95.401 |
| 8 | 0.330 | 1.320 | 96.721 |
| 9 | 0.223 | 0.891 | 97.612 |
| 10 | 0.190 | 0.759 | 98.371 |
| 11 | 0.125 | 0.498 | 98.870 |
| 12 | 0.101 | 0.404 | 99.274 |
| 13 | 0.065 | 0.261 | 99.535 |
| 14 | 0.047 | 0.189 | 99.725 |
| 15 | 0.027 | 0.107 | 99.832 |
| 16 | 0.019 | 0.076 | 99.908 |
| 17 | 0.012 | 0.047 | 99.954 |
| 18 | 0.007 | 0.029 | 99.983 |
| 19 | 0.002 | 0.010 | 99.993 |
| 20 | 0.002 | 0.007 | 100.000 |
| 21 | 0.000 | 0.000 | 100.000 |
| 22 | 0.000 | 0.000 | 100.000 |
| 23 | 0.000 | 0.000 | 100.000 |

**Supplementary Table 26. Sleep Paralysis. PCA Results**. Principal Component Analysis (PCA) shows that the first two components account for 66.65% of the total variance, with Component 1 explaining 44.68% and Component 2 explaining 21.98%. The first four components together capture 88.01% of the variance. Variance contributions gradually decrease beyond the fifth component, with cumulative variance reaching 99.83% at Component 15.

**SUBJECT 4. Contribution of Variables to the Dimensions (% Contribution)**

| Variables | Dim.1 | Dim.2 | Dim.3 | Dim.4 | Dim.5 |
|---|---|---|---|---|---|
| Delta Central | 5.120 | 6.310 | 0.391 | 0.574 | 3.519 |
| Theta Central | 4.379 | 3.842 | 5.029 | 0.069 | 0.361 |
| Alpha Central | 7.219 | 2.804 | 0.429 | 0.084 | 0.545 |
| Beta Central | 1.708 | 3.532 | 7.570 | 8.637 | 5.385 |
| Low-Gamma Central | 3.844 | 1.313 | 1.282 | 9.381 | 22.663 |
| Delta Frontal | 3.528 | 0.006 | 11.887 | 0.146 | 17.474 |
| Theta Frontal | 1.628 | 4.539 | 10.996 | 0.181 | 3.563 |
| Alpha Frontal | 5.801 | 2.035 | 0.927 | 2.052 | 5.376 |
| Beta Frontal | 2.147 | 3.154 | 6.234 | 9.970 | 13.544 |
| Low-Gamma Frontal | 1.166 | 12.241 | 0.179 | 1.511 | 0.044 |
| Delta Temporal | 6.497 | 0.908 | 3.920 | 0.128 | 0.086 |
| Theta Temporal | 4.005 | 0.518 | 12.566 | 0.508 | 3.865 |
| Alpha Temporal | 7.246 | 2.005 | 0.000 | 0.622 | 0.502 |
| Beta Temporal | 1.994 | 3.095 | 9.823 | 8.573 | 0.078 |
| Low-Gamma Temporal | 3.116 | 7.118 | 0.347 | 2.819 | 1.231 |
| Delta Parietal | 4.932 | 1.432 | 0.664 | 13.792 | 2.322 |
| Theta Parietal | 6.059 | 0.031 | 3.525 | 3.563 | 4.129 |
| Alpha Parietal | 7.154 | 1.690 | 0.086 | 3.911 | 0.195 |
| Beta Parietal | 0.017 | 2.424 | 9.811 | 18.832 | 2.691 |
| Low-Gamma Parietal | 0.302 | 11.578 | 1.658 | 9.357 | 0.443 |
| Delta Occipital | 7.431 | 0.014 | 1.515 | 0.304 | 4.704 |
| Theta Occipital | 6.076 | 1.235 | 4.078 | 1.962 | 4.878 |
| Alpha Occipital | 8.248 | 0.700 | 0.113 | 0.304 | 0.460 |
| Beta Occipital | 0.062 | 12.138 | 6.699 | 0.209 | 1.856 |
| Low-Gamma Occipital | 0.320 | 15.337 | 0.273 | 2.513 | 0.086 |

**Supplementary Table 27. Sleep Paralysis. Contribution of Variables to the Dimensions.** The table presents the percentage contribution of each frequency band and brain region to the first five principal components. Alpha occipital (8.25%) and delta occipital (7.43%) contribute the most to Dim.1, while low-gamma occipital (15.34%) and beta occipital (12.14%) are the strongest contributors to Dim.2. Delta frontal (11.89%) and theta temporal (12.57%) are prominent in Dim.3, while beta parietal (18.83%) and low-gamma central (9.38%) contribute notably to Dim.4. Low-gamma central (22.66%) and delta frontal (17.47%) show the highest influence on Dim.5.

**SUBJECT 4. Squared Cosine (Cos²) Values Indicating Variable Representation Across Principal Components**

| Variables | Dim.1 | Dim.2 | Dim.3 | Dim.4 | Dim.5 |
|---|---|---|---|---|---|
| Delta Central | 0.572 | 0.347 | 0.012 | 0.012 | 0.027 |
| Theta Central | 0.489 | 0.211 | 0.160 | 0.002 | 0.003 |
| Alpha Central | 0.806 | 0.154 | 0.014 | 0.002 | 0.004 |
| Beta Central | 0.191 | 0.194 | 0.240 | 0.187 | 0.041 |
| Low-Gamma Central | 0.429 | 0.072 | 0.041 | 0.203 | 0.174 |
| Delta Frontal | 0.394 | 0.000 | 0.377 | 0.003 | 0.134 |
| Theta Frontal | 0.182 | 0.249 | 0.349 | 0.004 | 0.027 |
| Alpha Frontal | 0.648 | 0.112 | 0.029 | 0.044 | 0.041 |
| Beta Frontal | 0.240 | 0.173 | 0.198 | 0.216 | 0.104 |
| Low-Gamma Frontal | 0.130 | 0.673 | 0.006 | 0.033 | 0.000 |
| Delta Temporal | 0.726 | 0.050 | 0.124 | 0.003 | 0.001 |
| Theta Temporal | 0.447 | 0.028 | 0.399 | 0.011 | 0.030 |
| Alpha Temporal | 0.809 | 0.110 | 0.000 | 0.013 | 0.004 |
| Beta Temporal | 0.223 | 0.170 | 0.312 | 0.186 | 0.001 |
| Low-Gamma Temporal | 0.348 | 0.391 | 0.011 | 0.061 | 0.009 |
| Delta Parietal | 0.551 | 0.079 | 0.021 | 0.299 | 0.018 |
| Theta Parietal | 0.677 | 0.002 | 0.112 | 0.077 | 0.032 |
| Alpha Parietal | 0.799 | 0.093 | 0.003 | 0.085 | 0.001 |
| Beta Parietal | 0.002 | 0.133 | 0.311 | 0.408 | 0.021 |
| Low-Gamma Parietal | 0.034 | 0.636 | 0.053 | 0.203 | 0.003 |
| Delta Occipital | 0.830 | 0.001 | 0.048 | 0.007 | 0.036 |
| Theta Occipital | 0.679 | 0.068 | 0.129 | 0.043 | 0.037 |
| Alpha Occipital | 0.921 | 0.038 | 0.004 | 0.007 | 0.004 |
| Beta Occipital | 0.007 | 0.667 | 0.212 | 0.005 | 0.014 |
| Low-Gamma Occipital | 0.036 | 0.843 | 0.009 | 0.054 | 0.001 |

**Supplementary Table 28. Sleep Paralysis. Squared Cosine (Cos²) Values for Dimensions 1 and 2**. The variables best represented in Dim.1 are alpha occipital (Cos² = 0.921), delta occipital (Cos² = 0.830), alpha temporal (Cos² = 0.809), and alpha parietal (Cos² = 0.799). In Dim.2, the highest representations are found in low-gamma occipital (Cos² = 0.843), beta occipital (Cos² = 0.667), low-gamma frontal (Cos² = 0.673), and low-gamma parietal (Cos² = 0.636).

**SUBJECT 4. Squared Cosine (Cos²) Values Indicating Condition Representation Across Principal Components**

| Condition | Dim.1 | Dim.2 | Dim.3 | Dim.4 | Dim.5 |
|---|---|---|---|---|---|
| SP | 0.141 | 0.420 | 0.211 | 0.073 | 0.050 |
| REM | 0.498 | 0.073 | 0.179 | 0.052 | 0.050 |
| S1 | 0.415 | 0.039 | 0.114 | 0.268 | 0.011 |
| Wakefulness | 0.666 | 0.127 | 0.071 | 0.048 | 0.028 |

**Supplementary Table 29. Sleep Paralysis. Squared Cosine (Cos²) Values for Condition Representation Across Dimensions.** Wakefulness is best represented in Dim.1 (Cos² = 0.666), followed by REM (Cos² = 0.498) and S1 (Cos² = 0.415). Sleep Paralysis (SP) shows its strongest representation in Dim.2 (Cos² = 0.420), while other conditions contribute less to this dimension.

**SUBJECT 4. Contribution of Conditions to the Principal Components (% Contribution)**

| Condition | Dim.1 | Dim.2 | Dim.3 | Dim.4 | Dim.5 |
|---|---|---|---|---|---|
| SP | 8.355 | 68.268 | 36.553 | 24.487 | 28.119 |
| REM | 17.746 | 5.625 | 22.115 | 9.310 | 27.779 |
| S1 | 15.435 | 3.082 | 18.706 | 42.043 | 5.185 |
| Wakefulness | 58.464 | 23.026 | 22.626 | 24.160 | 38.917 |

**Supplementary Table 30. Sleep Paralysis. Contribution of Conditions to Principal Components.** Wakefulness contributes most to Dim.1 (58.46%), followed by REM (17.75%) and S1 (15.44%). Sleep Paralysis (SP) is the dominant contributor to Dim.2 (68.27%), while Wakefulness also has a notable contribution (23.03%).

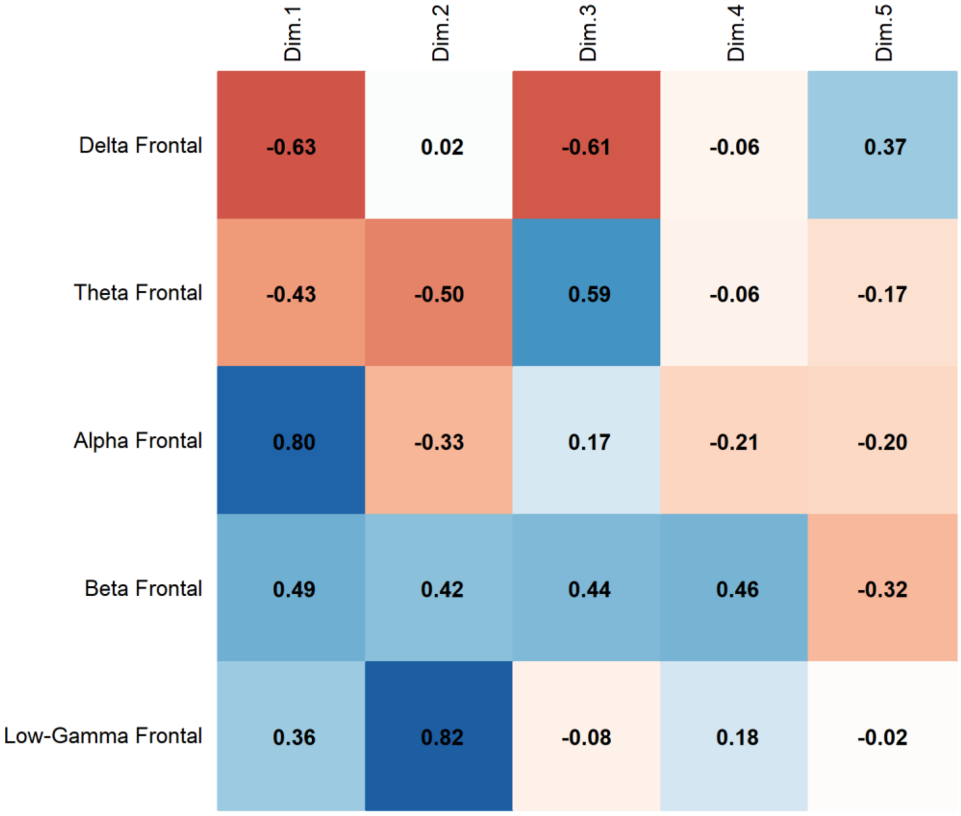


**Supplementary Fig. 11. Sleep Paralysis. PCA Results**. Correlation of Frontal Band Activity with Principal Components revealed that Alpha activity presents a strong positive correlation with Dim.1 (0.80), while Delta (-0.63) and Theta (-0.43) correlate negatively with Dim.1. Beta (0.46) is associated with Dim.4, and Low-Gamma (0.82) with Dim.2.

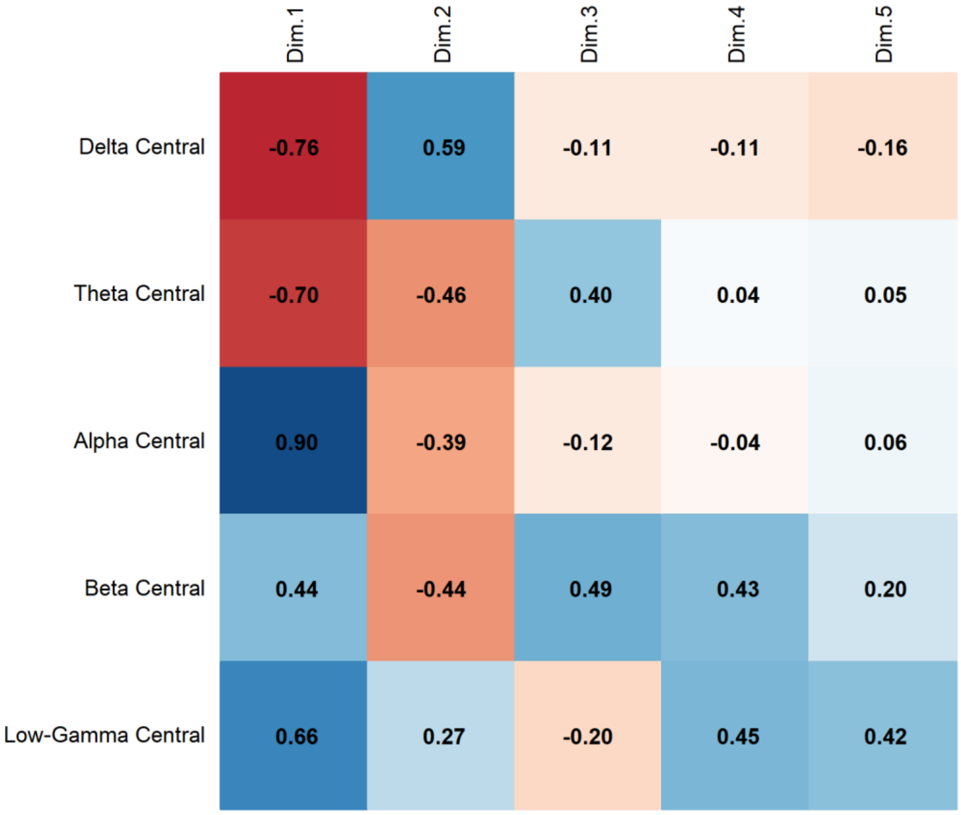


**Supplementary Fig. 12. Sleep Paralysis**. **PCA Results.** Correlation of Central Band Activity with Principal Components revealed that Alpha activity presents a strong positive correlation with Dim.1 (0.90), while Delta (-0.76) and Theta (-0.70) correlate negatively with Dim.1. Beta (0.49) is associated with Dim.3, and Low-Gamma (0.66) with Dim.1.

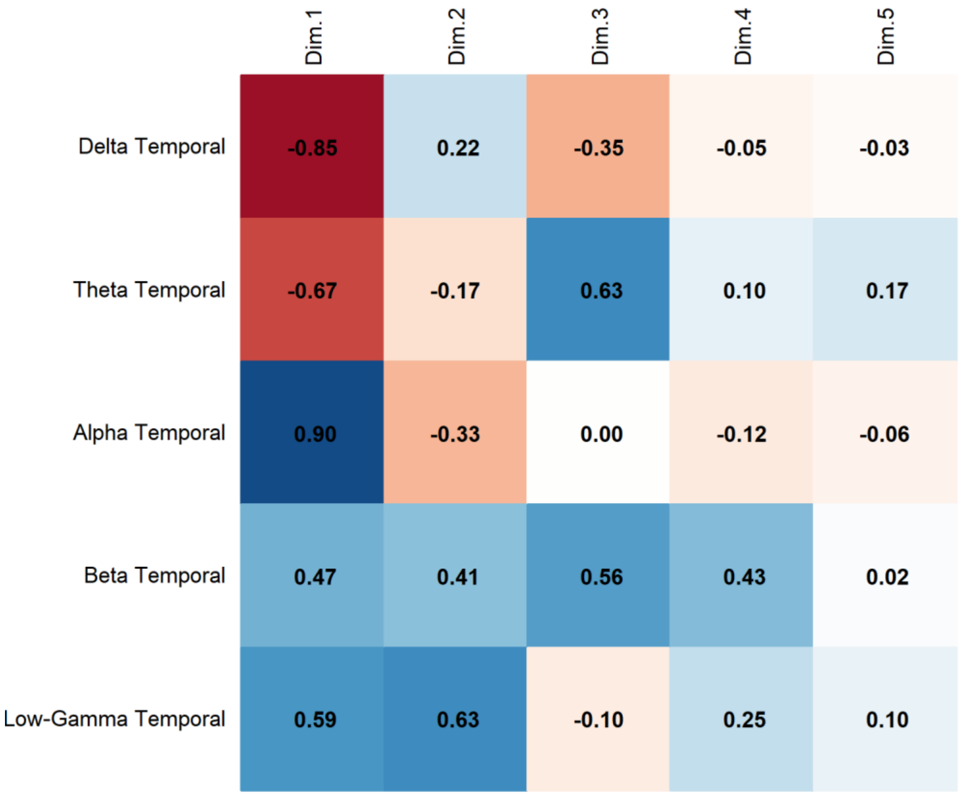


**Supplementary Fig. 13. Sleep Paralysis. PCA Results**. Correlation of Temporal Band Activity with Principal Components revealed that Alpha activity presents a strong positive correlation with Dim.1 (0.90), while Delta (-0.85) and Theta (-0.67) correlate negatively with Dim.1. Beta (0.56) is associated with Dim.3, and Low-Gamma (0.63) with Dim.2.

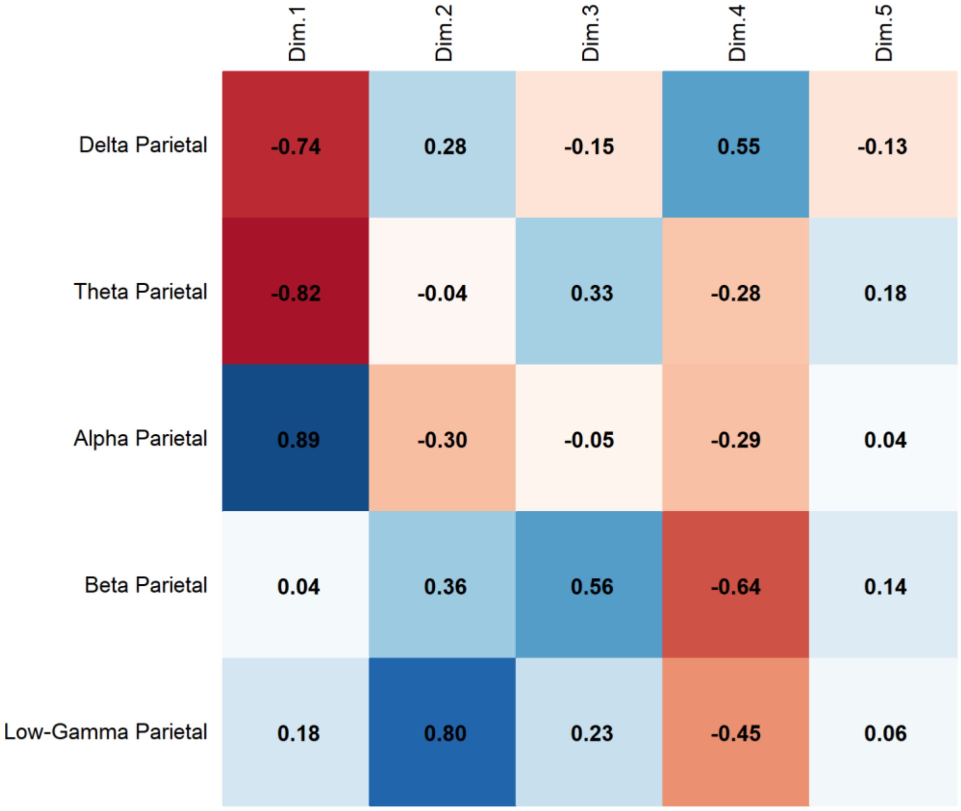


**Supplementary Fig. 14. Sleep Paralysis. PCA Results**. Correlation of Parietal Band Activity with Principal Components revealed that Alpha activity presents a strong positive correlation with Dim.1 (0.89), while Delta (-0.74) and Theta (-0.82) correlate negatively with Dim.1. Beta (0.56) is associated with Dim.3, and Low-Gamma (0.80) with Dim.2.

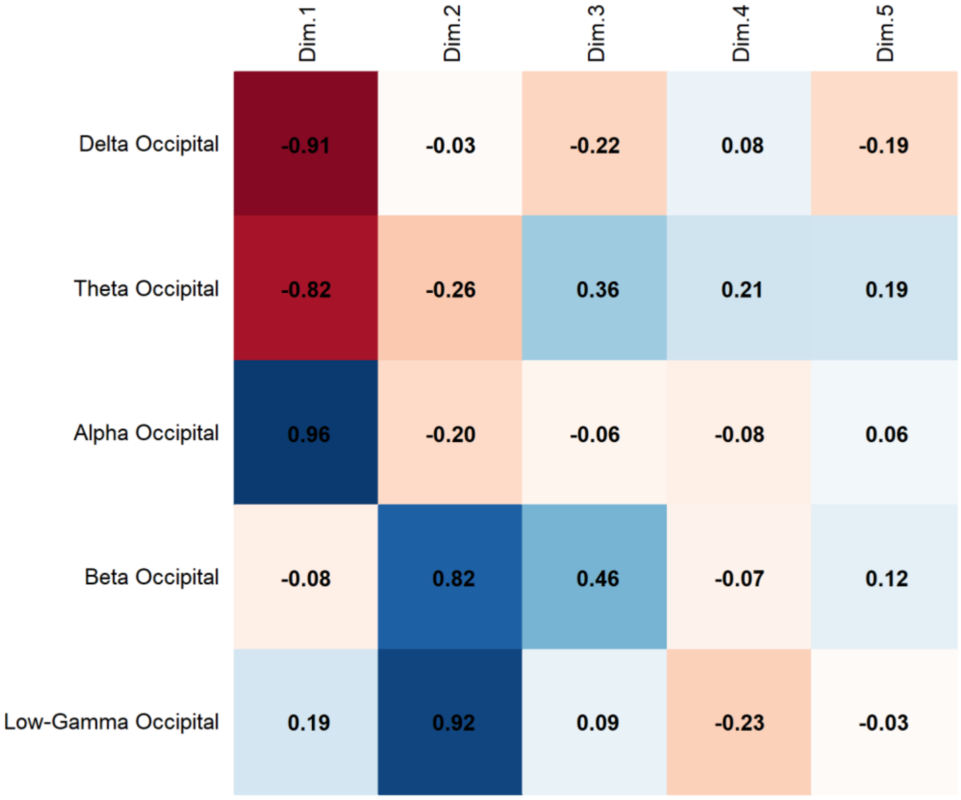


**Supplementary Fig. 15. Sleep Paralysis. PCA Results.** Correlation of Occipital Band Activity with Principal Components revealed that Alpha activity presents a strong positive correlation with Dim.1 (0.96), while Delta (-0.91) and Theta (-0.82) correlate negatively with Dim.1. Beta (0.82) is associated with Dim.2, and Low-Gamma (0.92) with Dim.2.

**SUBJECt 5. PCA Results.**

| Component | Eigenvalue | Variance Explained (%) | Cumulative Variance (%) |
|---|---|---|---|
| 1 | 17.076 | 68.305 | 68.305 |
| 2 | 3.161 | 12.642 | 80.948 |
| 3 | 1.405 | 5.619 | 86.566 |
| 4 | 0.935 | 3.741 | 90.308 |
| 5 | 0.865 | 3.460 | 93.768 |
| 6 | 0.451 | 1.804 | 95.572 |
| 7 | 0.246 | 0.985 | 96.557 |
| 8 | 0.227 | 0.906 | 97.463 |
| 9 | 0.192 | 0.768 | 98.231 |
| 10 | 0.139 | 0.558 | 98.789 |
| 11 | 0.097 | 0.387 | 99.176 |
| 12 | 0.077 | 0.306 | 99.483 |
| 13 | 0.049 | 0.196 | 99.679 |
| 14 | 0.031 | 0.124 | 99.803 |
| 15 | 0.022 | 0.089 | 99.892 |
| 16 | 0.014 | 0.056 | 99.948 |
| 17 | 0.008 | 0.033 | 99.981 |
| 18 | 0.002 | 0.010 | 99.991 |
| 19 | 0.002 | 0.009 | 99.999 |
| 20 | 0.000 | 0.001 | 100.000 |
| 21 | 0.000 | 0.000 | 100.000 |
| 22 | 0.000 | 0.000 | 100.000 |
| 23 | 0.000 | 0.000 | 100.000 |

**Supplementary Table 31. Sleep Paralysis. PCA Results.** Principal Component Analysis (PCA) shows that the first two components account for 80.95% of the total variance, with Component 1 explaining 68.31% and Component 2 explaining 12.64%. The first four components together capture 90.31% of the variance. Variance contributions gradually decrease beyond the fifth component, with cumulative variance reaching 99.89% at Component 15.

**SUBJECT 5. PCA Results. Contribution of Variables to the Principal Components (% Contribution)**

| Variables | Dim.1 | Dim.2 | Dim.3 | Dim.4 | Dim.5 |
|---|---|---|---|---|---|
| Delta Central | 4.794 | 1.165 | 0.948 | 3.869 | 0.229 |
| Theta Central | 4.398 | 2.089 | 8.582 | 0.366 | 2.551 |
| Alpha Central | 2.809 | 7.200 | 0.782 | 2.686 | 21.016 |
| Beta Central | 4.647 | 0.015 | 8.164 | 1.542 | 3.628 |
| Low-Gamma Central | 4.168 | 5.398 | 6.270 | 0.545 | 0.303 |
| Delta Frontal | 2.821 | 4.466 | 4.650 | 21.791 | 5.661 |
| Theta Frontal | 3.695 | 2.238 | 8.797 | 3.157 | 10.872 |
| Alpha Frontal | 1.353 | 15.187 | 0.710 | 11.157 | 12.088 |
| Beta Frontal | 4.552 | 0.332 | 2.214 | 7.735 | 5.277 |
| Low-Gamma Frontal | 4.117 | 5.149 | 6.203 | 0.156 | 0.169 |
| Delta Temporal | 5.017 | 0.440 | 4.009 | 0.001 | 0.198 |
| Theta Temporal | 5.041 | 0.815 | 2.333 | 0.177 | 3.935 |
| Alpha Temporal | 1.305 | 20.523 | 1.119 | 5.183 | 0.001 |
| Beta Temporal | 5.452 | 0.005 | 1.236 | 0.284 | 1.880 |
| Low-Gamma Temporal | 4.258 | 3.044 | 5.249 | 0.004 | 5.116 |
| Delta Parietal | 4.771 | 0.691 | 2.026 | 0.692 | 2.761 |
| Theta Parietal | 4.664 | 0.234 | 3.938 | 0.060 | 2.608 |
| Alpha Parietal | 3.582 | 7.024 | 1.902 | 6.594 | 1.827 |
| Beta Parietal | 4.936 | 0.117 | 5.484 | 0.215 | 4.172 |
| Low-Gamma Parietal | 3.538 | 7.760 | 7.542 | 0.879 | 0.276 |
| Delta Occipital | 4.556 | 0.437 | 0.640 | 6.530 | 8.329 |
| Theta Occipital | 4.852 | 0.026 | 2.761 | 1.587 | 0.003 |
| Alpha Occipital | 3.074 | 5.499 | 0.872 | 23.164 | 1.952 |
| Beta Occipital | 4.463 | 0.356 | 6.559 | 0.127 | 5.137 |
| Low-Gamma Occipital | 3.139 | 9.790 | 7.009 | 1.497 | 0.012 |

**Supplementary Table 32. Sleep Paralysis. Contribution of Variables to Principal Components.** The main contributors to Dim.1 are delta temporal (5.02%), beta temporal (5.45%), and theta occipital (4.85%). In Dim.2, the highest contributions come from alpha temporal (20.52%), alpha frontal (15.19%), and low-gamma occipital (9.79%).

**SUBJECT 5. PCA Results. Squared Cosine ($Cos^2$) Values Indicating Variable Representation Across Principal Components**

| Variables | Dim.1 | Dim.2 | Dim.3 | Dim.4 | Dim.5 |
|---|---|---|---|---|---|
| Delta Central | 0.819 | 0.037 | 0.013 | 0.036 | 0.002 |
| Theta Central | 0.751 | 0.066 | 0.121 | 0.003 | 0.022 |
| Alpha Central | 0.480 | 0.228 | 0.011 | 0.025 | 0.182 |
| Beta Central | 0.794 | 0.000 | 0.115 | 0.014 | 0.031 |
| Low-Gamma Central | 0.712 | 0.171 | 0.088 | 0.005 | 0.003 |
| Delta Frontal | 0.482 | 0.141 | 0.065 | 0.204 | 0.049 |
| Theta Frontal | 0.631 | 0.071 | 0.124 | 0.030 | 0.094 |
| Alpha Frontal | 0.231 | 0.480 | 0.010 | 0.104 | 0.105 |
| Beta Frontal | 0.777 | 0.010 | 0.031 | 0.072 | 0.046 |
| Low-Gamma Frontal | 0.703 | 0.163 | 0.087 | 0.001 | 0.001 |
| Delta Temporal | 0.857 | 0.014 | 0.056 | 0.000 | 0.002 |
| Theta Temporal | 0.861 | 0.026 | 0.033 | 0.002 | 0.034 |
| Alpha Temporal | 0.223 | 0.649 | 0.016 | 0.048 | 0.000 |
| Beta Temporal | 0.931 | 0.000 | 0.017 | 0.003 | 0.016 |
| Low-Gamma Temporal | 0.727 | 0.096 | 0.074 | 0.000 | 0.044 |
| Delta Parietal | 0.815 | 0.022 | 0.028 | 0.006 | 0.024 |
| Theta Parietal | 0.796 | 0.007 | 0.055 | 0.001 | 0.023 |
| Alpha Parietal | 0.612 | 0.222 | 0.027 | 0.062 | 0.016 |
| Beta Parietal | 0.843 | 0.004 | 0.077 | 0.002 | 0.036 |
| Low-Gamma Parietal | 0.604 | 0.245 | 0.106 | 0.008 | 0.002 |
| Delta Occipital | 0.778 | 0.014 | 0.009 | 0.061 | 0.072 |
| Theta Occipital | 0.829 | 0.001 | 0.039 | 0.015 | 0.000 |
| Alpha Occipital | 0.525 | 0.174 | 0.012 | 0.217 | 0.017 |

| | | | | | |
|---|---|---|---|---|---|
| Beta Occipital | 0.762 | 0.011 | 0.092 | 0.001 | 0.044 |
| Low-Gamma Occipital | 0.536 | 0.309 | 0.098 | 0.014 | 0.000 |

**Supplementary Table 33. Sleep Paralysis. Squared Cosine (Cos²) Values for Variable Representation Across Dimensions.** The variables best represented in Dim.1 are beta temporal (Cos² = 0.931), theta temporal (Cos² = 0.861), and delta temporal (Cos² = 0.857). In Dim.2, the highest representations are found in alpha temporal (Cos² = 0.649), alpha frontal (Cos² = 0.480), and low-gamma occipital (Cos² = 0.309).

**SUBJECT 5. PCA Results. Contribution of Conditions to the Principal Components (% Contribution)**

| Condition | Dim.1 | Dim.2 | Dim.3 | Dim.4 | Dim.5 |
|---|---|---|---|---|---|
| SP | 4.781 | 16.209 | 43.593 | 17.652 | 32.607 |
| REM | 29.163 | 10.100 | 7.350 | 17.123 | 23.268 |
| S1 | 14.375 | 26.449 | 24.729 | 16.657 | 16.906 |
| Wakefulness | 51.682 | 47.243 | 24.328 | 48.568 | 27.219 |

**Supplementary Table 34. Sleep Paralysis. Contribution of Conditions to Principal Components.** Wakefulness contributes most to Dim.1 (51.68%), followed by REM (29.16%) and S1 (14.38%). In Dim.2, the highest contributions come from Wakefulness (47.24%) and S1 (26.45%), while Sleep Paralysis (SP) shows its strongest influence in Dim.3 (43.59%).

**SUBJECT 5. PCA Results. Squared Cosine (Cos²) Values Indicating Condition Representation Across Principal Components.**

| Condition | Dim.1 | Dim.2 | Dim.3 | Dim.4 | Dim.5 |
|---|---|---|---|---|---|
| SP | 0.184 | 0.239 | 0.144 | 0.081 | 0.079 |
| REM | 0.798 | 0.071 | 0.017 | 0.023 | 0.027 |
| S1 | 0.580 | 0.179 | 0.067 | 0.036 | 0.030 |
| Wakefulness | 0.758 | 0.127 | 0.029 | 0.044 | 0.020 |

**Supplementary Table 35. Sleep Paralysis. Squared Cosine (Cos²) Values for Condition Representation Across Dimensions.** REM (Cos² = 0.798) and Wakefulness (Cos² = 0.758) are best represented in Dim.1, while Sleep Paralysis (SP) shows its highest representation in Dim.2 (Cos² = 0.239). S1 is moderately represented in both Dim.1 (Cos² = 0.580) and Dim.2 (Cos² = 0.179).

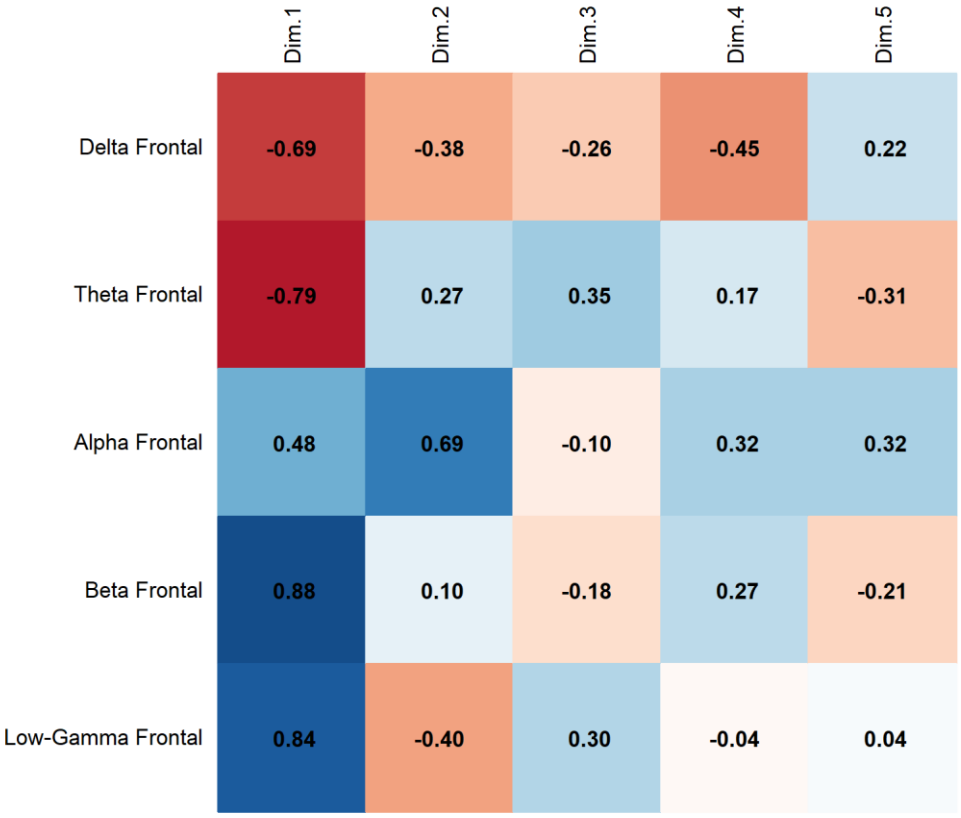


**Supplementary Fig. 16. Sleep Paralysis**, **PCA Results**. Correlation of Frontal Band Activity with Principal Components revealed that Beta activity presents a strong positive correlation with Dim.1 (0.88), while Delta (-0.69) and Theta (-0.79) correlate negatively with Dim.1. Alpha (0.69) is associated with Dim.2, and Low-Gamma (0.84) with Dim.1.

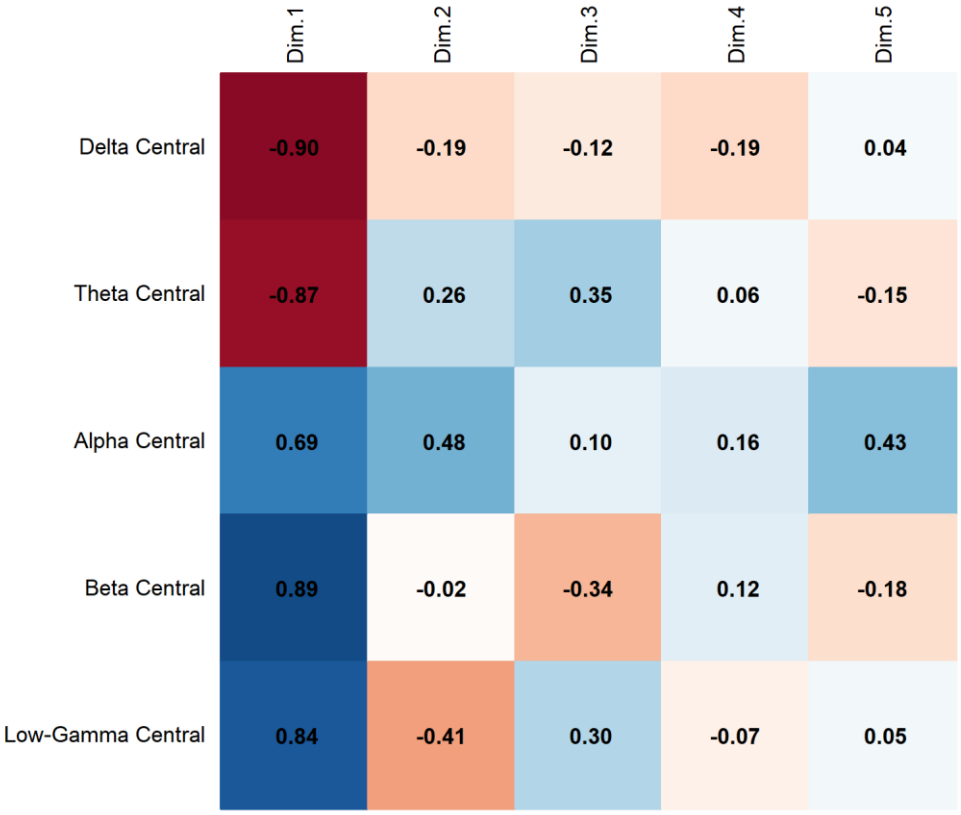


**Supplementary Fig. 17. Sleep Paralysis**, **PCA Results**. Correlation of Central Band Activity with Principal Components revealed that Beta activity presents a strong positive correlation with Dim.1 (0.89), while Delta (-0.90) and Theta (-0.87) correlate negatively with Dim.1. Alpha (0.48) is associated with Dim.2, and Low-Gamma (0.84) with Dim.1.

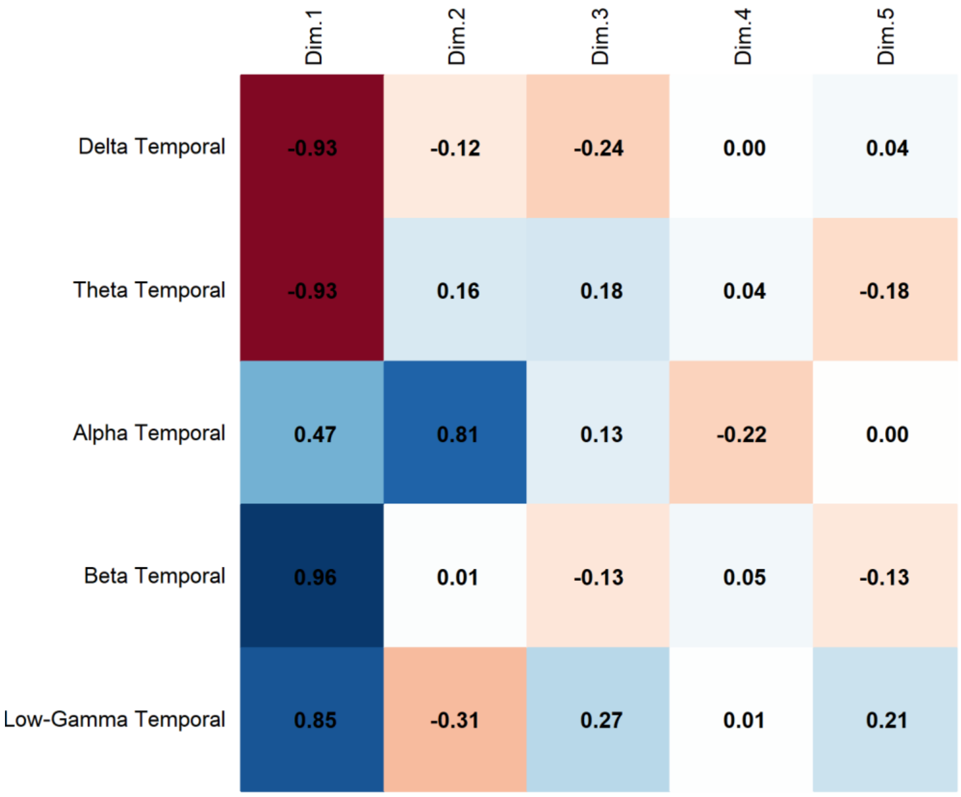


**Supplementary Fig. 18. Sleep Paralysis**, **PCA Results**. Correlation of Temporal Band Activity with Principal Components revealed that Beta activity presents a strong positive correlation with Dim.1 (0.96), while Delta (-0.93) and Theta (-0.93) correlate negatively with Dim.1. Alpha (0.81) is associated with Dim.2, and Low-Gamma (0.85) with Dim.1.

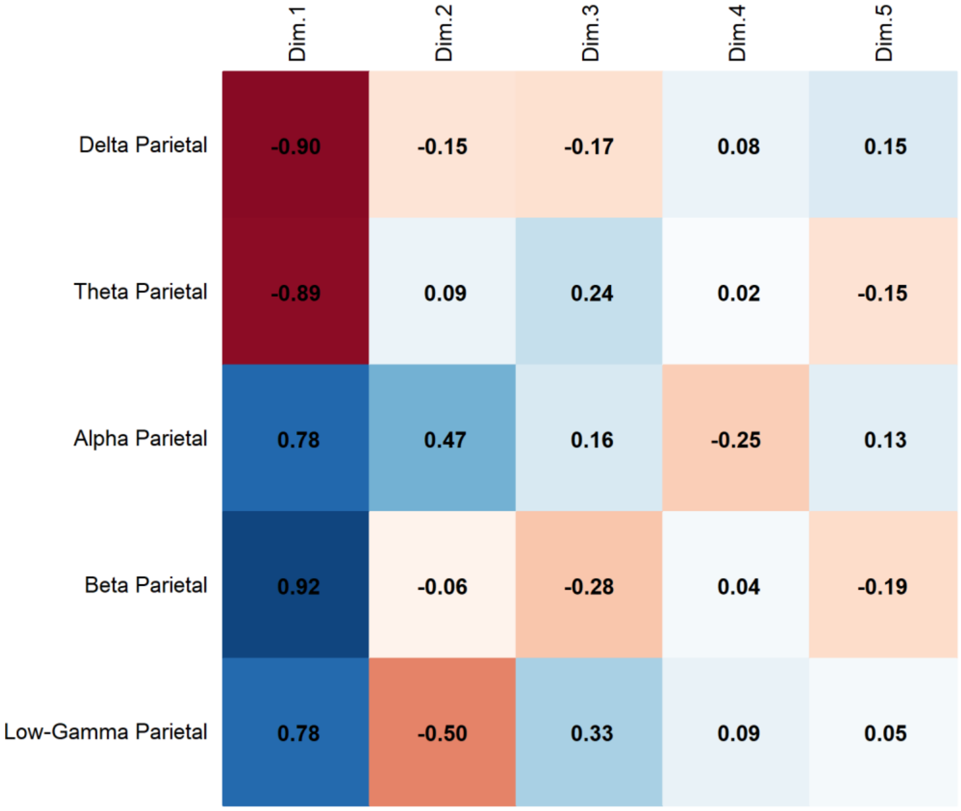


**Supplementary Fig. 19. Sleep Paralysis**. **PCA Results**. Correlation of Parietal Band Activity with Principal Components revealed that Beta activity presents a strong positive correlation with Dim.1 (0.92), while Delta (-0.90) and Theta (-0.89) correlate negatively with Dim.1. Alpha (0.47) is associated with Dim.2, and Low-Gamma (0.78) with Dim.1.

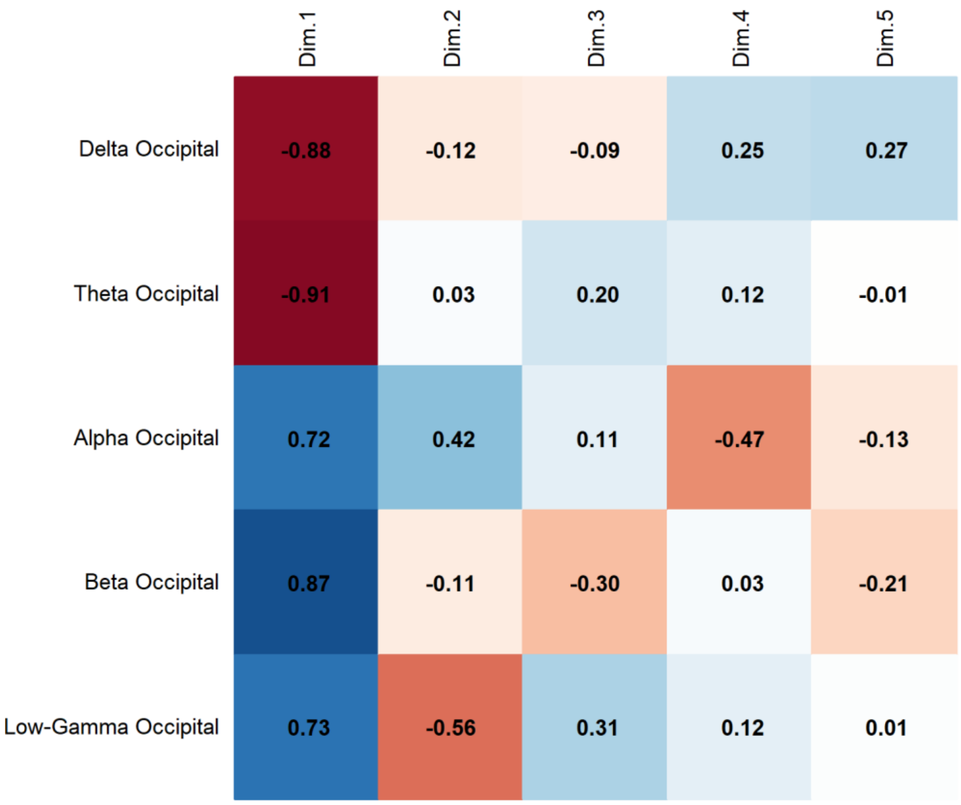


**Supplementary Fig. 20. Sleep Paralysis. PCA Results**. Correlation of Occipital Band Activity with Principal Components revealed that Beta activity presents a strong positive correlation with Dim.1 (0.87), while Delta (-0.88) and Theta (-0.91) correlate negatively with Dim.1. Alpha (0.72) is associated with Dim.1, and Low-Gamma (0.73) with Dim.1.

### PERMANOVA

**SUBJECT 4. PERMANOVA Results.**

| | df | Sum of Squares | $R^2$ | F | p-value |
|---|---|---|---|---|---|
| Model | 3 | 4.9020 | 0.73855 | 18.832 | 1e-04 *** |
| Residual | 20 | 1.7353 | 0.26145 | | |
| Dispersion (Homogeneity) | 3 | 0.07369 | | 1.3843 | 0.2765 |
| Dispersion Residuals | 20 | 0.35489 | | | |

**Supplementary Table 36. Sleep Paralysis. PERMANOVA Results.** The model explains 73.86% of the variance ($R^2$ = 0.73855) and is statistically significant ($p < 0.001$, Bonferroni-corrected). The homogeneity of dispersion test does not indicate significant differences in variance across groups ($p = 0.2765$), suggesting that the assumption of homogeneity is met.

**SUBJECT 4. Post-hoc PERMANOVA Results.**

| Comparison | Df | Sum Of Squares | F-value | $R^2$ | p-value | Adjusted p-value |
|---|---|---|---|---|---|---|
| Sleep Paralysis vs REM | 1 | 0.05324637 | 5.637 | 0.361 | 0.0019 | 0.011 |
| S1 vs Sleep Paralysis | 1 | 0.05541397 | 5.462 | 0.353 | 0.0070 | 0.042 |
| Sleep Paralysis vs Wakefulness | 1 | 0.31220876 | 21.756 | 0.685 | 0.0027 | 0.016 |
| REM vs Wakefulness | 1 | 0.37820861 | 36.783 | 0.786 | 0.0017 | 0.010 |
| S1 vs REM | 1 | 0.01762512 | 2.900 | 0.225 | 0.0298 | 0.179 |
| S1 vs Wakefulness | 1 | 0.39876502 | 36.308 | 0.784 | 0.0024 | 0.014 |

**Supplementary Table 37. Sleep Paralysis. Post-hoc PERMANOVA Results.** The model assesses differences between sleep states with Bonferroni-adjusted p-values.

**SUBJECT 4. Comparisons between Sleep Paralysis and REM**

| Frequency Band | Brain Region | F-value | $R^2$ | p-value |
|---|---|---|---|---|
| Delta | Frontal | 0.188 | 0.018 | 0.672 |
| | Central | 15.815 | 0.613 | 0.006 |
| | Temporal | 0.111 | 0.011 | 0.709 |

| | | | | |
|---|---|---|---|---|
| | Parietal | 0.599 | 0.056 | 0.485 |
| | Occipital | 0.085 | 0.008 | 0.784 |
| Theta | Frontal | 8.234 | 0.452 | 0.020 |
| | Central | 31.307 | 0.758 | 0.005 |
| | Temporal | 24.810 | 0.713 | 0.002 |
| | Parietal | 62.873 | 0.863 | 0.002 |
| | Occipital | 76.238 | 0.884 | 0.002 |
| Alpha | Frontal | 0.054 | 0.005 | 0.854 |
| | Central | 0.327 | 0.032 | 0.571 |
| | Temporal | 1.594 | 0.137 | 0.226 |
| | Parietal | 1.799 | 0.152 | 0.202 |
| | Occipital | 1.418 | 0.124 | 0.261 |
| Beta | Frontal | 3.772 | 0.274 | 0.098 |
| | Central | 4.735 | 0.321 | 0.025 |
| | Temporal | 0.438 | 0.042 | 0.525 |
| | Parietal | 0.049 | 0.005 | 0.829 |
| | Occipital | 1.013 | 0.092 | 0.333 |
| Low-Gamma | Frontal | 9.448 | 0.486 | 0.003 |
| | Central | 4.263 | 0.299 | 0.074 |
| | Temporal | 4.760 | 0.322 | 0.028 |
| | Parietal | 3.654 | 0.268 | 0.101 |
| | Occipital | 7.674 | 0.434 | 0.026 |

**Supplementary Table 38. Sleep Paralysis. Comparisons between Sleep Paralysis and REM.** The model evaluates differences in spectral power across frequency bands and brain regions. Values are uncorrected for multiple comparisons.

**SUBJECT 4. PERMANOVA Results. Comparisons between S1 and Sleep Paralysis.**

| Frequency Band | Brain Region | F-value | $R^2$ | p-value |
|---|---|---|---|---|

| | | | | |
|---|---|---|---|---|
| Delta | Frontal | 0.017 | 0.002 | 0.898 |
| | Central | 17.994 | 0.643 | 0.004 |
| | Temporal | 0.009 | 0.001 | 0.929 |
| | Parietal | 7.960 | 0.443 | 0.013 |
| | Occipital | 0.073 | 0.007 | 0.792 |
| Theta | Frontal | 11.225 | 0.529 | 0.004 |
| | Central | 12.221 | 0.550 | 0.005 |
| | Temporal | 15.416 | 0.607 | 0.002 |
| | Parietal | 1.185 | 0.106 | 0.301 |
| | Occipital | 42.366 | 0.809 | 0.003 |
| Alpha | Frontal | 7.661 | 0.434 | 0.018 |
| | Central | 0.152 | 0.015 | 0.695 |
| | Temporal | 1.523 | 0.132 | 0.233 |
| | Parietal | 15.465 | 0.607 | 0.007 |
| | Occipital | 1.532 | 0.133 | 0.243 |
| Beta | Frontal | 0.013 | 0.001 | 0.906 |
| | Central | 11.549 | 0.536 | 0.002 |
| | Temporal | 0.077 | 0.008 | 0.792 |
| | Parietal | 6.412 | 0.391 | 0.028 |
| | Occipital | 0.462 | 0.044 | 0.510 |
| Low-Gamma | Frontal | 7.300 | 0.422 | 0.011 |
| | Central | 0.000 | 0.000 | 0.994 |
| | Temporal | 4.060 | 0.289 | 0.045 |
| | Parietal | 10.727 | 0.518 | 0.002 |
| | Occipital | 6.839 | 0.406 | 0.049 |

**Supplementary Table 39. Sleep Paralysis. Comparisons between S1 and Sleep Paralysis.** The model evaluates differences in spectral power across frequency bands and brain regions. Significant differences are observed in multiple bands and regions ($p \leq 0.05$). Values are uncorrected for multiple comparisons.

**Subject 4. PERMANOVA Results. Comparisons between Sleep Paralysis and Wakefulness.**

| Frequency Band | Brain Region | F-value | $R^2$ | p-value |
|---|---|---|---|---|
| Delta | Frontal | 3.010 | 0.231 | 0.110 |
| | Central | 150.590 | 0.938 | 0.003 |
| | Temporal | 11.350 | 0.532 | 0.013 |
| | Parietal | 9.255 | 0.481 | 0.019 |
| | Occipital | 10.917 | 0.522 | 0.002 |
| Theta | Frontal | 0.267 | 0.026 | 0.619 |
| | Central | 0.093 | 0.009 | 0.774 |
| | Temporal | 3.273 | 0.247 | 0.095 |
| | Parietal | 22.604 | 0.693 | 0.002 |
| | Occipital | 8.105 | 0.448 | 0.008 |
| Alpha | Frontal | 15.687 | 0.611 | 0.009 |
| | Central | 149.399 | 0.937 | 0.002 |
| | Temporal | 19.814 | 0.665 | 0.003 |
| | Parietal | 25.221 | 0.716 | 0.002 |
| | Occipital | 81.690 | 0.891 | 0.002 |
| Beta | Frontal | 0.001 | 0.000 | 0.979 |
| | Central | 89.334 | 0.899 | 0.002 |
| | Temporal | 0.084 | 0.008 | 0.783 |
| | Parietal | 1.352 | 0.119 | 0.276 |
| | Occipital | 3.712 | 0.271 | 0.101 |
| Low-Gamma | Frontal | 1.331 | 0.117 | 0.252 |
| | Central | 3.303 | 0.248 | 0.063 |
| | Temporal | 0.004 | 0.000 | 0.953 |
| | Parietal | 5.916 | 0.372 | 0.046 |

| | | | | |
|---|---|---|---|---|
| | Occipital | 6.009 | 0.375 | 0.037 |

**Supplementary Table 41. Sleep Paralysis. Comparisons between Sleep Paralysis and Wakefulness.** The model evaluates differences in spectral power across frequency bands and brain regions. Significant differences are observed in multiple bands and regions ($p \leq 0.05$). Values are uncorrected for multiple comparisons.

**Subject 5. PERMANOVA Results.**

| | df | Sum of Squares | R² | F-value | p-value |
|---|---|---|---|---|---|
| Model | 3 | 3.1528 | 0.72036 | 17.174 | 1e-04 *** |
| Residual | 20 | 1.2239 | 0.27964 | | |
| Dispersion (Homogeneity) | 3 | 0.021934 | | 0.8786 | 0.4688 |
| Dispersion Residuals | 20 | 0.166436 | | | |

**Supplementary Table 42. Sleep Paralysis. PERMANOVA Results.** The model explains 72.04% of the variance ($R^2 = 0.72036$) and is statistically significant ($p < 0.001$, Bonferroni-corrected). The homogeneity of dispersion test does not indicate significant differences in variance across groups ($p = 0.4688$).

**Subject 5. Post-hoc PERMANOVA Results.**

| | Df | Sum of Squares | F-value | R² | p-value | p-adjusted |
|---|---|---|---|---|---|---|
| SP vs REM | 1 | 0.15070097 | 17.868521 | 0.6411722 | 0.0026 | 0.0156 |
| SP vs S1 | 1 | 0.06653356 | 5.894393 | 0.3708473 | 0.0053 | 0.0318 |
| SP vs Wakefulness | 1 | 0.11218564 | 11.122590 | 0.5265732 | 0.0022 | 0.0132 |
| REM vs Wakefulness | 1 | 0.51992340 | 78.841853 | 0.8874404 | 0.0024 | 0.0144 |
| REM vs S1 | 1 | 0.03313133 | 4.249882 | 0.2982398 | 0.0051 | 0.0306 |
| S1 vs Wakefulness | 1 | 0.34221172 | 36.219664 | 0.7836419 | 0.0027 | 0.0162 |

**Supplementary Table 43. Sleep Paralysis. Post-hoc PERMANOVA Results**. The model assesses differences between sleep states with Bonferroni-adjusted p-values.

**Subject 5. Comparisons between Sleep Paralysis and REM**

| Frequency Band | Brain Region | F-value | R² | p-value |
|---|---|---|---|---|
| Delta | Frontal | 19.763 | 0.664 | 0.002 |
| | Central | 38.168 | 0.792 | 0.002 |
| | Temporal | 19.102 | 0.656 | 0.004 |
| | Parietal | 23.797 | 0.704 | 0.002 |
| | Occipital | 21.346 | 0.681 | 0.006 |
| Theta | Frontal | 2.621 | 0.208 | 0.130 |

| | | | | |
|---|---|---|---|---|
| | Central | 4.018 | 0.287 | 0.082 |
| | Temporal | 12.590 | 0.557 | 0.008 |
| | Parietal | 14.079 | 0.585 | 0.004 |
| | Occipital | 14.920 | 0.599 | 0.011 |
| Alpha | Frontal | 16.836 | 0.627 | 0.008 |
| | Central | 9.813 | 0.495 | 0.016 |
| | Temporal | 9.092 | 0.476 | 0.011 |
| | Parietal | 10.744 | 0.518 | 0.014 |
| | Occipital | 7.664 | 0.434 | 0.016 |
| Beta | Frontal | 22.775 | 0.695 | 0.002 |
| | Central | 12.645 | 0.558 | 0.005 |
| | Temporal | 18.241 | 0.646 | 0.002 |
| | Parietal | 13.471 | 0.574 | 0.002 |
| | Occipital | 15.121 | 0.602 | 0.002 |
| Low-Gamma | Frontal | 11.859 | 0.543 | 0.012 |
| | Central | 8.881 | 0.470 | 0.018 |
| | Temporal | 5.139 | 0.339 | 0.043 |
| | Parietal | 7.670 | 0.434 | 0.017 |
| | Occipital | 3.459 | 0.257 | 0.090 |

**Supplementary Table 44. Sleep Paralysis. Comparisons between Sleep Paralysis and REM.** The model evaluates differences in spectral power across frequency bands and brain regions. Values are uncorrected for multiple comparisons.

**Subject 5. Comparisons between S1 and Sleep Paralysis**

| Frequency Band | Brain Region | F-value | $R^2$ | p-value |
|---|---|---|---|---|
| Delta | Frontal | 1.069 | 0.097 | 0.329 |
| | Central | 7.864 | 0.440 | 0.027 |
| | Temporal | 0.231 | 0.023 | 0.626 |
| | Parietal | 0.944 | 0.086 | 0.325 |
| | Occipital | 2.953 | 0.228 | 0.114 |

| | | | | |
|---|---|---|---|---|
| Theta | Frontal | 9.607 | 0.490 | 0.013 |
| | Central | 10.309 | 0.508 | 0.002 |
| | Temporal | 13.929 | 0.582 | 0.006 |
| | Parietal | 13.033 | 0.566 | 0.003 |
| | Occipital | 8.951 | 0.472 | 0.015 |
| Alpha | Frontal | 4.863 | 0.327 | 0.056 |
| | Central | 3.292 | 0.248 | 0.078 |
| | Temporal | 0.325 | 0.031 | 0.588 |
| | Parietal | 0.937 | 0.086 | 0.364 |
| | Occipital | 1.063 | 0.096 | 0.312 |
| Beta | Frontal | 8.909 | 0.471 | 0.012 |
| | Central | 7.852 | 0.440 | 0.005 |
| | Temporal | 6.396 | 0.390 | 0.017 |
| | Parietal | 6.588 | 0.397 | 0.014 |
| | Occipital | 9.865 | 0.497 | 0.004 |
| Low-Gamma | Frontal | 2.566 | 0.204 | 0.147 |
| | Central | 15.302 | 0.605 | 0.004 |
| | Temporal | 0.077 | 0.008 | 0.778 |
| | Parietal | 6.408 | 0.391 | 0.034 |
| | Occipital | 1.862 | 0.157 | 0.209 |

**Supplementary Table 45. Sleep Paralysis. Comparisons between S1 and Sleep Paralysis.** The model evaluates differences in spectral power across frequency bands and brain regions. Values are uncorrected for multiple comparisons.

**Subject 5. Comparisons between Sleep Paralysis and Wakefulness.**

| Frequency Band | Brain Region | F-value | $R^2$ | p-value |
|---|---|---|---|---|
| Delta | Frontal | 1.338 | 0.118 | 0.301 |
| | Central | 9.264 | 0.481 | 0.027 |
| | Temporal | 62.499 | 0.862 | 0.002 |

| | | | | |
|---|---|---|---|---|
| | Parietal | 12.269 | 0.551 | 0.011 |
| | Occipital | 4.975 | 0.332 | 0.044 |
| Theta | Frontal | 14.582 | 0.593 | 0.010 |
| | Central | 13.206 | 0.569 | 0.008 |
| | Temporal | 31.190 | 0.757 | 0.001 |
| | Parietal | 11.629 | 0.538 | 0.007 |
| | Occipital | 11.377 | 0.532 | 0.002 |
| Alpha | Frontal | 0.064 | 0.006 | 0.807 |
| | Central | 2.295 | 0.187 | 0.155 |
| | Temporal | 0.088 | 0.009 | 0.769 |
| | Parietal | 4.356 | 0.303 | 0.048 |
| | Occipital | 2.386 | 0.193 | 0.143 |
| Beta | Frontal | 0.909 | 0.083 | 0.344 |
| | Central | 1.179 | 0.105 | 0.312 |
| | Temporal | 10.280 | 0.507 | 0.015 |
| | Parietal | 2.544 | 0.203 | 0.131 |
| | Occipital | 1.403 | 0.123 | 0.254 |
| Low-Gamma | Frontal | 10.683 | 0.517 | 0.012 |
| | Central | 12.208 | 0.550 | 0.005 |
| | Temporal | 57.528 | 0.852 | 0.002 |
| | Parietal | 6.247 | 0.385 | 0.029 |
| | Occipital | 5.812 | 0.368 | 0.029 |

**Supplementary Table 46. Sleep Paralysis. Comparisons between Sleep Paralysis and Wakefulness.** The model evaluates differences in spectral power across frequency bands and brain regions. Values are uncorrected for multiple comparisons.

**Out-of-body Experiences**

The results of the Principal Component Analysis (PCA) and PERMANOVA for Subject 3 in the Out-of-Body Experience (OBE) condition are presented below. This includes two episodes (OBE 1 and OBE 2). First, the PCA results are shown, followed by the PERMANOVA results.

**PCA**

**Subject 3 - OBE1. PCA results**

| Component | Eigenvalue | Variance Explained (%) | Cumulative Variance (%) |
|---|---|---|---|
| 1 | 15.508 | 62.032 | 62.032 |
| 2 | 4.528 | 18.114 | 80.145 |
| 3 | 1.769 | 7.077 | 87.222 |
| 4 | 0.946 | 3.786 | 91.008 |
| 5 | 0.607 | 2.429 | 93.437 |
| 6 | 0.447 | 1.787 | 95.224 |
| 7 | 0.346 | 1.385 | 96.609 |
| 8 | 0.296 | 1.183 | 97.791 |
| 9 | 0.130 | 0.519 | 98.311 |
| 10 | 0.117 | 0.468 | 98.778 |
| 11 | 0.085 | 0.340 | 99.118 |
| 12 | 0.065 | 0.259 | 99.377 |
| 13 | 0.056 | 0.225 | 99.602 |
| 14 | 0.040 | 0.160 | 99.762 |
| 15 | 0.021 | 0.084 | 99.847 |
| 16 | 0.015 | 0.061 | 99.907 |
| 17 | 0.012 | 0.047 | 99.955 |
| 18 | 0.006 | 0.024 | 99.979 |
| 19 | 0.004 | 0.016 | 99.996 |
| 20 | 0.001 | 0.004 | 100.000 |
| 21 | 0.000 | 0.000 | 100.000 |
| 22 | 0.000 | 0.000 | 100.000 |

| | | | |
|---|---|---|---|
| 23 | 0.000 | 0.000 | 100.000 |

**Supplementary Table 47. OBE1. PCA Results.** Principal Component Analysis (PCA) shows that the first two components account for 80.15% of the total variance, with Component 1 explaining 62.03% and Component 2 explaining 18.11%. The first four components together capture 91.01% of the variance.

**SUBJECT 3 - OBE1. Contribution of Variables to the Principal Components (% Contribution).**

| Variables | Dim.1 | Dim.2 | Dim.3 | Dim.4 | Dim.5 |
|---|---|---|---|---|---|
| Delta Central | 5.846 | 0.309 | 1.997 | 0.149 | 0.799 |
| Theta Central | 5.261 | 1.907 | 1.250 | 1.087 | 1.109 |
| Alpha Central | 5.847 | 1.224 | 0.329 | 2.027 | 0.771 |
| Beta Central | 2.808 | 10.488 | 0.708 | 0.502 | 4.660 |
| Low-Gamma Central | 3.762 | 1.043 | 4.888 | 25.653 | 2.497 |
| Delta Frontal | 4.946 | 0.756 | 5.162 | 0.003 | 3.968 |
| Theta Frontal | 2.610 | 7.176 | 8.096 | 2.170 | 3.045 |
| Alpha Frontal | 5.097 | 0.879 | 0.816 | 3.088 | 6.543 |
| Beta Frontal | 4.482 | 4.096 | 1.234 | 1.888 | 2.678 |
| Low-Gamma Frontal | 4.713 | 0.033 | 0.010 | 13.282 | 11.225 |
| Delta Temporal | 5.351 | 0.289 | 5.665 | 1.602 | 1.921 |
| Theta Temporal | 3.372 | 4.135 | 11.907 | 3.549 | 0.823 |
| Alpha Temporal | 5.982 | 1.248 | 0.068 | 0.392 | 0.150 |
| Beta Temporal | 3.269 | 6.958 | 3.310 | 2.922 | 0.496 |
| Low-Gamma Temporal | 0.865 | 9.132 | 5.630 | 0.309 | 37.299 |
| Delta Parietal | 5.793 | 0.017 | 2.356 | 0.266 | 0.032 |
| Theta Parietal | 4.916 | 1.357 | 5.351 | 2.192 | 0.731 |
| Alpha Parietal | 6.005 | 1.328 | 0.035 | 0.004 | 0.000 |
| Beta Parietal | 2.624 | 10.913 | 0.015 | 3.925 | 3.164 |
| Low-Gamma Parietal | 0.514 | 9.976 | 16.518 | 4.637 | 1.268 |
| Delta Occipital | 5.320 | 0.159 | 1.050 | 0.191 | 0.047 |

| | | | | | |
|---|---|---|---|---|---|
| Theta Occipital | 4.550 | 1.779 | 5.794 | 1.690 | 0.381 |
| Alpha Occipital | 5.632 | 2.096 | 0.156 | 0.110 | 0.779 |
| Beta Occipital | 0.084 | 13.262 | 0.020 | 28.292 | 14.897 |
| Low-Gamma Occipital | 0.349 | 9.440 | 17.635 | 0.070 | 0.719 |

**Supplementary Table 48. OBE1. Contribution of Variables to Principal Components.** The main contributors to Dim.1 are alpha parietal (6.01%), alpha temporal (5.98%), and delta central (5.85%). In Dim.2, the highest contributions come from beta occipital (13.26%), beta parietal (10.91%), and low-gamma parietal (9.98%).

**SUBJECT 3 - OBE1. Squared Cosine (Cos²) Values Indicating Variable Representation Across Principal Components**

| Variables | Dim.1 | Dim.2 | Dim.3 | Dim.4 | Dim.5 |
|---|---|---|---|---|---|
| Delta Central | 0.907 | 0.014 | 0.035 | 0.001 | 0.005 |
| Theta Central | 0.816 | 0.086 | 0.022 | 0.010 | 0.007 |
| Alpha Central | 0.907 | 0.055 | 0.006 | 0.019 | 0.005 |
| Beta Central | 0.435 | 0.475 | 0.013 | 0.005 | 0.028 |
| Low-Gamma Central | 0.583 | 0.047 | 0.086 | 0.243 | 0.015 |
| Delta Frontal | 0.767 | 0.034 | 0.091 | 0.000 | 0.024 |
| Theta Frontal | 0.405 | 0.325 | 0.143 | 0.021 | 0.018 |
| Alpha Frontal | 0.790 | 0.040 | 0.014 | 0.029 | 0.040 |
| Beta Frontal | 0.695 | 0.186 | 0.022 | 0.018 | 0.016 |
| Low-Gamma Frontal | 0.731 | 0.002 | 0.000 | 0.126 | 0.068 |
| Delta Temporal | 0.830 | 0.013 | 0.100 | 0.015 | 0.012 |
| Theta Temporal | 0.523 | 0.187 | 0.211 | 0.034 | 0.005 |
| Alpha Temporal | 0.928 | 0.057 | 0.001 | 0.004 | 0.001 |
| Beta Temporal | 0.507 | 0.315 | 0.059 | 0.028 | 0.003 |
| Low-Gamma Temporal | 0.134 | 0.414 | 0.100 | 0.003 | 0.226 |
| Delta Parietal | 0.898 | 0.001 | 0.042 | 0.003 | 0.000 |
| Theta Parietal | 0.762 | 0.061 | 0.095 | 0.021 | 0.004 |

| | | | | | |
|---|---|---|---|---|---|
| Alpha Parietal | 0.931 | 0.060 | 0.001 | 0.000 | 0.000 |
| Beta Parietal | 0.407 | 0.494 | 0.000 | 0.037 | 0.019 |
| Low-Gamma Parietal | 0.080 | 0.452 | 0.292 | 0.044 | 0.008 |
| Delta Occipital | 0.825 | 0.007 | 0.019 | 0.002 | 0.000 |
| Theta Occipital | 0.706 | 0.081 | 0.103 | 0.016 | 0.002 |
| Alpha Occipital | 0.873 | 0.095 | 0.003 | 0.001 | 0.005 |
| Beta Occipital | 0.013 | 0.601 | 0.000 | 0.268 | 0.090 |
| Low-Gamma Occipital | 0.054 | 0.427 | 0.312 | 0.001 | 0.004 |

**Supplementary Table 49. OBE1. Squared Cosine (Cos²) Values for Variable Representation Across Dimensions.** The variables best represented in Dim.1 are alpha parietal (Cos² = 0.931), alpha temporal (Cos² = 0.928), and delta central (Cos² = 0.907). In Dim.2, the highest representations are found in beta occipital (Cos² = 0.601), beta parietal (Cos² = 0.494), and low-gamma parietal (Cos² = 0.452).

**SUBJECT 3 – OBE1. Contribution of Conditions to the Principal Components (% Contribution)**

| Condition | Dim.1 | Dim.2 | Dim.3 | Dim.4 | Dim.5 |
|---|---|---|---|---|---|
| OBE | 20.363 | 16.331 | 27.484 | 14.447 | 13.273 |
| REM | 5.354 | 25.679 | 19.927 | 10.424 | 33.383 |
| S1 | 8.120 | 44.592 | 36.158 | 12.357 | 18.226 |
| Wakefulness | 66.164 | 13.398 | 16.430 | 62.771 | 35.117 |

**Supplementary Table 50. OBE1. Contribution of Conditions to Principal Components**. Wakefulness contributes most to Dim.1 (66.16%) and Dim.4 (62.77%). S1 has the highest contribution to Dim.2 (44.59%) and Dim.3 (36.16%), while REM shows the largest effect in Dim.5 (33.38%).

**SUBJECT 3 – OBE1. Squared Cosine (Cos²) Values Indicating Condition Representation Across Principal Components**

| Condition | Dim.1 | Dim.2 | Dim.3 | Dim.4 | Dim.5 |
|---|---|---|---|---|---|
| OBE | 0.615 | 0.126 | 0.116 | 0.036 | 0.018 |
| REM | 0.372 | 0.232 | 0.144 | 0.043 | 0.067 |
| S1 | 0.250 | 0.417 | 0.174 | 0.028 | 0.026 |

| | | | | | |
|---|---|---|---|---|---|
| Wakefulness | 0.829 | 0.050 | 0.019 | 0.042 | 0.019 |

**Supplementary Table 51. OBE1. Squared Cosine ($Cos^2$) Values for Condition Representation Across Dimensions.** Wakefulness is best represented in Dim.1 ($Cos^2 = 0.829$), while S1 has the highest representation in Dim.2 ($Cos^2 = 0.417$). OBE shows strong representation in Dim.1 ($Cos^2 = 0.615$), and REM is moderately represented in Dim.2 ($Cos^2 = 0.232$).

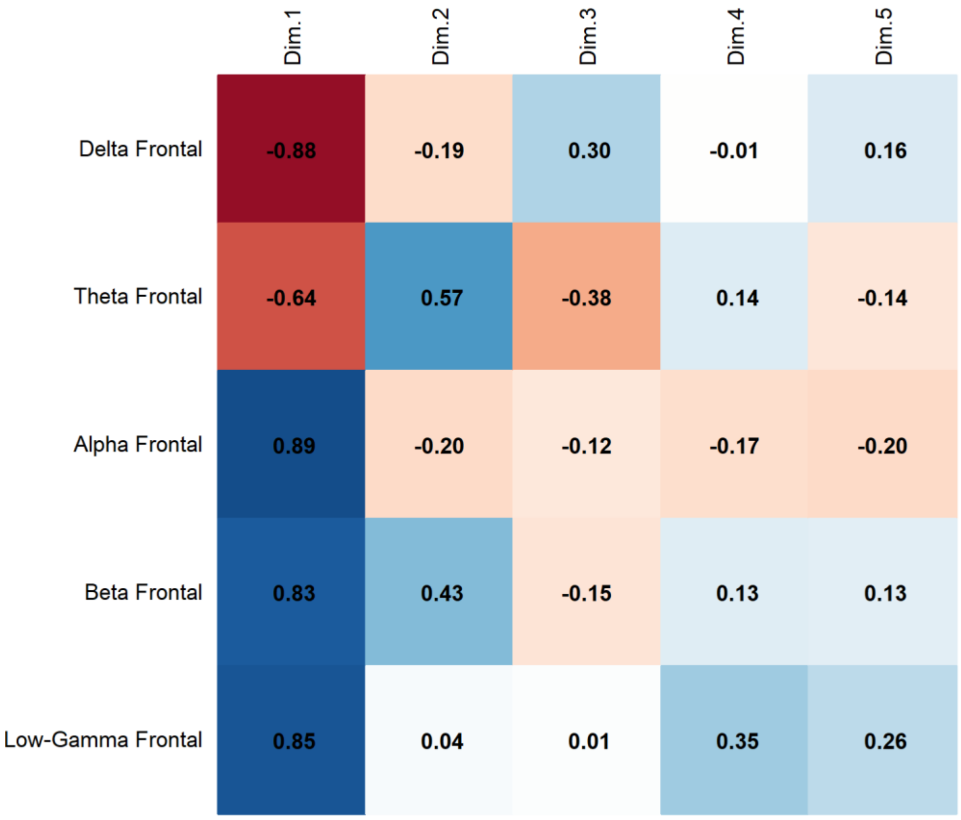


**Supplementary Fig. 21. OBE1**. **PCA Results**. Correlation of Frontal Band Activity with Principal Components revealed that Alpha activity presents a strong positive correlation with Dim.1 (0.89), while Delta (-0.88) and Theta (-0.64) correlate negatively with Dim.1. Beta (0.83) is associated with Dim.1, and Low-Gamma (0.85) with Dim.1.

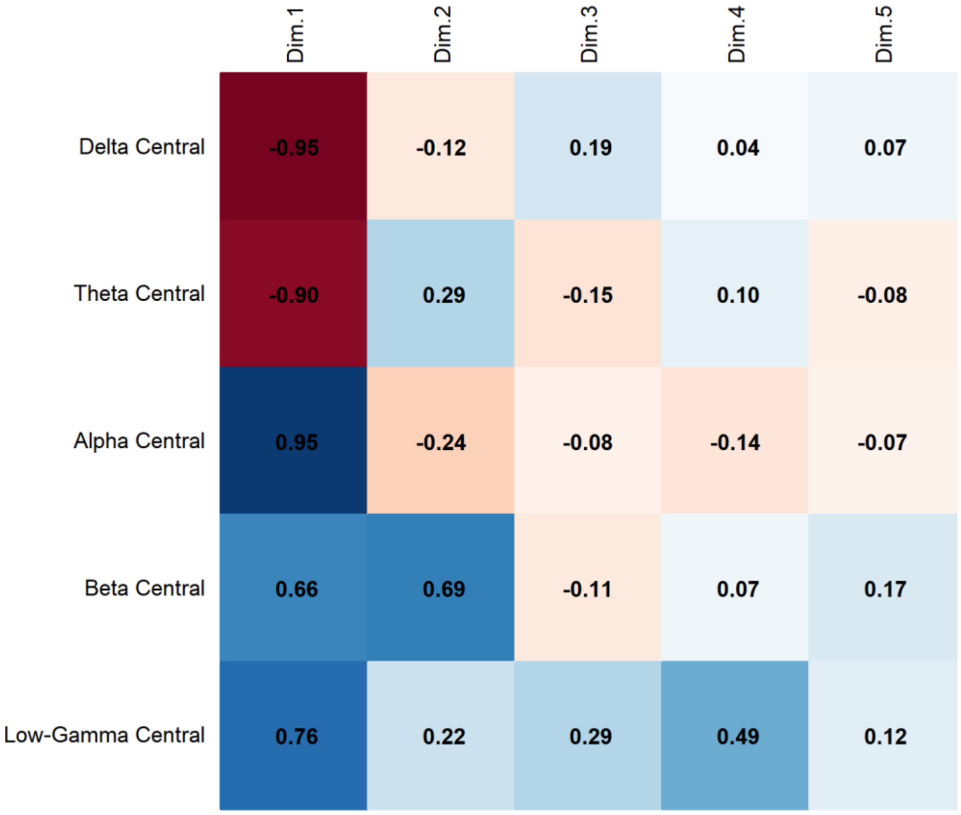


**Supplementary Fig. 22. OBE1. PCA Results**. Correlation of Central Band Activity with Principal Components revealed that Alpha activity presents a strong positive correlation with Dim.1 (0.95), while Delta (-0.95) and Theta (-0.90) correlate negatively with Dim.1. Beta (0.69) is associated with Dim.2, and Low-Gamma (0.76) with Dim.1.

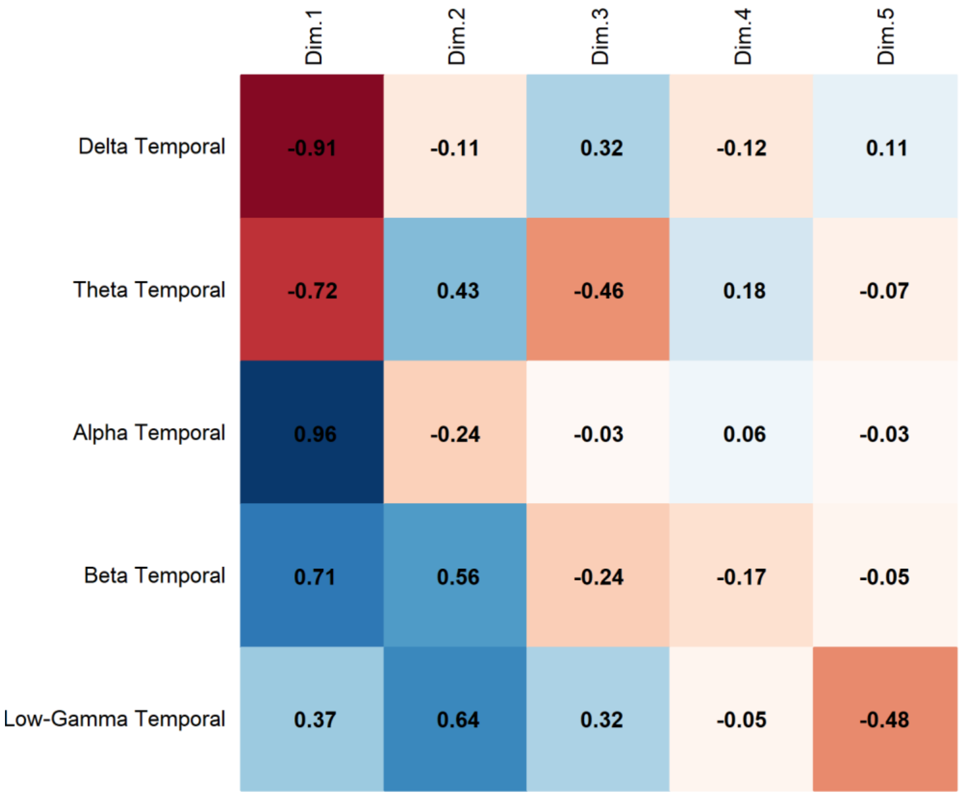


**Supplementary Fig. 23. OBE1**. **PCA Results**. Correlation of Temporal Band Activity with Principal Components revealed that Alpha activity presents a strong positive correlation with Dim.1 (0.96), while Delta (-0.91) and Theta (-0.72) correlate negatively with Dim.1. Beta (0.56) is associated with Dim.2, and Low-Gamma (0.64) with Dim.2.

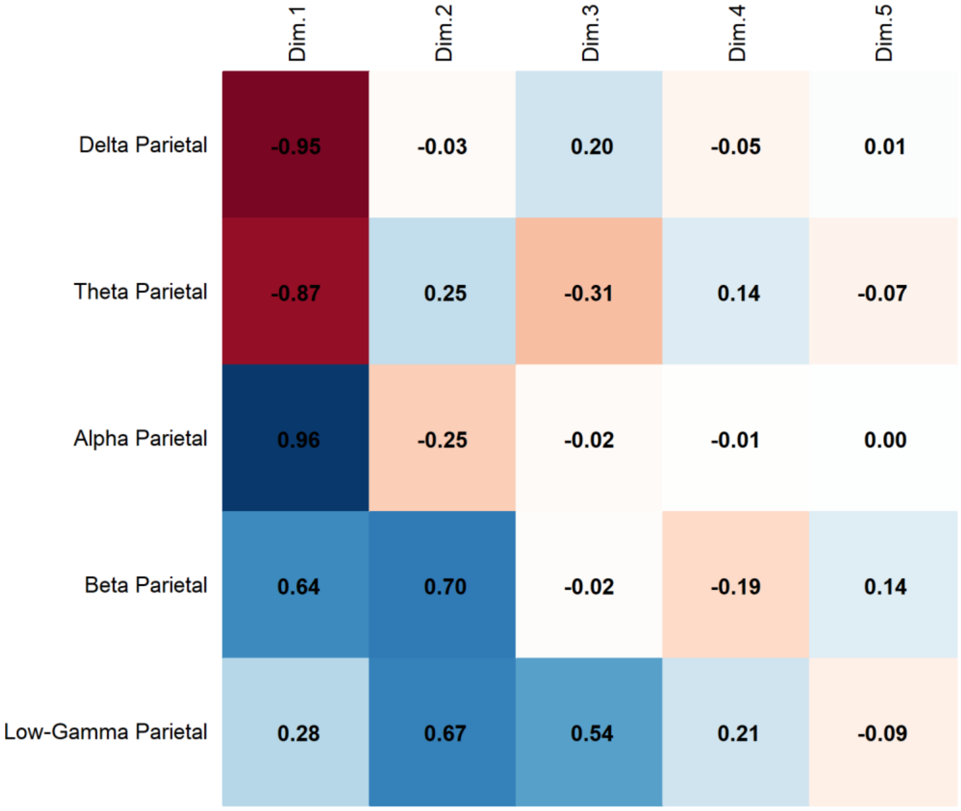


**Supplementary Fig. 24. OBE1. PCA Results**. Correlation of Parietal Band Activity with Principal Components revealed that Alpha activity presents a strong positive correlation with Dim.1 (0.96), while Delta (-0.95) and Theta (-0.87) correlate negatively with Dim.1. Beta (0.70) is associated with Dim.2, and Low-Gamma (0.67) with Dim.2.

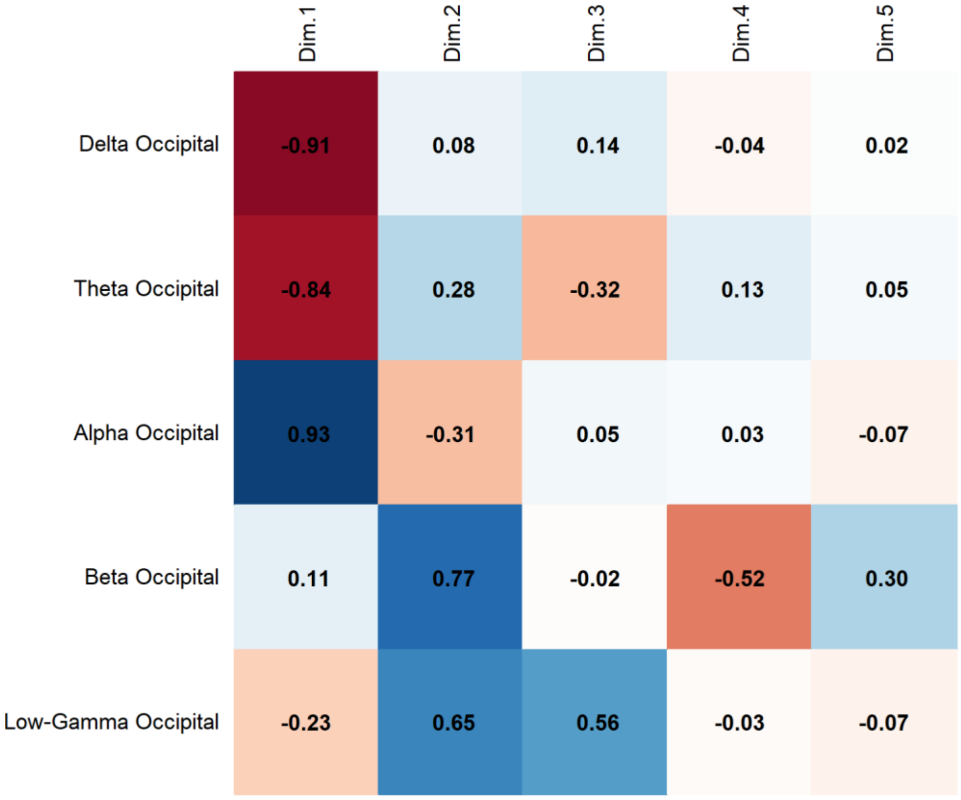


**Supplementary Fig. 25. OBE1. PCA Results**. Correlation of Occipital Band Activity with Principal Components revealed that Alpha activity presents a strong positive correlation with Dim.1 (0.93), while Delta (-0.91) and Theta (-0.84) correlate negatively with Dim.1. Beta (0.77) is associated with Dim.2, and Low-Gamma (0.65) with Dim.2.

**SUBJECT 3 – OBE2. PCA Results.**

| Component | Eigenvalue | Variance Explained (%) | Cumulative Variance (%) |
|---|---|---|---|
| 1 | 10.275 | 51.376 | 51.376 |
| 2 | 7.011 | 35.057 | 86.433 |
| 3 | 1.132 | 5.660 | 92.093 |
| 4 | 0.581 | 2.905 | 94.998 |
| 5 | 0.306 | 1.528 | 96.526 |
| 6 | 0.218 | 1.089 | 97.616 |
| 7 | 0.138 | 0.689 | 98.304 |
| 8 | 0.108 | 0.541 | 98.846 |
| 9 | 0.095 | 0.474 | 99.319 |

| | | | |
|---|---|---|---|
| 10 | 0.066 | 0.332 | 99.651 |
| 11 | 0.029 | 0.144 | 99.796 |
| 12 | 0.022 | 0.110 | 99.905 |
| 13 | 0.012 | 0.062 | 99.967 |
| 14 | 0.004 | 0.022 | 99.989 |
| 15 | 0.002 | 0.008 | 99.997 |
| 16 | 0.001 | 0.003 | 100.000 |
| 17 | 0.000 | 0.000 | 100.000 |
| 18 | 0.000 | 0.000 | 100.000 |
| 19 | 0.000 | 0.000 | 100.000 |
| 20 | 0.000 | 0.000 | 100.000 |

**Supplementary Table 52. OBE2. PCA Results.** Principal Component Analysis (PCA) shows that the first two components account for 86.43% of the total variance, with Component 1 explaining 51.38% and Component 2 explaining 35.06%. The first four components together capture 95.00% of the variance.

**SUBJECT 3 – OBE2. Contribution of Variables to the Principal Components (% Contribution)**

| Variable | Dim.1 | Dim.2 | Dim.3 | Dim.4 | Dim.5 |
|---|---|---|---|---|---|
| Delta Central | 8.356 | 1.053 | 2.118 | 0.022 | 0.639 |
| Theta Central | 7.518 | 0.025 | 12.049 | 2.573 | 6.373 |
| Alpha Central | 9.099 | 0.666 | 0.039 | 1.346 | 0.376 |
| Beta Central | 0.445 | 12.794 | 0.000 | 1.495 | 0.711 |
| Low-Gamma Central | 0.024 | 12.557 | 7.561 | 0.198 | 0.005 |
| Delta Frontal | 7.014 | 1.549 | 2.694 | 3.849 | 35.629 |
| Theta Frontal | 5.763 | 0.045 | 21.085 | 12.016 | 8.399 |
| Alpha Frontal | 8.355 | 0.541 | 0.229 | 3.148 | 20.717 |
| Beta Frontal | 1.218 | 10.076 | 0.468 | 22.335 | 0.008 |
| Low-Gamma Frontal | 0.254 | 11.570 | 7.434 | 8.141 | 2.003 |
| Delta Parietal | 6.630 | 0.849 | 17.212 | 1.230 | 9.405 |

| | | | | | |
|---|---|---|---|---|---|
| Theta Parietal | 8.476 | 0.001 | 3.897 | 1.398 | 1.105 |
| Alpha Parietal | 8.726 | 0.659 | 2.268 | 0.586 | 3.863 |
| Beta Parietal | 0.001 | 12.258 | 7.689 | 1.041 | 0.276 |
| Low-Gamma Parietal | 0.362 | 13.073 | 0.891 | 1.615 | 0.400 |
| Delta Occipital | 8.477 | 0.277 | 3.989 | 2.100 | 3.966 |
| Theta Occipital | 9.081 | 0.052 | 1.648 | 0.004 | 2.943 |
| Alpha Occipital | 8.831 | 0.487 | 0.083 | 4.312 | 3.182 |
| Beta Occipital | 0.328 | 10.662 | 8.360 | 16.303 | 0.001 |
| Low-Gamma Occipital | 1.043 | 10.807 | 0.284 | 16.287 | 0.000 |

**Supplementary Table 53. OBE2. Contribution of Variables to Principal Components.** The main contributors to Dim.1 are alpha occipital (8.83%), theta occipital (9.08%), and alpha central (9.10%). In Dim.2, the highest contributions come from beta central (12.79%), low-gamma parietal (13.07%), and low-gamma central (12.56%).

**SUBJECT 3 – OBE2. Squared Cosine ($Cos^2$) Values Indicating Variable Representation Across Principal Components.**

| Variable | Dim.1 | Dim.2 | Dim.3 | Dim.4 | Dim.5 |
|---|---|---|---|---|---|
| Delta Central | 0.859 | 0.074 | 0.024 | 0.000 | 0.002 |
| Theta Central | 0.772 | 0.002 | 0.136 | 0.015 | 0.019 |
| Alpha Central | 0.935 | 0.047 | 0.000 | 0.008 | 0.001 |
| Beta Central | 0.046 | 0.897 | 0.000 | 0.009 | 0.002 |
| Low-Gamma Central | 0.002 | 0.880 | 0.086 | 0.001 | 0.000 |
| Delta Frontal | 0.721 | 0.109 | 0.030 | 0.022 | 0.109 |
| Theta Frontal | 0.592 | 0.003 | 0.239 | 0.070 | 0.026 |
| Alpha Frontal | 0.858 | 0.038 | 0.003 | 0.018 | 0.063 |
| Beta Frontal | 0.125 | 0.706 | 0.005 | 0.130 | 0.000 |
| Low-Gamma Frontal | 0.026 | 0.811 | 0.084 | 0.047 | 0.006 |
| Delta Parietal | 0.681 | 0.060 | 0.195 | 0.007 | 0.029 |
| Theta Parietal | 0.871 | 0.000 | 0.044 | 0.008 | 0.003 |
| Alpha Parietal | 0.897 | 0.046 | 0.026 | 0.003 | 0.012 |

| | | | | | |
|---|---|---|---|---|---|
| Beta Parietal | 0.000 | 0.859 | 0.087 | 0.006 | 0.001 |
| Low-Gamma Parietal | 0.037 | 0.917 | 0.010 | 0.009 | 0.001 |
| Delta Occipital | 0.871 | 0.019 | 0.045 | 0.012 | 0.012 |
| Theta Occipital | 0.933 | 0.004 | 0.019 | 0.000 | 0.009 |
| Alpha Occipital | 0.907 | 0.034 | 0.001 | 0.025 | 0.010 |
| Beta Occipital | 0.034 | 0.748 | 0.095 | 0.095 | 0.000 |
| Low-Gamma Occipital | 0.107 | 0.758 | 0.003 | 0.095 | 0.000 |

**Supplementary Table 54. OBE2. Squared Cosine ($Cos^2$) Values for Variable Representation Across Dimensions**. The variables best represented in Dim.1 are alpha central ($Cos^2 = 0.935$), theta occipital ($Cos^2 = 0.933$), and alpha occipital ($Cos^2 = 0.907$). In Dim.2, the highest representations are found in beta central ($Cos^2 = 0.897$), low-gamma parietal ($Cos^2 = 0.917$), and beta parietal ($Cos^2 = 0.859$).

**SUBJECT 3 – OBE2. Contribution of Conditions to the Principal Components (% Contribution).**

| Condition | Dim.1 | Dim.2 | Dim.3 | Dim.4 | Dim.5 |
|---|---|---|---|---|---|
| OBE | 15.820 | 15.265 | 26.406 | 15.438 | 44.886 |
| REM | 16.580 | 7.643 | 36.469 | 16.271 | 19.938 |
| S1 | 4.178 | 69.770 | 31.217 | 63.072 | 6.636 |
| Wakefulness | 63.423 | 7.322 | 5.908 | 5.219 | 28.540 |

**Supplementary Table 55. OBE2. Contribution of Conditions to Principal Components**. Wakefulness contributes most to Dim.1 (63.42%), while S1 dominates Dim.2 (69.77%) and Dim.4 (63.07%). REM has the highest contribution to Dim.3 (36.47%), and OBE shows the largest effect in Dim.5 (44.89%).

**SUBJECT 3 – OBE2. Squared Cosine ($Cos^2$) Values Indicating Condition Representation Across Principal Components.**

| Condition | Dim.1 | Dim.2 | Dim.3 | Dim.4 | Dim.5 |
|---|---|---|---|---|---|
| OBE | 0.398 | 0.353 | 0.061 | 0.045 | 0.044 |
| REM | 0.542 | 0.158 | 0.185 | 0.032 | 0.020 |
| S1 | 0.120 | 0.679 | 0.060 | 0.077 | 0.005 |
| Wakefulness | 0.889 | 0.068 | 0.011 | 0.004 | 0.014 |

**Supplementary Table 56. OBE2. Squared Cosine ($Cos^2$) Values for Condition Representation Across Dimensions.** Wakefulness is best represented in Dim.1 ($Cos^2 = 0.889$), while S1 has the highest representation in Dim.2 ($Cos^2 = 0.679$). REM is moderately represented in Dim.1 ($Cos^2 = 0.542$) and Dim.3 ($Cos^2 = 0.185$), while OBE shows balanced contributions across Dim.1 ($Cos^2 = 0.398$) and Dim.2 ($Cos^2 = 0.353$).

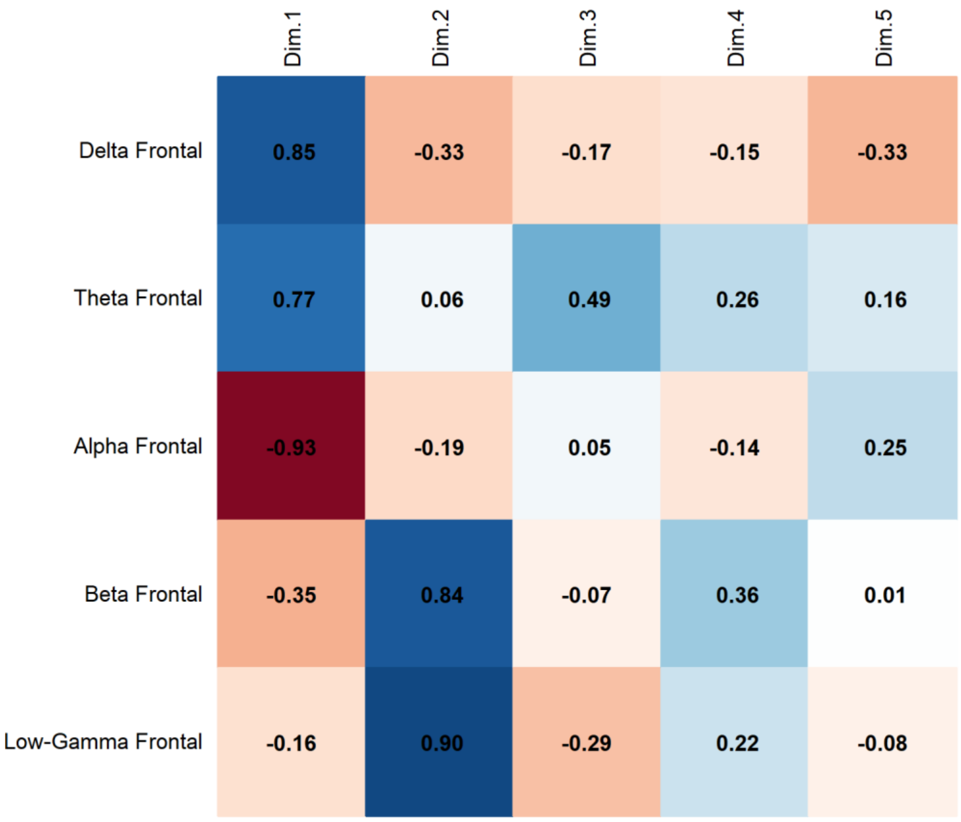


**Supplementary Fig. 26. OBE2**. **PCA Results**. Correlation of Frontal Band Activity with Principal Components revealed that Delta activity presents a strong positive correlation with Dim.1 (0.85), while Alpha (-0.93) and Theta (-0.77) correlate negatively with Dim.1. Beta (0.84) is associated with Dim.2, and Low-Gamma (0.90) with Dim.2.

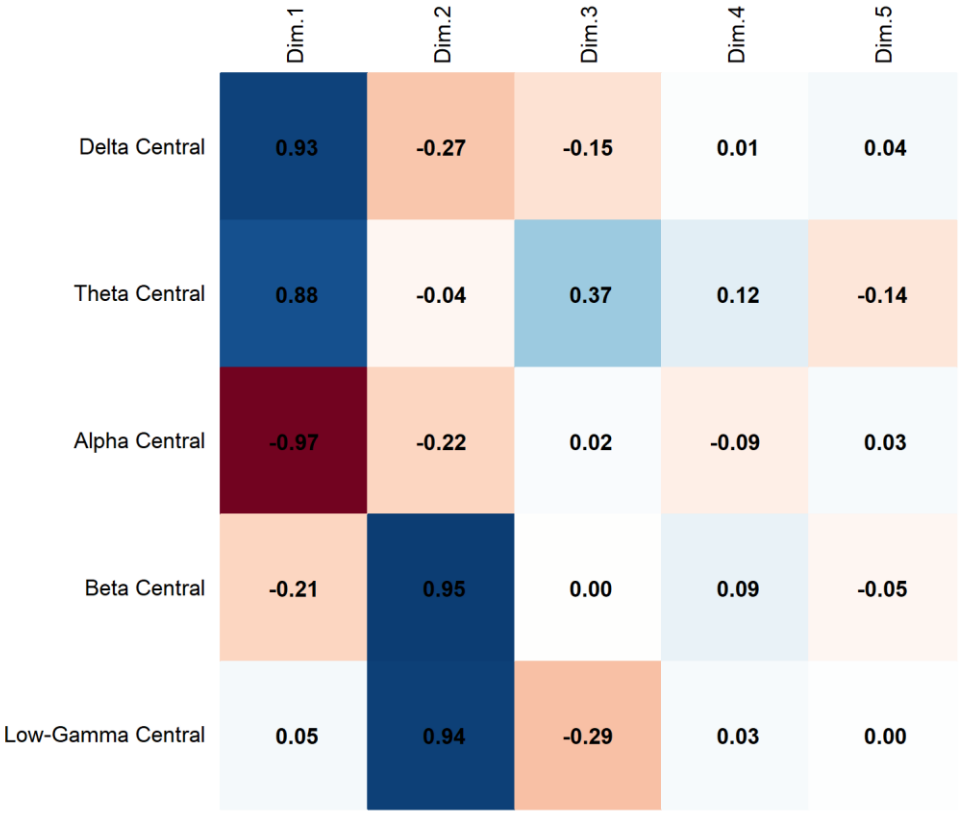


**Supplementary Fig. 27. OBE2**. **PCA Results**. Correlation of Central Band Activity with Principal Components revealed that Alpha activity presents a strong negative correlation with Dim.1 (-0.97), while Delta (0.93) and Theta (0.88) correlate positively with Dim.1. Beta (0.95) is associated with Dim.2, and Low-Gamma (0.94) with Dim.2.

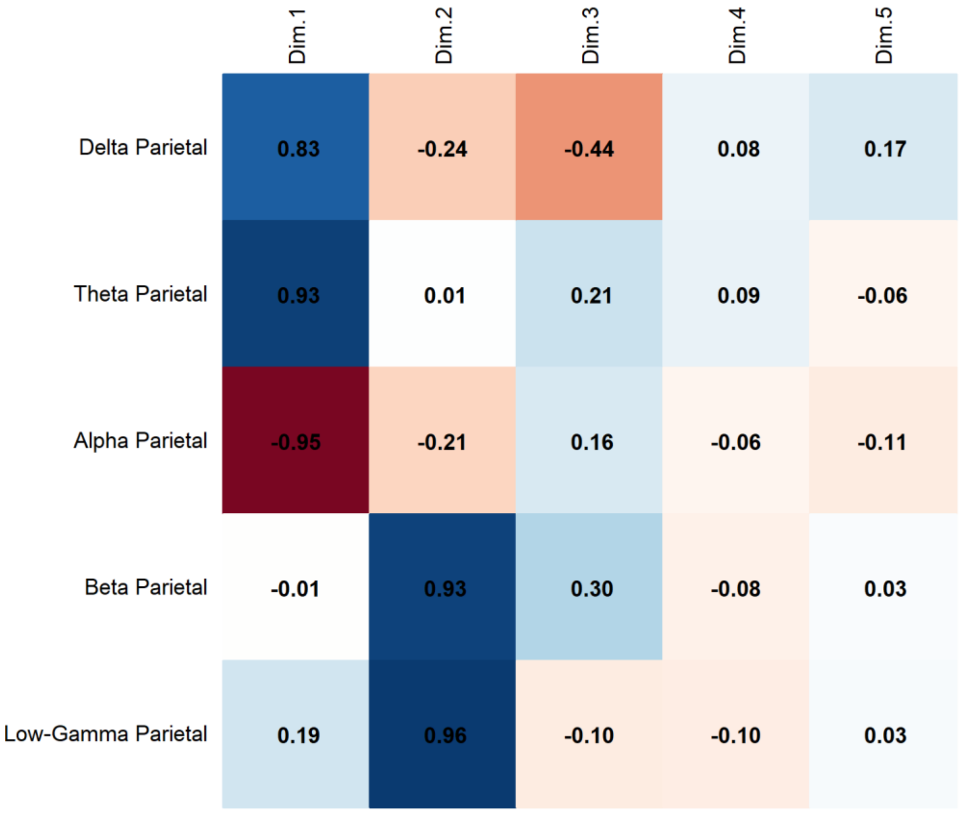


**Supplementary Fig. 28. OBE2. PCA Results**. Correlation of Parietal Band Activity with Principal Components revealed that Theta activity presents a strong positive correlation with Dim.1 (0.93), while Alpha (-0.95) and Delta (0.83) correlate negatively with Dim.1. Beta (0.93) is associated with Dim.2, and Low-Gamma (0.96) with Dim.2.

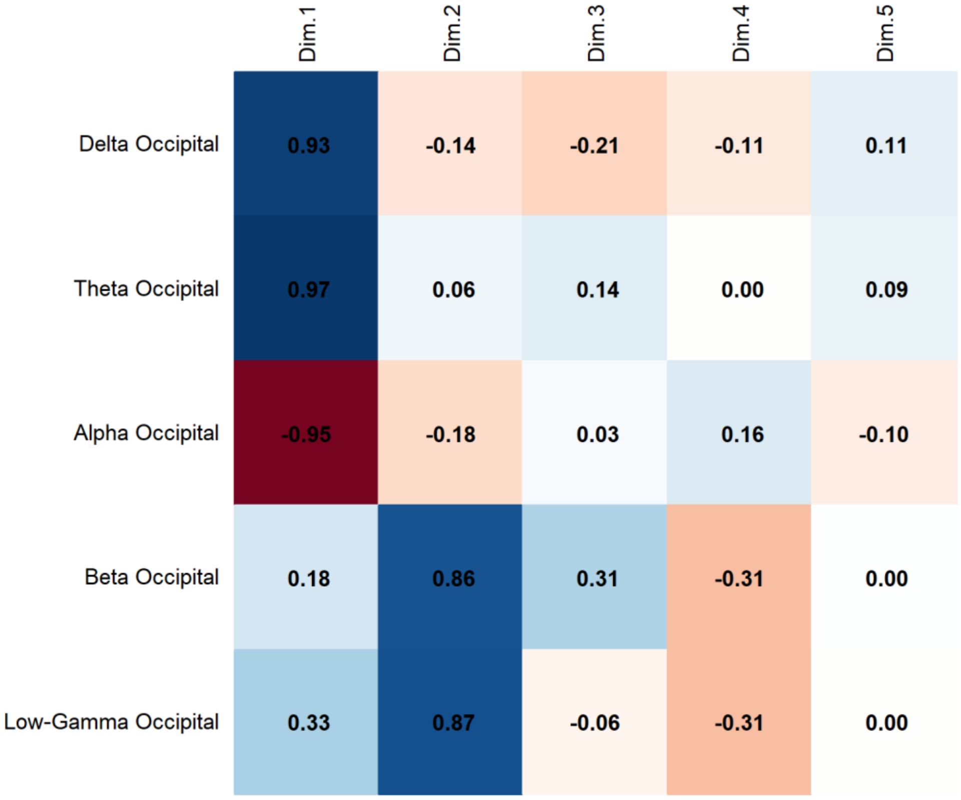


**Supplementary Fig. 29. OBE2. PCA Results**. Correlation of Occipital Band Activity with Principal Components revealed that Theta activity presents a strong positive correlation with Dim.1 (0.97), while Alpha (-0.95) and Delta (0.93) correlate negatively with Dim.1. Beta (0.86) is associated with Dim.2, and Low-Gamma (0.87) with Dim.2.

### PERMANOVA

**SUBJECT 3 - OBE1. PERMANOVA Results.**

| | df | Sum of Squares | $R^2$ | F-value | p-value |
|---|---|---|---|---|---|
| Model | 3 | 10.3187 | 0.88852 | 53.134 | 1e-04 *** |
| Residual | 20 | 1.2947 | 0.11148 | | |
| Dispersion (Homogeneity) | 3 | 0.002708 | | 0.0745 | 0.973 |
| Dispersion Residuals | 20 | 0.242259 | | | |

**Supplementary Table 57. OBE1. PERMANOVA Results**. The model explains 88.85% of the variance ($R^2 = 0.88852$) and is statistically significant ($p < 0.001$, Bonferroni-corrected). The homogeneity of dispersion test does not indicate significant differences in variance across groups ($p = 0.973$), confirming that the assumption of homogeneity is met.

**SUBJECT 3 – OBE1. Post-hoc PERMANOVA Results.**

| Comparison | Df | Sum Of Squares | F-value | $R^2$ | p-value | Adjusted p-value |
|---|---|---|---|---|---|---|
| OBE vs REM | 1 | 0.05088516 | 6.075 | 0.378 | 0.0081 | 0.049 |
| S1 vs OBE | 1 | 0.10175870 | 11.652 | 0.538 | 0.0049 | 0.029 |
| OBE vs Wakefulness | 1 | 1.17009704 | 178.139 | 0.947 | 0.0016 | 0.010 |
| REM vs Wakefulness | 1 | 0.82967842 | 110.759 | 0.917 | 0.0014 | 0.008 |
| S1 vs REM | 1 | 0.01544641 | 1.600 | 0.138 | 0.1573 | 0.944 |
| S1 vs Wakefulness | 1 | 0.84866454 | 108.134 | 0.915 | 0.0023 | 0.014 |

**Supplementary Table 58. OBE1. Post-hoc PERMANOVA Results.** The model assesses differences between conditions with Bonferroni-adjusted p-values.

**SUBJECT 3 – OBE1. Comparisons between OBE and REM**

| Frequency Band | Brain Region | F-value | $R^2$ | p-value |
|---|---|---|---|---|
| Delta | Frontal | 7.618 | 0.432 | 0.014 |
| | Central | 4.544 | 0.312 | 0.057 |
| | Temporal | 23.155 | 0.698 | 0.004 |
| | Parietal | 8.304 | 0.454 | 0.020 |
| | Occipital | 0.133 | 0.013 | 0.715 |
| Theta | Frontal | 2.409 | 0.194 | 0.157 |
| | Central | 0.201 | 0.020 | 0.649 |

| | | | | |
|---|---|---|---|---|
| | Temporal | 3.057 | 0.234 | 0.114 |
| | Parietal | 1.019 | 0.093 | 0.342 |
| | Occipital | 0.422 | 0.040 | 0.548 |
| Alpha | Frontal | 13.203 | 0.569 | 0.008 |
| | Central | 5.636 | 0.360 | 0.044 |
| | Temporal | 19.346 | 0.659 | 0.004 |
| | Parietal | 6.936 | 0.410 | 0.025 |
| | Occipital | 0.381 | 0.037 | 0.586 |
| Beta | Frontal | 12.239 | 0.550 | 0.005 |
| | Central | 12.580 | 0.557 | 0.005 |
| | Temporal | 24.777 | 0.712 | 0.002 |
| | Parietal | 4.170 | 0.294 | 0.066 |
| | Occipital | 1.573 | 0.136 | 0.235 |
| Low-Gamma | Frontal | 4.559 | 0.313 | 0.067 |
| | Central | 3.911 | 0.281 | 0.067 |
| | Temporal | 4.992 | 0.333 | 0.044 |
| | Parietal | 1.176 | 0.105 | 0.305 |
| | Occipital | 0.027 | 0.003 | 0.878 |

**Supplementary Table 59. OBE1. Comparisons between OBE and REM.** The model evaluates differences in spectral power across frequency bands and brain regions. Values are uncorrected for multiple comparisons.

**SUBJECT 3 – OBE1. Comparisons between S1 and OBE**

| Frequency Band | Brain Region | F-value | $R^2$ | p-value |
|---|---|---|---|---|
| Delta | Frontal | 36.669 | 0.786 | 0.002 |
| | Central | 10.002 | 0.500 | 0.013 |
| | Temporal | 45.653 | 0.820 | 0.002 |
| | Parietal | 12.719 | 0.560 | 0.008 |
| | Occipital | 1.736 | 0.148 | 0.218 |

| | | | | |
|---|---|---|---|---|
| Theta | Frontal | 22.606 | 0.693 | 0.006 |
| | Central | 0.054 | 0.005 | 0.831 |
| | Temporal | 18.038 | 0.643 | 0.005 |
| | Parietal | 3.173 | 0.241 | 0.108 |
| | Occipital | 1.590 | 0.137 | 0.239 |
| Alpha | Frontal | 11.775 | 0.541 | 0.011 |
| | Central | 4.096 | 0.291 | 0.073 |
| | Temporal | 13.415 | 0.573 | 0.012 |
| | Parietal | 3.247 | 0.245 | 0.104 |
| | Occipital | 0.391 | 0.038 | 0.541 |
| Beta | Frontal | 12.430 | 0.554 | 0.010 |
| | Central | 10.909 | 0.522 | 0.009 |
| | Temporal | 15.369 | 0.606 | 0.003 |
| | Parietal | 4.832 | 0.326 | 0.056 |
| | Occipital | 3.404 | 0.254 | 0.091 |
| Low-Gamma | Frontal | 8.119 | 0.448 | 0.019 |
| | Central | 2.522 | 0.201 | 0.138 |
| | Temporal | 1.541 | 0.134 | 0.240 |
| | Parietal | 0.327 | 0.032 | 0.577 |
| | Occipital | 0.016 | 0.002 | 0.912 |

**Supplementary Table 60. OBE1. Comparisons between S1 and OBE.** The model evaluates differences in spectral power across frequency bands and brain regions. Values are uncorrected for multiple comparisons.

**SUBJECT 3 – OBE1. Comparisons between OBE and Wakefulness**

| Frequency Band | Brain Region | F-value | $R^2$ | p-value |
|---|---|---|---|---|
| Delta | Frontal | 71.637 | 0.878 | 0.002 |
| | Central | 113.796 | 0.919 | 0.002 |

| | | | | |
|---|---|---|---|---|
| | Temporal | 223.682 | 0.957 | 0.003 |
| | Parietal | 136.905 | 0.932 | 0.002 |
| | Occipital | 59.562 | 0.856 | 0.002 |
| Theta | Frontal | 23.468 | 0.701 | 0.002 |
| | Central | 195.762 | 0.951 | 0.003 |
| | Temporal | 24.970 | 0.714 | 0.002 |
| | Parietal | 126.904 | 0.927 | 0.002 |
| | Occipital | 59.251 | 0.856 | 0.002 |
| Alpha | Frontal | 31.757 | 0.761 | 0.002 |
| | Central | 145.223 | 0.936 | 0.002 |
| | Temporal | 1,491.177 | 0.993 | 0.002 |
| | Parietal | 880.691 | 0.989 | 0.002 |
| | Occipital | 394.371 | 0.975 | 0.002 |
| Beta | Frontal | 107.931 | 0.915 | 0.002 |
| | Central | 61.049 | 0.859 | 0.002 |
| | Temporal | 143.015 | 0.935 | 0.003 |
| | Parietal | 31.544 | 0.759 | 0.003 |
| | Occipital | 0.576 | 0.054 | 0.459 |
| Low-Gamma | Frontal | 24.852 | 0.713 | 0.003 |
| | Central | 9.800 | 0.495 | 0.002 |
| | Temporal | 9.899 | 0.497 | 0.013 |
| | Parietal | 0.416 | 0.040 | 0.707 |
| | Occipital | 2.745 | 0.215 | 0.137 |

**Supplementary Table 61. OBE1. Comparisons between OBE and Wakefulness.** The model evaluates differences in spectral power across frequency bands and brain regions. Values are uncorrected for multiple comparisons.

**SUBJECT 3 – OBE2. PERMANOVA Results.**

| | df | Sum of Squares | R² | F-value | p-value |
|---|---|---|---|---|---|
| Model | 3 | 6.8626 | 0.83226 | 33.078 | 1e-04 *** |
| Residual | 20 | 1.3831 | 0.16774 | | |
| Dispersion (Homogeneity) | 3 | 0.095954 | | 2.7046 | 0.07277 |
| Dispersion Residuals | 20 | 0.236516 | | | |

**Supplementary Table 62. OBE2. PERMANOVA Results.** The model explains 83.23% of the variance ($R^2 = 0.83226$) and is statistically significant ($p < 0.001$, Bonferroni-corrected). The homogeneity of dispersion test does not indicate significant differences in variance across groups ($p = 0.07277$), confirming that the assumption of homogeneity is met.

**SUBJECT 3 – OBE2. Post-hoc PERMANOVA Results.**

| Comparison | Df | Sum of Squares | F-value | R² | p-value | Adjusted p-value |
|---|---|---|---|---|---|---|
| OBE vs REM | 1 | 0.02035176 | 1.524 | 0.132 | 0.2017 | 1.000 |
| S1 vs OBE | 1 | 0.17418382 | 11.616 | 0.537 | 0.0028 | 0.017 |
| OBE vs Wakefulness | 1 | 0.77911952 | 74.655 | 0.882 | 0.0022 | 0.013 |
| REM vs Wakefulness | 1 | 0.79833762 | 116.681 | 0.921 | 0.0022 | 0.013 |
| S1 vs REM | 1 | 0.13606850 | 11.935 | 0.544 | 0.0038 | 0.023 |
| S1 vs Wakefulness | 1 | 0.57448083 | 67.746 | 0.871 | 0.0020 | 0.012 |

**Supplementary Table 63. OBE2. Post-hoc PERMANOVA Results.** The model assesses differences between conditions with Bonferroni-adjusted p-values.

**SUBJECT 3 – OBE2. Comparisons between OBE and REM**

| Frequency Band | Brain Region | F-value | R² | p-value |
|---|---|---|---|---|
| Delta | Frontal | 0.213 | 0.021 | 0.651 |
| | Central | 1.466 | 0.128 | 0.239 |
| | Parietal | 4.567 | 0.314 | 0.074 |
| | Occipital | 0.014 | 0.001 | 0.887 |
| Theta | Frontal | 0.106 | 0.011 | 0.739 |
| | Central | 5.806 | 0.367 | 0.037 |
| | Parietal | 1.918 | 0.161 | 0.210 |
| | Occipital | 2.584 | 0.205 | 0.162 |

| | | | | |
|---|---|---|---|---|
| Alpha | Frontal | 2.062 | 0.171 | 0.181 |
| | Central | 0.002 | 0.000 | 0.970 |
| | Parietal | 2.102 | 0.174 | 0.168 |
| | Occipital | 1.700 | 0.145 | 0.287 |
| Beta | Frontal | 0.206 | 0.020 | 0.613 |
| | Central | 0.210 | 0.021 | 0.653 |
| | Parietal | 1.419 | 0.124 | 0.267 |
| | Occipital | 8.928 | 0.472 | 0.016 |
| Low-Gamma | Frontal | 0.506 | 0.048 | 0.506 |
| | Central | 0.419 | 0.040 | 0.534 |
| | Parietal | 0.836 | 0.077 | 0.373 |
| | Occipital | 20.325 | 0.670 | 0.002 |

**Supplementary Table 64. OBE2. Comparisons between OBE and REM.** The model evaluates differences in spectral power across frequency bands and brain regions. Values are uncorrected for multiple comparisons.

**SUBJECT 3 – OBE2. Comparisons between S1 and OBE**

| Frequency Band | Brain Region | F-value | $R^2$ | p-value |
|---|---|---|---|---|
| Delta | Frontal | 33.756 | 0.771 | 0.002 |
| | Central | 38.790 | 0.795 | 0.002 |
| | Parietal | 11.258 | 0.530 | 0.004 |
| | Occipital | 4.495 | 0.310 | 0.086 |
| Theta | Frontal | 0.320 | 0.031 | 0.622 |
| | Central | 0.274 | 0.027 | 0.611 |
| | Parietal | 0.720 | 0.067 | 0.412 |
| | Occipital | 0.611 | 0.058 | 0.503 |
| Alpha | Frontal | 1.722 | 0.147 | 0.230 |
| | Central | 1.761 | 0.150 | 0.221 |
| | Parietal | 1.009 | 0.092 | 0.322 |

| | | | | |
|---|---|---|---|---|
| | Occipital | 0.009 | 0.001 | 0.921 |
| Beta | Frontal | 25.342 | 0.717 | 0.003 |
| | Central | 39.200 | 0.797 | 0.002 |
| | Parietal | 24.757 | 0.712 | 0.004 |
| | Occipital | 27.764 | 0.735 | 0.004 |
| Low-Gamma | Frontal | 23.013 | 0.697 | 0.002 |
| | Central | 14.016 | 0.584 | 0.003 |
| | Parietal | 16.520 | 0.623 | 0.003 |
| | Occipital | 17.558 | 0.637 | 0.002 |

**Supplementary Table 65. OBE2. Comparisons between S1 and OBE**. The model evaluates differences in spectral power across frequency bands and brain regions. Values are uncorrected for multiple comparisons.

**SUBJECT 3 – OBE2. Comparisons between OBE and Wakefulness**

| Frequency Band | Brain Region | F-value | $R^2$ | p-value |
|---|---|---|---|---|
| Delta | Frontal | 40.785 | 0.803 | 0.002 |
| | Central | 124.782 | 0.926 | 0.002 |
| | Parietal | 33.033 | 0.768 | 0.002 |
| | Occipital | 28.273 | 0.739 | 0.003 |
| Theta | Frontal | 132.186 | 0.930 | 0.003 |
| | Central | 66.980 | 0.870 | 0.002 |
| | Parietal | 343.443 | 0.972 | 0.002 |
| | Occipital | 74.515 | 0.882 | 0.002 |
| Alpha | Frontal | 51.762 | 0.838 | 0.002 |
| | Central | 305.430 | 0.968 | 0.002 |
| | Parietal | 98.602 | 0.908 | 0.002 |
| | Occipital | 35.530 | 0.780 | 0.002 |
| Beta | Frontal | 10.630 | 0.515 | 0.010 |
| | Central | 6.784 | 0.404 | 0.032 |

| | | | | |
|---|---|---|---|---|
| | Parietal | 0.325 | 0.031 | 0.634 |
| | Occipital | 0.351 | 0.034 | 0.607 |
| Low-Gamma | Frontal | 3.022 | 0.232 | 0.068 |
| | Central | 1.570 | 0.136 | 0.277 |
| | Parietal | 4.815 | 0.325 | 0.005 |
| | Occipital | 14.320 | 0.589 | 0.006 |

**Supplementary Table 66. OBE2.** Comparisons between OBE and Wakefulness. The model evaluates differences in spectral power across frequency bands and brain regions. Values are uncorrected for multiple comparisons.

### False Awakening

We present the Principal Component Analysis (PCA) and PERMANOVA results for Subjects 3, 6, and 7 in the False Awakening (FA) condition. The PCA results are shown first, followed by the PERMANOVA results.

### PCA results.

**SUBJECT 3 – FA. PCA Results.**

| Component | Eigenvalue | Variance Explained (%) | Cumulative Variance (%) |
|---|---|---|---|
| 1 | 14.587 | 58.349 | 58.349 |
| 2 | 4.738 | 18.952 | 77.301 |
| 3 | 2.446 | 9.783 | 87.084 |
| 4 | 1.243 | 4.972 | 92.056 |
| 5 | 0.471 | 1.883 | 93.939 |
| 6 | 0.412 | 1.650 | 95.588 |
| 7 | 0.292 | 1.166 | 96.754 |
| 8 | 0.262 | 1.046 | 97.801 |
| 9 | 0.197 | 0.789 | 98.589 |
| 10 | 0.106 | 0.424 | 99.013 |
| 11 | 0.078 | 0.312 | 99.325 |
| 12 | 0.047 | 0.188 | 99.513 |
| 13 | 0.033 | 0.131 | 99.644 |
| 14 | 0.031 | 0.122 | 99.766 |
| 15 | 0.023 | 0.090 | 99.856 |
| 16 | 0.018 | 0.070 | 99.926 |
| 17 | 0.011 | 0.044 | 99.970 |
| 18 | 0.004 | 0.015 | 99.985 |
| 19 | 0.003 | 0.012 | 99.997 |
| 20 | 0.001 | 0.003 | 100.000 |
| 21 | 0.000 | 0.000 | 100.000 |
| 22 | 0.000 | 0.000 | 100.000 |

| | | | |
|---|---|---|---|
| 23 | 0.000 | 0.000 | 100.000 |

**Supplementary Table 67. False Awakening. PCA Results**. Principal Component Analysis (PCA) shows that the first two components account for 77.30% of the total variance, with Component 1 explaining 58.35% and Component 2 explaining 18.95%. The first four components together capture 92.06% of the variance.

**SUBJECT 3 – FA. Contribution of Variables to the Principal Components (% Contribution).**

| Variables | Dim.1 | Dim.2 | Dim.3 | Dim.4 | Dim.5 |
|---|---|---|---|---|---|
| Delta Central | 5.65 | 1.77 | 1.32 | 1.24 | 3.96 |
| Theta Central | 2.17 | 11.13 | 3.25 | 0.00 | 3.08 |
| Alpha Central | 5.48 | 0.94 | 0.07 | 8.06 | 5.94 |
| Beta Central | 5.34 | 1.13 | 0.14 | 7.93 | 4.56 |
| Low-Gamma Central | 4.03 | 0.54 | 4.49 | 14.26 | 2.06 |
| Delta Frontal | 5.22 | 3.09 | 1.79 | 1.47 | 1.39 |
| Theta Frontal | 0.70 | 13.35 | 6.86 | 0.20 | 6.05 |
| Alpha Frontal | 4.44 | 0.31 | 1.42 | 21.01 | 1.01 |
| Beta Frontal | 4.90 | 0.20 | 2.94 | 9.53 | 10.19 |
| Low-Gamma Frontal | 5.04 | 0.01 | 0.01 | 11.41 | 8.81 |
| Delta Temporal | 5.41 | 1.92 | 3.02 | 1.11 | 0.82 |
| Theta Temporal | 1.73 | 12.08 | 5.54 | 0.15 | 0.12 |
| Alpha Temporal | 6.06 | 0.70 | 0.48 | 2.34 | 0.45 |
| Beta Temporal | 4.16 | 5.98 | 0.90 | 1.55 | 1.15 |
| Low-Gamma Temporal | 4.54 | 0.88 | 3.25 | 3.38 | 11.64 |
| Delta Parietal | 6.09 | 0.44 | 2.01 | 0.47 | 0.40 |
| Theta Parietal | 3.34 | 8.19 | 1.08 | 0.56 | 0.76 |
| Alpha Parietal | 5.86 | 1.33 | 1.99 | 0.08 | 0.01 |
| Beta Parietal | 3.45 | 6.06 | 4.69 | 0.04 | 0.46 |
| Low-Gamma Parietal | 0.73 | 6.09 | 21.39 | 0.99 | 1.75 |
| Delta Occipital | 5.32 | 0.19 | 4.05 | 2.16 | 10.33 |
| Theta Occipital | 3.28 | 5.97 | 3.68 | 0.05 | 6.66 |

| | | | | | |
|---|---|---|---|---|---|
| Alpha Occipital | 5.63 | 1.41 | 1.68 | 0.29 | 12.26 |
| Beta Occipital | 1.18 | 7.43 | 8.82 | 8.89 | 0.49 |
| Low-Gamma Occipital | 0.26 | 8.87 | 15.11 | 2.85 | 5.65 |

**Supplementary Table 68. False Awakening. Contribution of Variables to Principal Components.** The main contributors to Dim.1 are alpha temporal (6.06%), delta parietal (6.09%), and delta central (5.65%). In Dim.2, the highest contributions come from theta frontal (13.35%), theta central (11.13%), and low-gamma occipital (8.87%).

**SUBJECT 3 – FA. Squared Cosine (Cos²) Values Indicating Variable Representation Across Dimensions**

| Variables | Dim.1 | Dim.2 | Dim.3 | Dim.4 | Dim.5 |
|---|---|---|---|---|---|
| Delta Central | 0.82 | 0.08 | 0.03 | 0.02 | 0.02 |
| Theta Central | 0.32 | 0.53 | 0.08 | 0.00 | 0.01 |
| Alpha Central | 0.80 | 0.04 | 0.00 | 0.10 | 0.03 |
| Beta Central | 0.78 | 0.05 | 0.00 | 0.10 | 0.02 |
| Low-Gamma Central | 0.59 | 0.03 | 0.11 | 0.18 | 0.01 |
| Delta Frontal | 0.76 | 0.15 | 0.04 | 0.02 | 0.01 |
| Theta Frontal | 0.10 | 0.63 | 0.17 | 0.00 | 0.03 |
| Alpha Frontal | 0.65 | 0.01 | 0.03 | 0.26 | 0.00 |
| Beta Frontal | 0.72 | 0.01 | 0.07 | 0.12 | 0.05 |
| Low-Gamma Frontal | 0.73 | 0.00 | 0.00 | 0.14 | 0.04 |
| Delta Temporal | 0.79 | 0.09 | 0.07 | 0.01 | 0.00 |
| Theta Temporal | 0.25 | 0.57 | 0.14 | 0.00 | 0.00 |
| Alpha Temporal | 0.88 | 0.03 | 0.01 | 0.03 | 0.00 |
| Beta Temporal | 0.61 | 0.28 | 0.02 | 0.02 | 0.01 |
| Low-Gamma Temporal | 0.66 | 0.04 | 0.08 | 0.04 | 0.05 |
| Delta Parietal | 0.89 | 0.02 | 0.05 | 0.01 | 0.00 |
| Theta Parietal | 0.49 | 0.39 | 0.03 | 0.01 | 0.00 |
| Alpha Parietal | 0.85 | 0.06 | 0.05 | 0.00 | 0.00 |

| | | | | | |
|---|---|---|---|---|---|
| Beta Parietal | 0.50 | 0.29 | 0.11 | 0.00 | 0.00 |
| Low-Gamma Parietal | 0.11 | 0.29 | 0.52 | 0.01 | 0.01 |
| Delta Occipital | 0.78 | 0.01 | 0.10 | 0.03 | 0.05 |
| Theta Occipital | 0.48 | 0.28 | 0.09 | 0.00 | 0.03 |
| Alpha Occipital | 0.82 | 0.07 | 0.04 | 0.00 | 0.06 |
| Beta Occipital | 0.17 | 0.35 | 0.22 | 0.11 | 0.00 |
| Low-Gamma Occipital | 0.04 | 0.42 | 0.37 | 0.04 | 0.03 |

**Supplementary Table 69. False Awakening. Squared Cosine ($Cos^2$) Values for Variable Representation Across Dimensions.** The variables best represented in Dim.1 are delta parietal ($Cos^2 = 0.89$), alpha temporal ($Cos^2 = 0.88$), and delta central ($Cos^2 = 0.82$). In Dim.2, the highest representations are found in theta frontal ($Cos^2 = 0.63$), theta temporal ($Cos^2 = 0.57$), and low-gamma occipital ($Cos^2 = 0.42$).

**SUBJECT 3 – FA. Contribution of Conditions to the Principal Components (% Contribution)**

| Condition | Dim.1 | Dim.2 | Dim.3 | Dim.4 | Dim.5 |
|---|---|---|---|---|---|
| FA | 11.971 | 35.685 | 46.188 | 13.185 | 35.039 |
| REM | 34.473 | 34.063 | 13.137 | 3.073 | 13.230 |
| S1 | 1.854 | 17.299 | 22.941 | 32.085 | 43.172 |
| Wakefulness | 51.702 | 12.953 | 17.735 | 51.657 | 8.559 |

**Supplementary Table 70. False Awakening. Contribution of Conditions to Principal Components.** Wakefulness contributes most to Dim.1 (51.70%) and Dim.4 (51.66%). False Awakening (FA) dominates Dim.3 (46.19%) and has a strong contribution in Dim.2 (35.69%). S1 shows the highest effect in Dim.5 (43.17%), while REM is evenly distributed across Dim.1 (34.47%) and Dim.2 (34.06%).

**SUBJECT 3 – FA. Squared Cosine ($Cos^2$) Values Indicating Condition Representation Across Dimensions**

| Condition | Dim.1 | Dim.2 | Dim.3 | Dim.4 | Dim.5 |
|---|---|---|---|---|---|
| FA | 0.235 | 0.322 | 0.199 | 0.042 | 0.051 |
| REM | 0.680 | 0.169 | 0.055 | 0.006 | 0.010 |
| S1 | 0.131 | 0.315 | 0.219 | 0.126 | 0.081 |
| Wakefulness | 0.732 | 0.055 | 0.048 | 0.099 | 0.004 |

**Supplementary Table 71. False Awakening. Squared Cosine ($Cos^2$) Values for Condition Representation Across Dimensions.** Wakefulness is best represented in Dim.1 ($Cos^2 = 0.732$), while FA has the highest representation in Dim.2 ($Cos^2 = 0.322$). S1 is moderately represented in both Dim.2 ($Cos^2 = 0.315$) and Dim.3 ($Cos^2 = 0.219$), while REM is most strongly represented in Dim.1 ($Cos^2 = 0.680$).

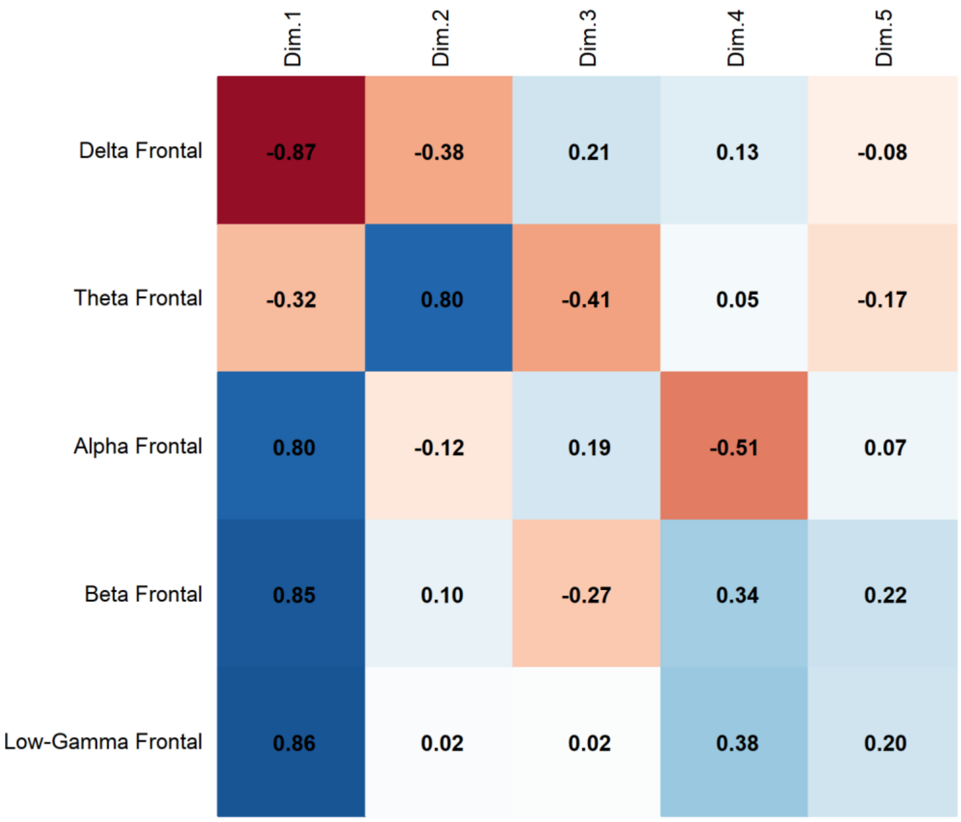


**Supplementary Fig. 30. False Awakening**, **PCA Results**. Correlation of Frontal Band Activity with Principal Components revealed that Beta activity presents a strong positive correlation with Dim.1 (0.85), while Delta (-0.87) and Theta (-0.32) correlate negatively with Dim.1. Alpha (0.80) is associated with Dim.1, and Low-Gamma (0.86) with Dim.1.

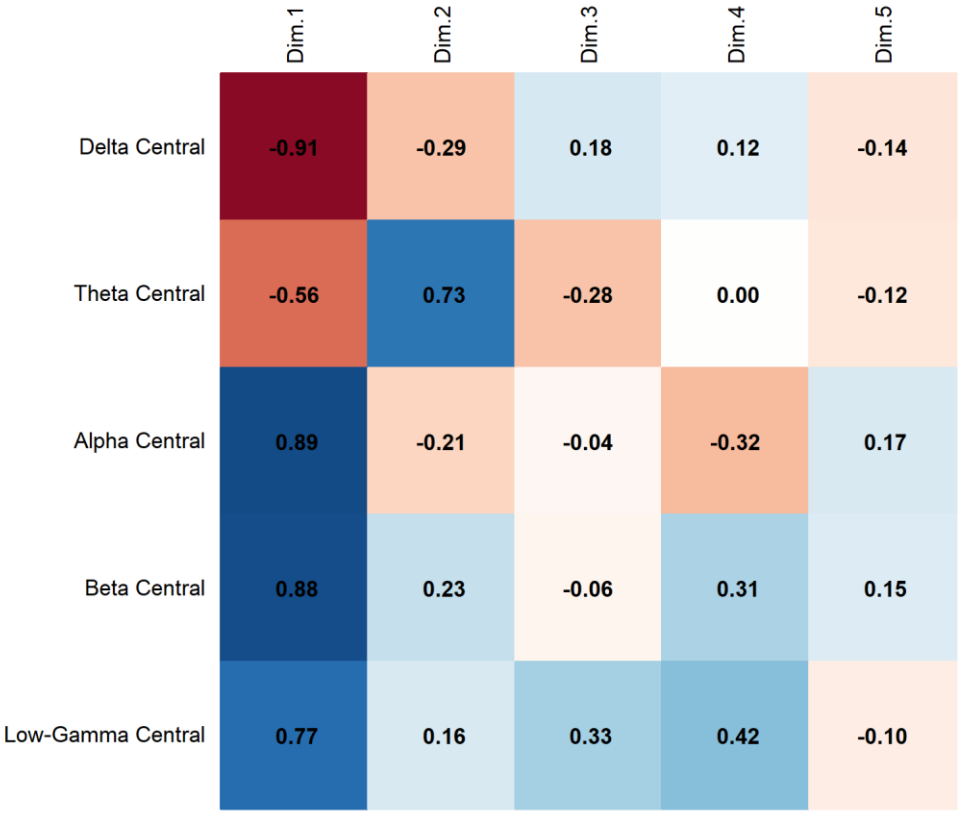


**Supplementary Fig. 31. False Awakening**. **PCA Results**. Correlation of Central Band Activity with Principal Components revealed that Beta activity presents a strong positive correlation with Dim.1 (0.88), while Delta (-0.91) and Theta (-0.56) correlate negatively with Dim.1. Alpha (0.89) is associated with Dim.1, and Low-Gamma (0.77) with Dim.1.

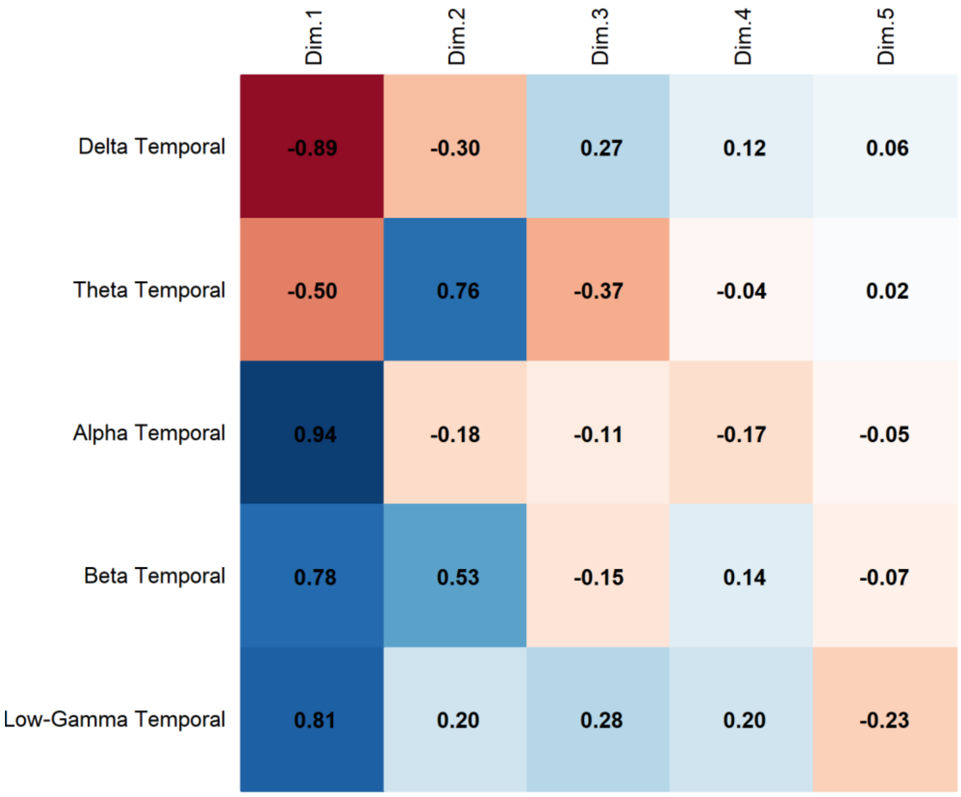


**Supplementary Fig. 32. False Awakening**, **PCA Results**. Correlation of Temporal Band Activity with Principal Components revealed that Alpha activity presents a strong positive correlation with Dim.1 (0.94), while Delta (-0.89) and Theta (-0.50) correlate negatively with Dim.1. Beta (0.78) is associated with Dim.1, and Low-Gamma (0.81) with Dim.1.

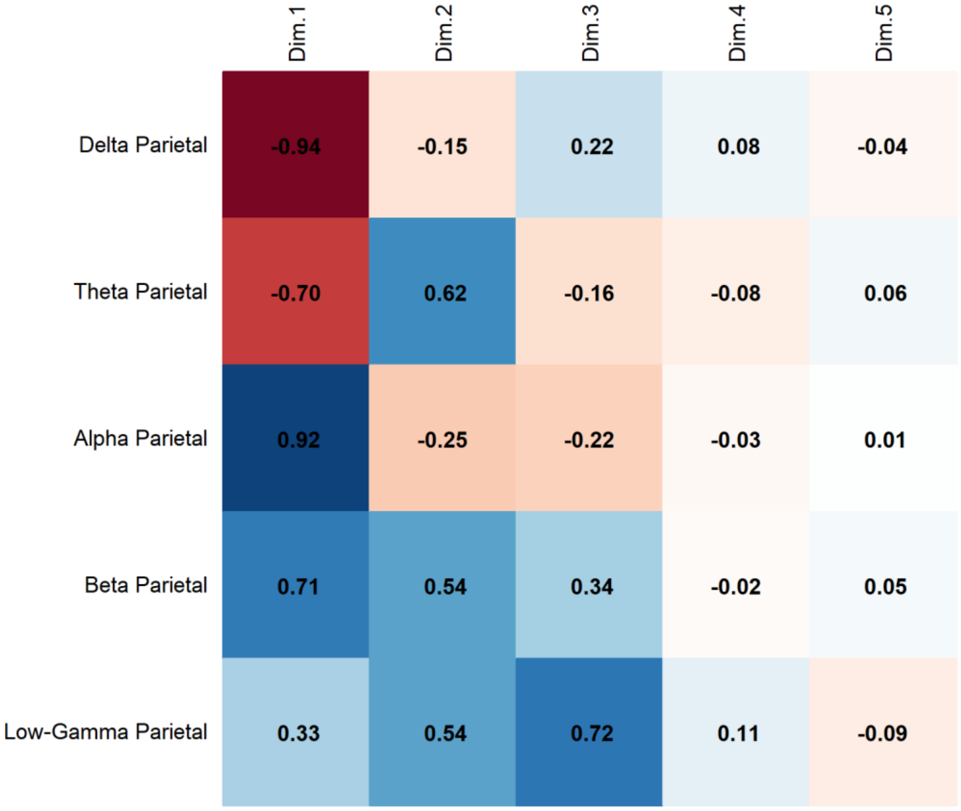


**Supplementary Fig. 33. False Awakening**, **PCA Results**. Correlation of Parietal Band Activity with Principal Components revealed that Alpha activity presents a strong positive correlation with Dim.1 (0.92), while Delta (-0.94) and Theta (-0.70) correlate negatively with Dim.1. Beta (0.71) is associated with Dim.1, and Low-Gamma (0.72) with Dim.3.

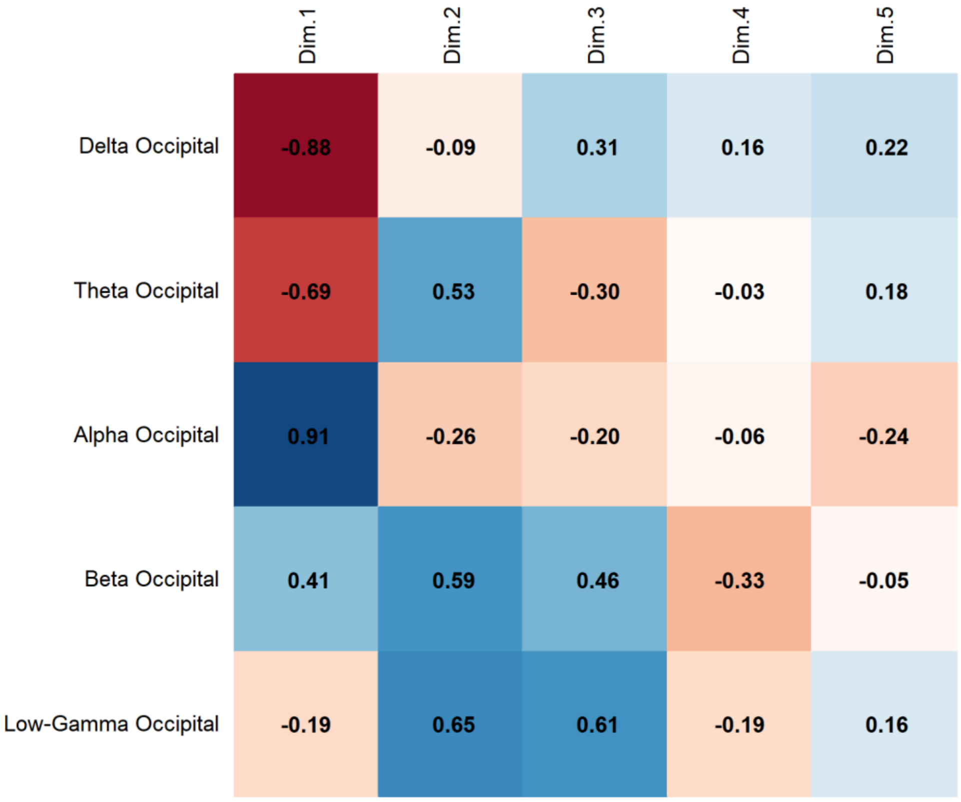


**Supplementary Fig. 34. False Awakening**, **PCA Results**. Correlation of Occipital Band Activity with Principal Components revealed that Alpha activity presents a strong positive correlation with Dim.1 (0.91), while Delta (-0.88) and Theta (-0.69) correlate negatively with Dim.1. Beta (0.59) is associated with Dim.2, and Low-Gamma (0.65) with Dim.2.

**SUBJECT 6. PCA Results.**

| Component | Eigenvalue | Variance Explained (%) | Cumulative Variance (%) |
|---|---|---|---|
| 1 | 11.846 | 47.385 | 47.385 |
| 2 | 6.908 | 27.632 | 75.017 |
| 3 | 2.735 | 10.940 | 85.957 |
| 4 | 1.367 | 5.469 | 91.426 |
| 5 | 0.984 | 3.935 | 95.361 |
| 6 | 0.379 | 1.516 | 96.876 |
| 7 | 0.202 | 0.808 | 97.684 |
| 8 | 0.143 | 0.572 | 98.256 |
| 9 | 0.127 | 0.508 | 98.763 |

| | | | |
|---|---|---|---|
| 10 | 0.106 | 0.424 | 99.187 |
| 11 | 0.057 | 0.229 | 99.416 |
| 12 | 0.043 | 0.170 | 99.586 |
| 13 | 0.032 | 0.130 | 99.716 |
| 14 | 0.026 | 0.104 | 99.820 |
| 15 | 0.022 | 0.089 | 99.909 |
| 16 | 0.016 | 0.063 | 99.971 |
| 17 | 0.004 | 0.017 | 99.989 |
| 18 | 0.002 | 0.008 | 99.997 |
| 19 | 0.001 | 0.002 | 99.999 |
| 20 | 0.000 | 0.001 | 100.000 |
| 21 | 0.000 | 0.000 | 100.000 |
| 22 | 0.000 | 0.000 | 100.000 |
| 23 | 0.000 | 0.000 | 100.000 |

**Supplementary Table 72. PCA Results.** Principal Component Analysis (PCA) shows that the first two components account for 75.02% of the total variance, with Component 1 explaining 47.39% and Component 2 explaining 27.63%. The first four components together capture 91.43% of the variance.

**SUBJECT 6. Contribution of Variables to the Dimensions (% Contribution)**

| Variables | Dim.1 | Dim.2 | Dim.3 | Dim.4 | Dim.5 |
|---|---|---|---|---|---|
| Delta Central | 2.898 | 2.093 | 17.373 | 0.154 | 0.506 |
| Theta Central | 3.291 | 0.561 | 16.727 | 0.186 | 6.037 |
| Alpha Central | 1.564 | 9.476 | 1.811 | 0.071 | 7.769 |
| Beta Central | 5.469 | 3.949 | 1.269 | 0.533 | 0.007 |
| Low-Gamma Central | 5.559 | 3.838 | 0.003 | 0.308 | 4.863 |
| Delta Frontal | 3.649 | 3.826 | 0.579 | 18.893 | 1.552 |
| Theta Frontal | 0.774 | 0.497 | 0.301 | 52.197 | 12.321 |
| Alpha Frontal | 1.657 | 10.343 | 0.323 | 1.144 | 2.522 |
| Beta Frontal | 6.711 | 0.902 | 0.120 | 5.057 | 1.809 |

| | | | | | |
|---|---|---|---|---|---|
| Low-Gamma Frontal | 5.651 | 3.187 | 0.011 | 0.034 | 7.052 |
| Delta Temporal | 5.205 | 2.050 | 6.205 | 2.720 | 0.012 |
| Theta Temporal | 4.334 | 0.186 | 16.054 | 0.580 | 0.862 |
| Alpha Temporal | 3.650 | 7.726 | 0.097 | 0.944 | 0.024 |
| Beta Temporal | 4.177 | 5.530 | 1.391 | 0.001 | 3.306 |
| Low-Gamma Temporal | 5.037 | 5.027 | 0.098 | 0.110 | 0.000 |
| Delta Parietal | 3.740 | 2.966 | 11.683 | 0.305 | 0.017 |
| Theta Parietal | 4.468 | 0.634 | 12.314 | 0.398 | 0.565 |
| Alpha Parietal | 1.666 | 11.077 | 0.677 | 0.001 | 0.616 |
| Beta Parietal | 4.414 | 5.692 | 0.379 | 0.003 | 0.060 |
| Low-Gamma Parietal | 5.162 | 4.308 | 0.000 | 0.346 | 3.614 |
| Delta Occipital | 5.402 | 1.882 | 0.118 | 3.022 | 5.740 |
| Theta Occipital | 5.498 | 0.638 | 7.846 | 0.039 | 0.073 |
| Alpha Occipital | 4.984 | 2.965 | 2.946 | 2.253 | 5.241 |
| Beta Occipital | 1.085 | 4.852 | 0.580 | 9.112 | 31.729 |
| Low-Gamma Occipital | 3.954 | 5.794 | 1.097 | 1.588 | 3.705 |

**Supplementary Table 73. Contribution of Variables to Dimensions.** The main contributors to Dim.1 are beta frontal (6.71%), low-gamma frontal (5.65%), and low-gamma central (5.56%). In Dim.2, the highest contributions come from alpha parietal (11.08%), alpha frontal (10.34%), and alpha central (9.48%).

**SUBJECT 6. Squared Cosine (Cos²) Values Indicating Variable Representation Across Principal Components**

| Variables | Dim.1 | Dim.2 | Dim.3 | Dim.4 | Dim.5 |
|---|---|---|---|---|---|
| Delta Central | 0.343 | 0.145 | 0.475 | 0.002 | 0.005 |
| Theta Central | 0.390 | 0.039 | 0.457 | 0.003 | 0.059 |
| Alpha Central | 0.185 | 0.655 | 0.050 | 0.001 | 0.076 |
| Beta Central | 0.648 | 0.273 | 0.035 | 0.007 | 0.000 |
| Low-Gamma Central | 0.659 | 0.265 | 0.000 | 0.004 | 0.048 |
| Delta Frontal | 0.432 | 0.264 | 0.016 | 0.258 | 0.015 |

| | | | | | |
|---|---|---|---|---|---|
| Theta Frontal | 0.092 | 0.034 | 0.008 | 0.714 | 0.121 |
| Alpha Frontal | 0.196 | 0.715 | 0.009 | 0.016 | 0.025 |
| Beta Frontal | 0.795 | 0.062 | 0.003 | 0.069 | 0.018 |
| Low-Gamma Frontal | 0.669 | 0.220 | 0.000 | 0.000 | 0.069 |
| Delta Temporal | 0.617 | 0.142 | 0.170 | 0.037 | 0.000 |
| Theta Temporal | 0.513 | 0.013 | 0.439 | 0.008 | 0.008 |
| Alpha Temporal | 0.432 | 0.534 | 0.003 | 0.013 | 0.000 |
| Beta Temporal | 0.495 | 0.382 | 0.038 | 0.000 | 0.033 |
| Low-Gamma Temporal | 0.597 | 0.347 | 0.003 | 0.002 | 0.000 |
| Delta Parietal | 0.443 | 0.205 | 0.320 | 0.004 | 0.000 |
| Theta Parietal | 0.529 | 0.044 | 0.337 | 0.005 | 0.006 |
| Alpha Parietal | 0.197 | 0.765 | 0.019 | 0.000 | 0.006 |
| Beta Parietal | 0.523 | 0.393 | 0.010 | 0.000 | 0.001 |
| Low-Gamma Parietal | 0.611 | 0.298 | 0.000 | 0.005 | 0.036 |
| Delta Occipital | 0.640 | 0.130 | 0.003 | 0.041 | 0.056 |
| Theta Occipital | 0.651 | 0.044 | 0.215 | 0.001 | 0.001 |
| Alpha Occipital | 0.590 | 0.205 | 0.081 | 0.031 | 0.052 |
| Beta Occipital | 0.129 | 0.335 | 0.016 | 0.125 | 0.312 |
| Low-Gamma Occipital | 0.468 | 0.400 | 0.030 | 0.022 | 0.036 |

**Supplementary Table 74. Squared Cosine ($Cos^2$) Values for Variable Representation Across Dimensions.** The variables best represented in Dim.1 are beta frontal ($Cos^2$ = 0.795), low-gamma central ($Cos^2$ = 0.659), and theta occipital ($Cos^2$ = 0.651). In Dim.2, the highest representations are found in alpha parietal ($Cos^2$ = 0.765), alpha frontal ($Cos^2$ = 0.715), and alpha central ($Cos^2$ = 0.655).

**SUBJECT 6. Contribution of Conditions to the Principal Components (% Contribution)**

| Condition | Dim.1 | Dim.2 | Dim.3 | Dim.4 | Dim.5 |
|---|---|---|---|---|---|
| FA | 10.700 | 63.827 | 9.544 | 11.100 | 14.933 |
| REM | 12.126 | 6.268 | 26.487 | 17.028 | 27.367 |
| S1 | 11.636 | 7.653 | 25.752 | 45.568 | 3.072 |

| | | | | | |
|---|---|---|---|---|---|
| Wakefulness | 65.539 | 22.252 | 38.216 | 26.304 | 54.628 |

**Supplementary Table 75. Contribution of Conditions to Principal Components.** Wakefulness contributes most to Dim.1 (65.54%) and Dim.5 (54.63%). False Awakening (FA) dominates Dim.2 (63.83%), while REM has the highest contribution to Dim.3 (26.49%) and Dim.5 (27.37%). S1 shows the strongest effect in Dim.4 (45.57%).

**SUBJECT 6. Squared Cosine ($Cos^2$) Values Indicating Condition Representation Across Principal Components.**

| Condition | Dim.1 | Dim.2 | Dim.3 | Dim.4 | Dim.5 |
|---|---|---|---|---|---|
| FA | 0.244 | 0.579 | 0.060 | 0.042 | 0.026 |
| REM | 0.391 | 0.106 | 0.184 | 0.101 | 0.094 |
| S1 | 0.446 | 0.150 | 0.203 | 0.128 | 0.009 |
| Wakefulness | 0.542 | 0.061 | 0.160 | 0.065 | 0.094 |

**Supplementary Table 76. Squared Cosine ($Cos^2$) Values for Condition Representation Across Dimensions.** FA is best represented in Dim.2 ($Cos^2 = 0.579$), while wakefulness shows the highest representation in Dim.1 ($Cos^2 = 0.542$). S1 has a moderate representation across multiple dimensions, with the highest in Dim.1 ($Cos^2 = 0.446$) and Dim.3 ($Cos^2 = 0.203$), while REM is most strongly represented in Dim.1 ($Cos^2 = 0.391$).

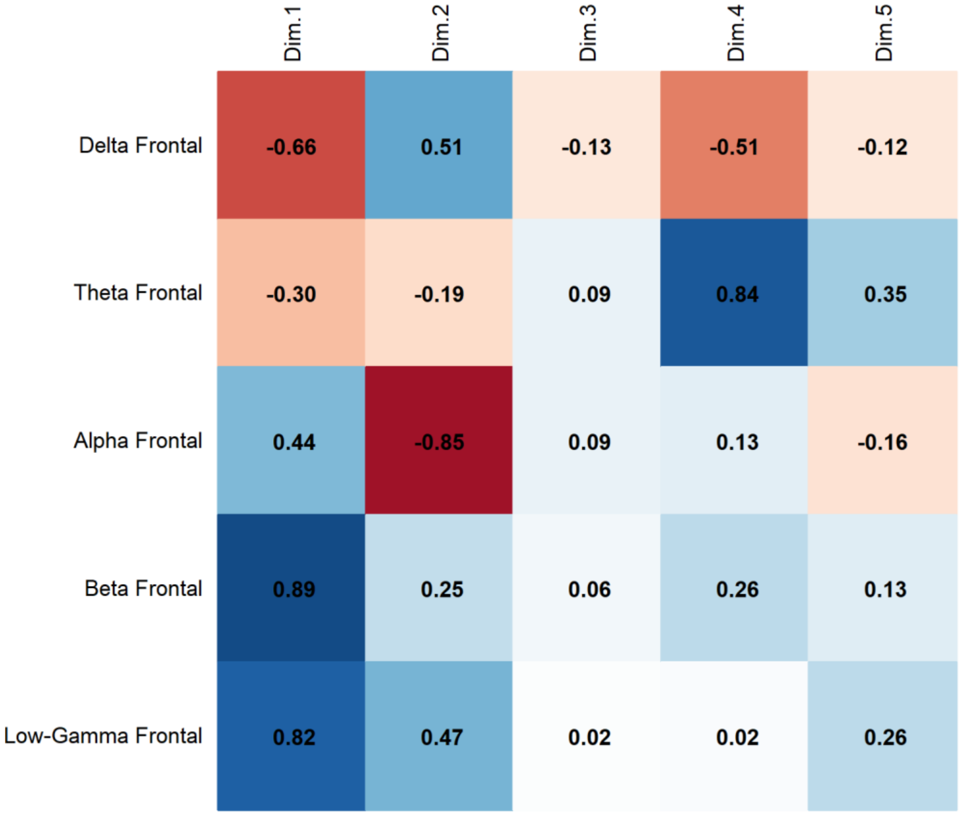


**Supplementary Fig. 35. False Awakening**, **PCA Results**. Correlation of Frontal Band Activity with Principal Components revealed that Beta activity presents a strong positive correlation with Dim.1 (0.89), while Alpha (-0.85) and Delta (-0.66) correlate negatively with Dim.1. Low-Gamma (0.82) is associated with Dim.1, and Theta (0.84) with Dim.4.

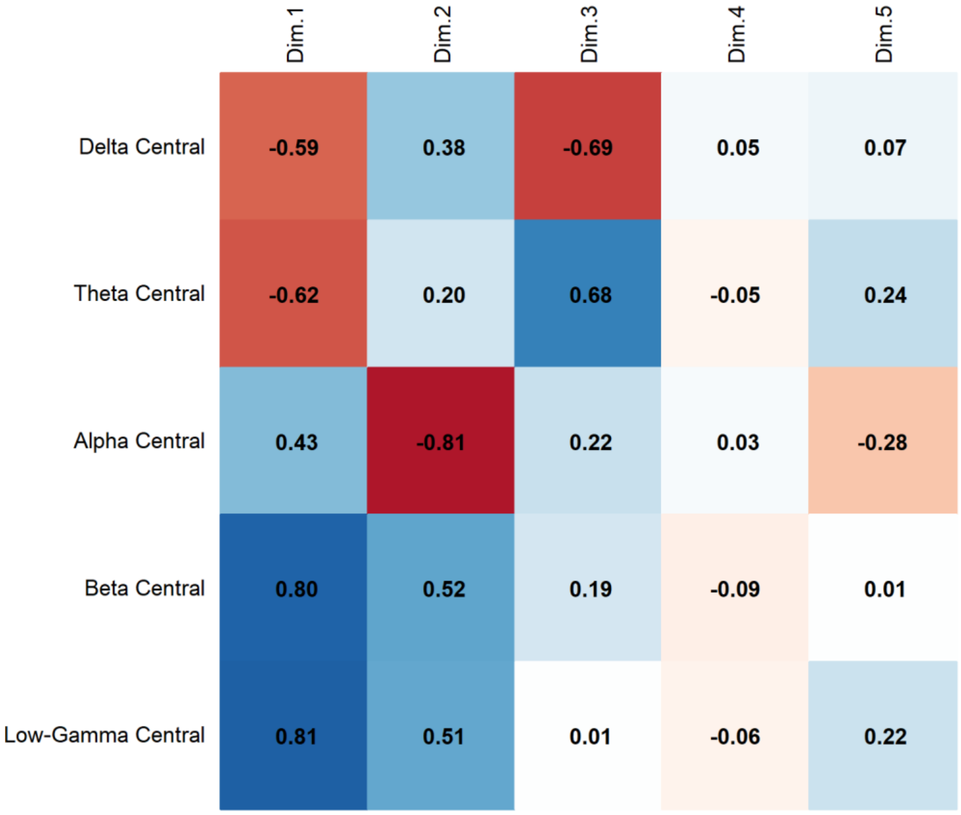


**Supplementary Fig. 36. False Awakening**, **PCA Results**. Correlation of Central Band Activity with Principal Components revealed that Beta activity presents a strong positive correlation with Dim.1 (0.80), while Alpha (-0.81) and Delta (-0.59) correlate negatively with Dim.1. Low-Gamma (0.81) is associated with Dim.1, and Theta (0.68) with Dim.3.

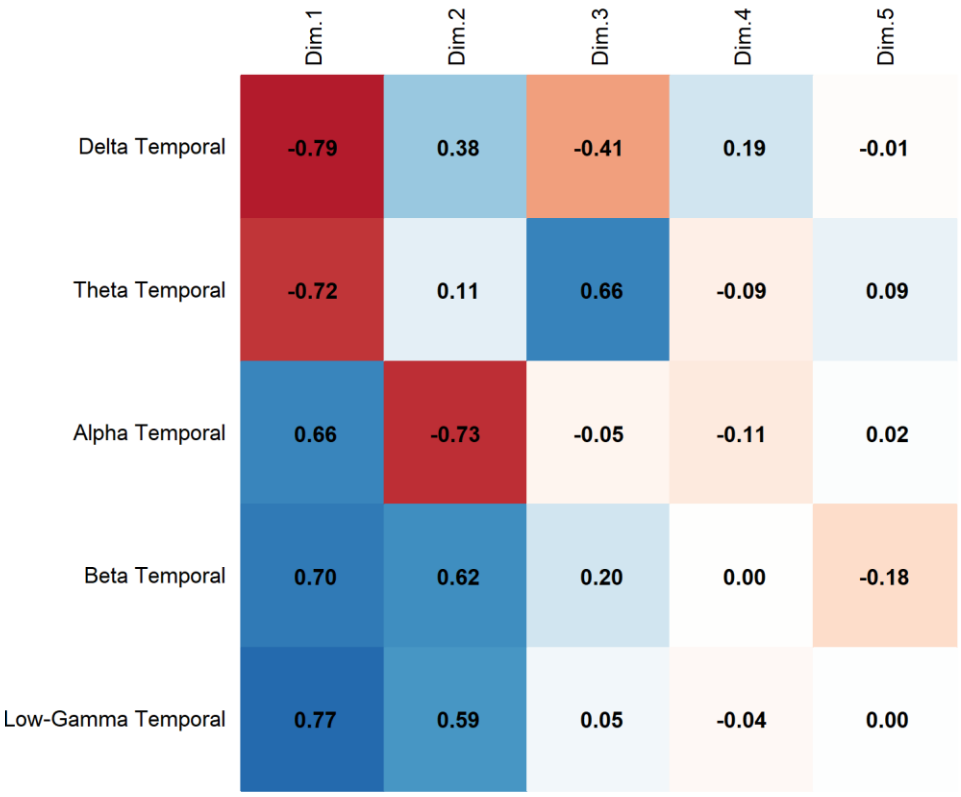


**Supplementary Fig. 37. False Awakening**, **PCA Results**. Correlation of Temporal Band Activity with Principal Components revealed that Beta activity presents a strong positive correlation with Dim.1 (0.70), while Delta (-0.79) and Theta (-0.72) correlate negatively with Dim.1. Alpha (0.66) is associated with Dim.1, and Low-Gamma (0.77) with Dim.1.

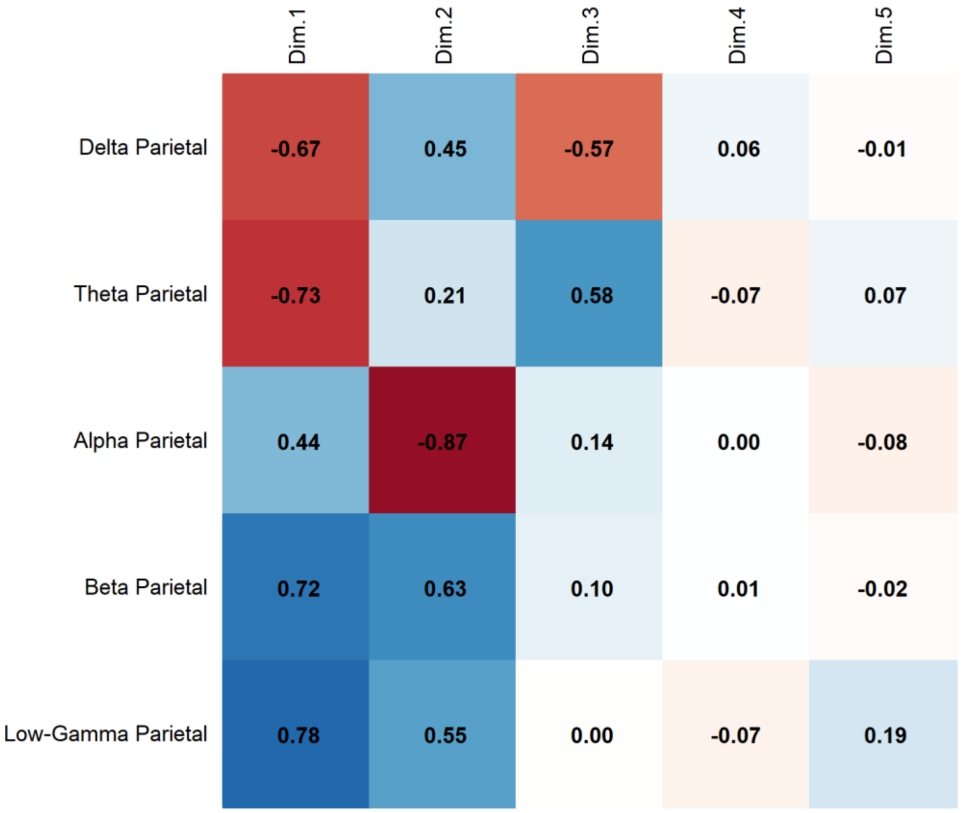


**Supplementary Fig. 38. False Awakening**, **PCA Results**. Correlation of Parietal Band Activity with Principal Components revealed that Beta activity presents a strong positive correlation with Dim.1 (0.72), while Alpha (-0.87) and Delta (-0.67) correlate negatively with Dim.1. Low-Gamma (0.78) is associated with Dim.1, and Theta (0.58) with Dim.3.

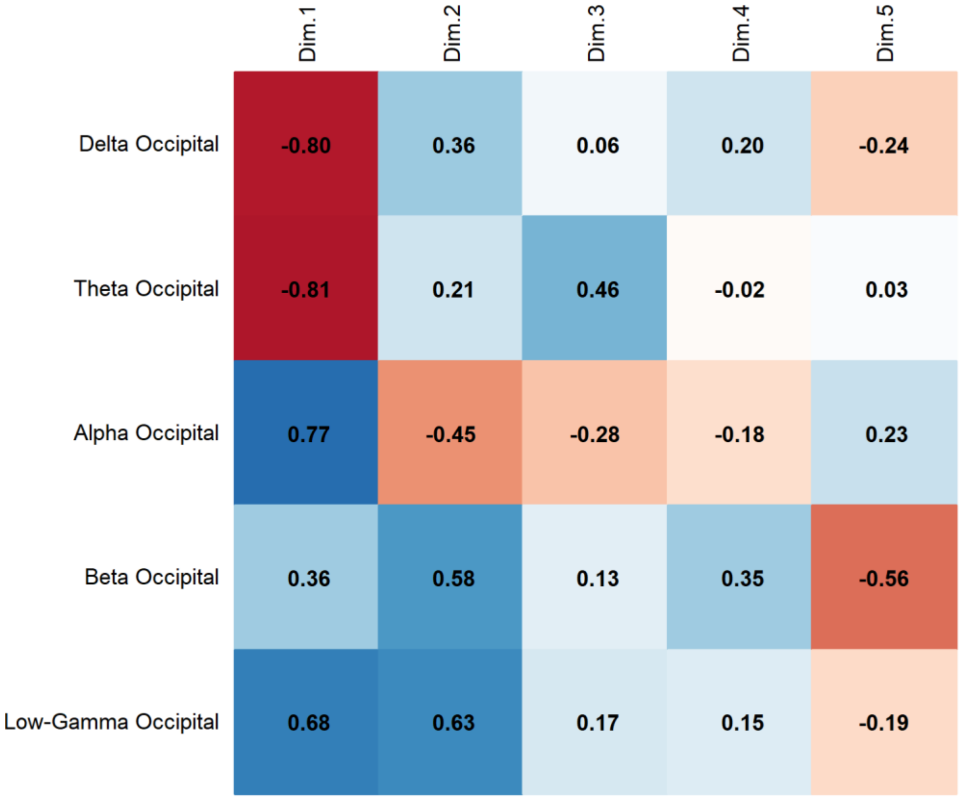


**Supplementary Fig. 39. False Awakening**, **PCA Results**. Correlation of Occipital Band Activity with Principal Components revealed that Alpha activity presents a strong positive correlation with Dim.1 (0.77), while Delta (-0.80) and Theta (-0.81) correlate negatively with Dim.1. Beta (0.58) is associated with Dim.2, and Low-Gamma (0.63) with Dim.2.

**SUBJECT 7. PCA Results**

| Component | Eigenvalue | Variance Explained (%) | Cumulative Variance (%) |
|---|---|---|---|
| 1 | 2.908 | 58.155 | 58.155 |
| 2 | 1.395 | 27.895 | 86.051 |
| 3 | 0.603 | 12.063 | 98.114 |
| 4 | 0.094 | 1.886 | 100.000 |
| 5 | 0.000 | 0.000 | 100.000 |

**Supplementary Table 77. PCA Results**. Principal Component Analysis (PCA) shows that the first two components account for 86.05% of the total variance, with Component 1 explaining 58.16% and Component 2 explaining 27.90%. The first three components together capture 98.11% of the variance.

**SUBJECT 7. Contribution of Variables to the Principal Components (% Contribution)**

| Variables | Dim.1 | Dim.2 | Dim.3 | Dim.4 |
|---|---|---|---|---|
| Delta Central | 24.54 | 5.40 | 34.36 | 3.93 |
| Theta Central | 20.68 | 2.40 | 60.30 | 1.57 |
| Alpha Central | 16.45 | 37.26 | 0.00 | 2.18 |
| Beta Central | 24.03 | 16.39 | 4.24 | 50.10 |
| Low-Gamma Central | 14.30 | 38.54 | 1.10 | 42.22 |

**Supplementary Table 78. Contribution of Variables to Principal Components.** The main contributors to Dim.1 are delta central (24.54%), beta central (24.03%), and theta central (20.68%). In Dim.2, the highest contributions come from low-gamma central (38.54%) and alpha central (37.26%).

**SUBJECT 7. Squared Cosine ($Cos^2$) Values Indicating Variable Representation Across Principal Components**

| Variables | Dim.1 | Dim.2 | Dim.3 | Dim.4 |
|---|---|---|---|---|
| Delta Central | 0.71 | 0.08 | 0.21 | 0.00 |
| Theta Central | 0.60 | 0.03 | 0.36 | 0.00 |
| Alpha Central | 0.48 | 0.52 | 0.00 | 0.00 |
| Beta Central | 0.70 | 0.23 | 0.03 | 0.05 |
| Low-Gamma Central | 0.42 | 0.54 | 0.01 | 0.04 |

**Supplementary Table 79. Squared Cosine ($Cos^2$) Values for Variable Representation Across Dimensions.** The variables best represented in Dim.1 are delta central ($Cos^2 = 0.71$) and beta central ($Cos^2 = 0.70$). In Dim.2, the highest representations are found in low-gamma central ($Cos^2 = 0.54$) and alpha central ($Cos^2 = 0.52$).

**SUBJECT 7. Contribution of Conditions to the Principal Components (% Contribution)**

| Condition | Dim.1 | Dim.2 | Dim.3 | Dim.4 |
|---|---|---|---|---|
| FA | 12.588 | 16.375 | 46.232 | 19.020 |
| REM | 24.021 | 0.638 | 23.884 | 17.751 |
| S1 | 20.528 | 48.345 | 27.073 | 28.127 |
| Wakefulness | 42.864 | 34.642 | 2.811 | 35.102 |

**Supplementary Table 80. Contribution of Conditions to Principal Components.** Wakefulness contributes most to Dim.1 (42.86%) and Dim.4 (35.10%). S1 dominates Dim.2 (48.35%), while FA has the highest contribution to Dim.3 (46.23%). REM shows a more balanced contribution across dimensions, with the highest in Dim.1 (24.02%) and Dim.3 (23.88%).

**SUBJECT 7. Squared Cosine (Cos²) Values Indicating Condition Representation Across Principal Components**

| Condition | Dim.1 | Dim.2 | Dim.3 | Dim.4 |
|---|---|---|---|---|
| FA | 0.224 | 0.578 | 0.186 | 0.012 |
| REM | 0.773 | 0.028 | 0.132 | 0.067 |
| S1 | 0.381 | 0.449 | 0.151 | 0.019 |
| Wakefulness | 0.700 | 0.271 | 0.017 | 0.012 |

**Supplementary Table 81. Squared Cosine ($Cos^2$) Values for Condition Representation Across Dimensions.** REM is best represented in Dim.1 ($Cos^2 = 0.773$), while FA has the highest representation in Dim.2 ($Cos^2 = 0.578$). S1 is moderately represented in both Dim.1 ($Cos^2 = 0.381$) and Dim.2 ($Cos^2 = 0.449$), whereas wakefulness shows strong representation in Dim.1 ($Cos^2 = 0.700$) and moderate representation in Dim.2 ($Cos^2 = 0.271$).

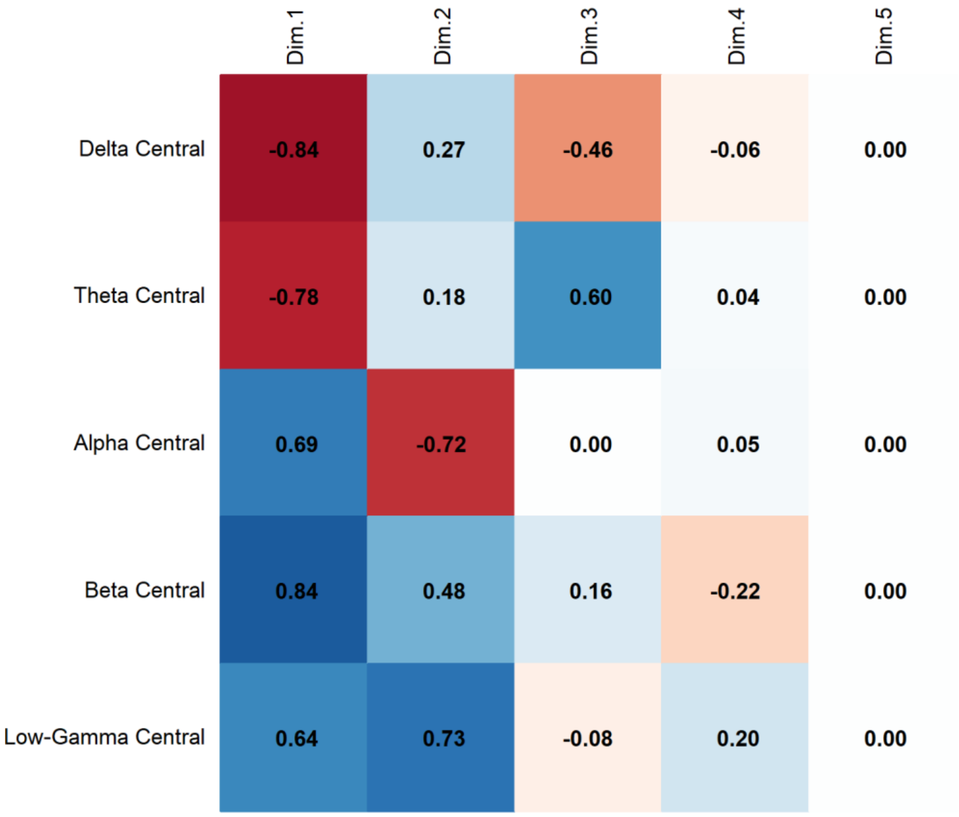


**Supplementary Fig. 40. False Awakening, PCA Results**. Correlation of Central Band Activity with Principal Components revealed that Beta activity presents a strong positive correlation with Dim.1 (0.84), while Delta (-0.84) and Theta (-0.78) correlate negatively with Dim.1. Low-Gamma (0.73) is associated with Dim.2, and Alpha (-0.72) with Dim.2.

### PERMANOVA

**SUBJECT 3 – FA. PERMANOVA Results.**

| | df | Sum of Squares | R² | F-value | p-value |
|---|---|---|---|---|---|
| Model | 3 | 6.1187 | 0.72949 | 17.978 | 1e-04 *** |
| Residual | 20 | 2.2690 | 0.27051 | | |
| Dispersion (Homogeneity) | 3 | 0.06725 | | 1.22 | 0.3283 |
| Dispersion Residuals | 20 | 0.36746 | | | |

**Supplementary Table 82. False Awakening. PERMANOVA Results.** The model explains 72.95% of the variance ($R^2$ = 0.72949) and is statistically significant ($p < 0.001$, Bonferroni-corrected). The homogeneity of dispersion test does not indicate significant differences in variance across groups ($p = 0.3283$), confirming that the assumption of homogeneity is met.

**SUBJECT 3 – FA. Post-hoc PERMANOVA Results**

| Comparison | Df | Sums Of Squares | F-value | R² | p-value | Adjusted p-value |
|---|---|---|---|---|---|---|
| False Awakening vs REM | 1 | 0.17159406 | 9.252 | 0.481 | 0.0052 | 0.031 |
| S1 vs False Awakening | 1 | 0.01500579 | 0.780 | 0.072 | 0.5451 | 1.000 |
| False Awakening vs Wakefulness | 1 | 0.32562040 | 15.087 | 0.601 | 0.0039 | 0.023 |
| REM vs Wakefulness | 1 | 0.76534440 | 67.174 | 0.870 | 0.0019 | 0.011 |
| S1 vs REM | 1 | 0.13925879 | 15.382 | 0.606 | 0.0017 | 0.010 |
| S1 vs Wakefulness | 1 | 0.30408821 | 25.153 | 0.716 | 0.0022 | 0.013 |

**Supplementary Table 83. False Awakening. Post-hoc PERMANOVA Results.** The model assesses differences between conditions with Bonferroni-adjusted p-values.

**SUBJECT 6. Comparisons between False Awakening and REM**

| Frequency Band | Brain Region | F-value | R² | p-value |
|---|---|---|---|---|
| Delta | Frontal | 31.740 | 0.760 | 0.002 |
| | Central | 17.302 | 0.634 | 0.007 |
| | Temporal | 21.796 | 0.685 | 0.005 |
| | Parietal | 17.693 | 0.639 | 0.002 |

| | | | | |
|---|---|---|---|---|
| | Occipital | 12.647 | 0.558 | 0.006 |
| Theta | Frontal | 12.448 | 0.555 | 0.010 |
| | Central | 3.356 | 0.251 | 0.096 |
| | Temporal | 7.215 | 0.419 | 0.022 |
| | Parietal | 1.480 | 0.129 | 0.272 |
| | Occipital | 1.344 | 0.119 | 0.286 |
| Alpha | Frontal | 9.137 | 0.477 | 0.018 |
| | Central | 3.329 | 0.250 | 0.032 |
| | Temporal | 3.498 | 0.259 | 0.040 |
| | Parietal | 3.289 | 0.248 | 0.059 |
| | Occipital | 2.675 | 0.211 | 0.138 |
| Beta | Frontal | 6.734 | 0.402 | 0.012 |
| | Central | 9.356 | 0.483 | 0.017 |
| | Temporal | 23.556 | 0.702 | 0.004 |
| | Parietal | 7.725 | 0.436 | 0.025 |
| | Occipital | 2.984 | 0.230 | 0.115 |
| Low-Gamma | Frontal | 2.325 | 0.189 | 0.123 |
| | Central | 0.835 | 0.077 | 0.379 |
| | Temporal | 0.197 | 0.019 | 0.665 |
| | Parietal | 0.173 | 0.017 | 0.685 |
| | Occipital | 0.000 | 0.000 | 0.990 |

**Supplementary Table 84. False Awakening. Comparisons between False Awakening and REM.** The model evaluates differences in spectral power across frequency bands and brain regions. Values are uncorrected for multiple comparisons.

**SUBJECT 3 – FA. Comparisons between False Awakening and Wakefulness**

| Frequency Band | Brain Region | F-value | $R^2$ | p-value |
|---|---|---|---|---|
| Delta | Frontal | 6.217 | 0.383 | 0.022 |
| | Central | 12.499 | 0.556 | 0.008 |

| | | | | |
|---|---|---|---|---|
| | Temporal | 10.113 | 0.503 | 0.013 |
| | Parietal | 14.764 | 0.596 | 0.009 |
| | Occipital | 11.788 | 0.541 | 0.011 |
| Theta | Frontal | 25.702 | 0.720 | 0.003 |
| | Central | 24.133 | 0.707 | 0.005 |
| | Temporal | 22.355 | 0.691 | 0.004 |
| | Parietal | 15.653 | 0.610 | 0.005 |
| | Occipital | 14.983 | 0.600 | 0.004 |
| Alpha | Frontal | 24.204 | 0.708 | 0.002 |
| | Central | 25.804 | 0.721 | 0.004 |
| | Temporal | 21.173 | 0.679 | 0.007 |
| | Parietal | 12.351 | 0.553 | 0.013 |
| | Occipital | 11.866 | 0.543 | 0.010 |
| Beta | Frontal | 1.303 | 0.115 | 0.283 |
| | Central | 2.778 | 0.217 | 0.130 |
| | Temporal | 0.637 | 0.060 | 0.482 |
| | Parietal | 2.452 | 0.197 | 0.164 |
| | Occipital | 0.877 | 0.081 | 0.378 |
| Low-Gamma | Frontal | 11.707 | 0.539 | 0.010 |
| | Central | 27.163 | 0.731 | 0.003 |
| | Temporal | 21.294 | 0.680 | 0.003 |
| | Parietal | 1.052 | 0.095 | 0.357 |
| | Occipital | 0.028 | 0.003 | 0.874 |

**Supplementary Table 85. False Awakening. Comparisons between False Awakening and Wakefulness.** The model evaluates differences in spectral power across frequency bands and brain regions. Values are uncorrected for multiple comparisons.

**SUBJECT 6. PERMANOVA Results**

| | df | Sum of Squares | R² | F | p-value |
|---|---|---|---|---|---|
| Model | 3 | 3.8165 | 0.48146 | 6.19 | 3e-04 *** |
| Residual | 20 | 4.1104 | 0.51854 | | |
| Dispersion (Homogeneity) | 3 | 0.090503 | | 1.844 | 0.1717 |
| Dispersion Residuals | 20 | 0.98162 | | | |

**Supplementary Table 86. PERMANOVA Results.** The model explains 48.15% of the variance ($R^2 = 0.48146$) and is statistically significant ($p < 0.001$, Bonferroni-corrected). The homogeneity of dispersion test does not indicate significant differences in variance across groups ($p = 0.1717$), confirming that the assumption of homogeneity is met.

**SUBJECT 6. Post-hoc PERMANOVA Results**

| Comparison | Df | Sum of Squares | F-value | R² | p-value | Adjusted p-value |
|---|---|---|---|---|---|---|
| False Awakening vs REM | 1 | 0.18171319 | 5.718 | 0.364 | 0.0379 | 0.227 |
| S1 vs False Awakening | 1 | 0.25660141 | 9.575 | 0.489 | 0.0060 | 0.036 |
| False Awakening vs Wakefulness | 1 | 0.12616930 | 3.069 | 0.235 | 0.0361 | 0.217 |
| REM vs Wakefulness | 1 | 0.25672498 | 10.051 | 0.501 | 0.0025 | 0.015 |
| S1 vs REM | 1 | 0.03855565 | 3.434 | 0.256 | 0.0228 | 0.137 |
| S1 vs Wakefulness | 1 | 0.29258366 | 14.229 | 0.587 | 0.0027 | 0.016 |

**Supplementary Table 87. Post-hoc PERMANOVA Results.** The model assesses differences between conditions with Bonferroni-adjusted p-values.

**SUBJECT 6. Comparisons between False Awakening and REM**

| Frequency Band | Brain Region | F-value | R² | p-value |
|---|---|---|---|---|
| Delta | Frontal | 7.465 | 0.427 | 0.031 |
| | Central | 3.620 | 0.266 | 0.090 |
| | Temporal | 5.133 | 0.339 | 0.054 |
| | Parietal | 3.509 | 0.260 | 0.087 |
| | Occipital | 3.322 | 0.249 | 0.090 |
| Theta | Frontal | 0.006 | 0.001 | 0.927 |
| | Central | 1.439 | 0.126 | 0.256 |

| | | | | |
|---|---|---|---|---|
| | Temporal | 1.019 | 0.093 | 0.345 |
| | Parietal | 4.651 | 0.317 | 0.073 |
| | Occipital | 2.738 | 0.215 | 0.124 |
| Alpha | Frontal | 6.002 | 0.375 | 0.039 |
| | Central | 5.971 | 0.374 | 0.031 |
| | Temporal | 7.178 | 0.418 | 0.027 |
| | Parietal | 8.432 | 0.457 | 0.023 |
| | Occipital | 7.137 | 0.416 | 0.029 |
| Beta | Frontal | 0.674 | 0.063 | 0.431 |
| | Central | 1.548 | 0.134 | 0.236 |
| | Temporal | 8.718 | 0.466 | 0.010 |
| | Parietal | 10.663 | 0.516 | 0.013 |
| | Occipital | 13.154 | 0.568 | 0.002 |
| Low-Gamma | Frontal | 3.278 | 0.247 | 0.056 |
| | Central | 14.717 | 0.595 | 0.002 |
| | Temporal | 21.751 | 0.685 | 0.002 |
| | Parietal | 18.922 | 0.654 | 0.003 |
| | Occipital | 15.698 | 0.611 | 0.006 |

**Supplementary Table 88. Comparisons between False Awakening and REM.** The model evaluates differences in spectral power across frequency bands and brain regions. Values are uncorrected for multiple comparisons.

**SUBJECT 6. Comparisons between S1 and False Awakening**

| Frequency Band | Brain Region | F-value | $R^2$ | p-value |
|---|---|---|---|---|
| Delta | Frontal | 15.958 | 0.615 | 0.009 |
| | Central | 0.425 | 0.041 | 0.527 |
| | Temporal | 0.746 | 0.069 | 0.417 |
| | Parietal | 0.930 | 0.085 | 0.351 |
| | Occipital | 4.715 | 0.320 | 0.064 |

| | | | | |
|---|---|---|---|---|
| Theta | Frontal | 4.393 | 0.305 | 0.069 |
| | Central | 21.514 | 0.683 | 0.002 |
| | Temporal | 27.527 | 0.734 | 0.002 |
| | Parietal | 53.226 | 0.842 | 0.002 |
| | Occipital | 17.402 | 0.635 | 0.002 |
| Alpha | Frontal | 8.574 | 0.462 | 0.027 |
| | Central | 7.078 | 0.414 | 0.042 |
| | Temporal | 8.617 | 0.463 | 0.016 |
| | Parietal | 9.720 | 0.493 | 0.018 |
| | Occipital | 13.095 | 0.567 | 0.005 |
| Beta | Frontal | 5.017 | 0.334 | 0.054 |
| | Central | 16.085 | 0.617 | 0.003 |
| | Temporal | 32.340 | 0.764 | 0.002 |
| | Parietal | 13.597 | 0.576 | 0.005 |
| | Occipital | 9.117 | 0.477 | 0.007 |
| Low-Gamma | Frontal | 12.450 | 0.555 | 0.011 |
| | Central | 76.299 | 0.884 | 0.002 |
| | Temporal | 59.285 | 0.856 | 0.003 |
| | Parietal | 52.153 | 0.839 | 0.003 |
| | Occipital | 42.541 | 0.810 | 0.003 |

**Supplementary Table 89. Comparisons between S1 and False Awakening.** The model evaluates differences in spectral power across frequency bands and brain regions. Values are uncorrected for multiple comparisons.

**SUBJECT 6. Comparisons between False Awakening and Wakefulness**

| Frequency Band | Brain Region | F-value | $R^2$ | p-value |
|---|---|---|---|---|
| Delta | Frontal | 0.253 | 0.025 | 0.626 |
| | Central | 0.003 | 0.000 | 0.985 |
| | Temporal | 0.739 | 0.069 | 0.417 |

| | | | | |
|---|---|---|---|---|
| | Parietal | 0.003 | 0.000 | 0.966 |
| | Occipital | 1.104 | 0.099 | 0.302 |
| Theta | Frontal | 5.063 | 0.336 | 0.055 |
| | Central | 1.056 | 0.096 | 0.307 |
| | Temporal | 6.532 | 0.395 | 0.026 |
| | Parietal | 0.934 | 0.085 | 0.351 |
| | Occipital | 7.391 | 0.425 | 0.029 |
| Alpha | Frontal | 2.427 | 0.195 | 0.157 |
| | Central | 2.060 | 0.171 | 0.186 |
| | Temporal | 0.210 | 0.021 | 0.660 |
| | Parietal | 2.724 | 0.214 | 0.129 |
| | Occipital | 0.753 | 0.070 | 0.393 |
| Beta | Frontal | 10.309 | 0.508 | 0.011 |
| | Central | 15.207 | 0.603 | 0.004 |
| | Temporal | 22.190 | 0.689 | 0.003 |
| | Parietal | 12.318 | 0.552 | 0.002 |
| | Occipital | 7.928 | 0.442 | 0.032 |
| Low-Gamma | Frontal | 6.506 | 0.394 | 0.002 |
| | Central | 7.005 | 0.412 | 0.002 |
| | Temporal | 12.007 | 0.546 | 0.002 |
| | Parietal | 5.269 | 0.345 | 0.002 |
| | Occipital | 8.720 | 0.466 | 0.005 |

**Supplementary Table 90. Comparisons between False Awakening and Wakefulness.** The model evaluates differences in spectral power across frequency bands and brain regions. Values are uncorrected for multiple comparisons.

**SUBJECT 7. PERMANOVA Results.**

| | df | Sum of Squares | $R^2$ | F | p-value |
|---|---|---|---|---|---|
| Model | 3 | 1.15418 | 0.60879 | 10.374 | 1e-04 *** |
| Residual | 20 | 0.74168 | 0.39121 | | |
| Dispersion (Homogeneity) | 3 | 0.04282 | | 0.8523 | 0.4817 |
| Dispersion Residuals | 20 | 0.33495 | | | |

**Supplementary Table 91. PERMANOVA Results.** The model explains 60.88% of the variance ($R^2 = 0.60879$) and is statistically significant ($p < 0.001$, Bonferroni-corrected). The homogeneity of dispersion test does not indicate significant differences in variance across groups ($p = 0.4817$), confirming that the assumption of homogeneity is met.

**SUBJECT 7. Post-hoc PERMANOVA Results**

| Comparison | Df | Sum of Squares | F-value | $R^2$ | p-value | Adjusted p-value |
|---|---|---|---|---|---|---|
| False Awakening vs REM | 1 | 0.0837143 | 3.282 | 0.247 | 0.0573 | 0.344 |
| S1 vs False Awakening | 1 | 0.1270971 | 3.036 | 0.233 | 0.0476 | 0.286 |
| False Awakening vs Wakefulness | 1 | 0.3749598 | 12.442 | 0.554 | 0.0033 | 0.020 |
| REM vs Wakefulness | 1 | 0.7399182 | 51.465 | 0.837 | 0.0025 | 0.015 |
| S1 vs REM | 1 | 0.1153989 | 4.421 | 0.307 | 0.0165 | 0.099 |
| S1 vs Wakefulness | 1 | 0.5074057 | 16.510 | 0.623 | 0.0019 | 0.011 |

**Supplementary Table 92. Post-hoc PERMANOVA Results.** The model assesses differences between conditions with Bonferroni-adjusted p-values.

**Subject 7. Comparisons between False Awakening and Wakefulness**

| Frequency Band | Brain Region | F-value | $R^2$ | p-value |
|---|---|---|---|---|
| Delta | Central | 10.738 | 0.518 | 0.002 |
| Theta | Central | 18.886 | 0.654 | 0.004 |
| Alpha | Central | 7.443 | 0.427 | 0.022 |
| Beta | Central | 11.599 | 0.537 | 0.002 |
| Low-Gamma | Central | 10.701 | 0.517 | 0.002 |

**Supplementary Table 93. Comparisons between False Awakening and Wakefulness.** The model evaluates differences in spectral power across frequency bands in the central region. Values are uncorrected for multiple comparisons.
